# Genotyping-by-sequencing illuminates high levels of divergence among sympatric forms of coregonines in the Laurentian Great Lakes

**DOI:** 10.1101/784355

**Authors:** Amanda S. Ackiss, Wesley A. Larson, Wendylee Stott

**Affiliations:** Wisconsin Cooperative Fishery Research Unit, College of Natural Resources, University of Wisconsin-Stevens Point, 800 Reserve St., Stevens Point, WI 54481; U.S. Geological Survey, Wisconsin Cooperative Fishery Research Unit, College of Natural Resources, University of Wisconsin-Stevens Point, 800 Reserve St., Stevens Point, WI 54481; U.S. Geological Survey, Great Lakes Science Center, 1451 Green Rd., Ann Arbor, MI 48105

**Keywords:** adaptive divergence, management units, genomic islands of divergence, species complex, RAD sequencing, coregonines, population genomics, hybridization

## Abstract

Effective resource management depends on our ability to partition diversity into biologically meaningful units. Recent evolutionary divergence, however, can often lead to ambiguity in morphological and genetic differentiation, complicating the delineation of valid conservation units. Such is the case with the “coregonine problem,” where recent post-glacial radiations of coregonines into lacustrine habitats resulted in the evolution of numerous species flocks, often with ambiguous taxonomy. The application of genomics methods is beginning to shed light on this problem and the evolutionary mechanisms underlying divergence in these ecologically and economically important fishes. Here, we used restriction site-associated DNA (RAD) sequencing to examine genetic diversity and differentiation among sympatric species in the *Coregonus artedi* complex in the Apostle Islands of Lake Superior, the largest lake in the Laurentian Great Lakes. Using 29,068 SNPs, we were not only able to clearly distinguish the three most common forms for the first time, but putative hybrids and potentially mis-identified specimens as well. Assignment rates to form with our RAD data were 93-100% with the only mis-assignments arising from putative F1 hybrids, an improvement from 62-77% using microsatellites. Estimates of pairwise differentiation (*F*_ST_: 0.045-0.056) were large given the detection of hybrids, suggesting that hybridization among forms may not be successful beyond the F1 state. We also used a newly built *C. artedi* linkage map to look for islands of adaptive genetic divergence among forms and found widespread differentiation across the genome, a pattern indicative of long-term drift, suggesting that these forms have been reproductively isolated for a substantial amount of time. The results of this study provide valuable information that can be applied to develop well-informed management strategies and stress the importance of re-evaluating conservation units with genomic tools to ensure they accurately reflect species diversity.

## Introduction

Defining conservation units is one of the most fundamental yet challenging aspects of resource management (Coates, Byrne, & Moritz, 2018). Partitioning species into units with substantial reproductive isolation provides managers with the ability to monitor and regulate independently evolving groups that may respond differently to harvest, disease, habitat alteration, or climate change (Allendorf & Luikart, 2007; Ryder, 1986). Over the past several decades, advancements in genetic analysis have provided scientists with powerful tools to estimate the amount of gene flow between species or populations to inform the creation of conservation units (Olsen et al., 2014; Palsbøll, Bérubé, & Allendorf, 2007; Palsbøll, Peery, & Bérubé, 2010; Schwartz, Luikart, & Waples, 2006). With the arrival of the genomics era, the power and accuracy to discern levels of reproductive isolation, inbreeding, and effective population size has vastly improved (Allendorf, Hohenlohe, & Luikart, 2010), and tools such as genome scans have revolutionized our ability to identify and understand adaptative genetic variation (Funk, McKay, Hohenlohe, & Allendorf, 2012; Waples & Lindley, 2018). Despite these advancements, delineating discrete conservation units can still be problematic. For example, taxonomic uncertainty can lead to confusion regarding species boundaries (Bickford et al., 2007; Hey, Waples, Arnold, Butlin, & Harrison, 2003), and observed phenotypic, spatial, temporal, or behavioral differences can be opposed by apparent genetic panmixia (Als et al., 2011; Hoey & Pinsky, 2018; Palm, Dannewitz, Prestegaard, & Wickström, 2009).

Perhaps no other group embodies the challenges of defining conservation units better than the coregonines. A subfamily of the Salmonidae, coregonines are comprised of three genera of freshwater and anadromous fishes distributed throughout cold water habitat in North America, Europe, and Asia. The most speciose genus, *Coregonus*, includes the ciscoes and whitefishes, which exhibit an extreme array of phenotype variability that is attributed to recent adaptive radiation into lacustrine habitat following glacial retreat during the Pleistocene epoch (Schluter, 1996). Often, distinct phenotypes can be found both in sympatry and allopatry which leads to difficulty in distinguishing a single, monophyletic origin of forms from parallel ecological speciation in individual lakes. Several coregonines exhibit sympatric dwarf and normal forms, including European whitefish *C. lavaretus*, North American whitefish *C. clupeaformis*, cisco *C. artedi,* and least cisco *C. sardinella* (Huitfeldt-Kaas, 1918; Mann & McCart, 1981; Shields, Guise, & Underhill, 1990; Vuorinen, Bodaly, Reist, Bernatchez, & Dodson, 1993). Empirical evidence of hybridization and introgression (Garside & Christie, 1962; Kahilainen et al., 2011; Lu, Basley, & Bernatchez, 2001) raises the question of how to manage forms when reproductive isolation is incomplete and has led some to suggest that the broad phenotypic variation observed in coregonines is a result of reticulate evolution (Svärdson, 1998; Turgeon & Bernatchez, 2003). This breadth of taxonomic ambiguity in the coregonines was first termed “the coregonid problem” by Svärdson (1949) and persists today as the more accurate “coregonine problem” (Eshenroder et al., 2016; Mee, Bernatchez, Reist, Rogers, & Taylor, 2015).

In recent decades phenotypic data have been combined with genetic analysis in an attempt to untangle the complicated relationships among coregonines. Taxonomic units in coregonine systematics have traditionally relied on morphological characteristics such as head and body shape, morphometrics, and meristics such as gill raker counts (Himberg, 1970; Koelz, 1929; Svärdson, 1979). Phylogeographic analyses with allozymes, restriction fragment length polymorphisms (RLFPs), mitochondrial DNA (mtDNA), and microsatellites have helped resolve evolutionary relationships among species and forms (Bernatchez & Dodson, 1994; Østbye, Bernatchez, Naesje, Himberg, & Hindar, 2005), and this growing body of research indicates that relying solely on phenotypic traits can be problematic for determining phylogenetic relationships when environmental plasticity occurs (e.g. Muir et al., 2013; Todd, 1998; Todd, Smith, & Cable, 1981). Amplified fragment length polymorphisms (AFLPs), mtDNA, and microsatellite analyses all show support for broad colonization of monophyletic lineages followed by the parallel ecological speciation of sympatric forms of European whitefish *C. lavaretus* (Hudson, 2011; Hudson, Lundsgaard-Hansen, Lucek, Vonlanthen, & Seehausen, 2017; Præbel et al., 2013). Recently, genomics methods have been applied to the coregonine problem in lake whitefish *C. clupeaformis*, and restriction site-associated DNA (RAD) sequencing, quantitative trait loci (QTL) analysis, and genome scans have provided valuable insight into the evolutionary mechanisms of speciation-with-gene-flow in dwarf and normal forms (Gagnaire, Normandeau, Pavey, & Bernatchez, 2013; Gagnaire, Pavey, Normandeau, & Bernatchez, 2013; Laporte et al., 2015; Rogers & Bernatchez, 2007).

The most extensive regional adaptive radiation within the *Coregonus* genus in North America occurred in the Laurentian Great Lakes, but detection of genetic differentiation or reproductive isolation among Great Lakes forms has been mostly unsuccessful. Rapid diversification of the shallow water cisco *C. artedi* (Eshenroder et al., 2016; Turgeon & Bernatchez, 2003) into newly available deepwater habitat following the Wisconsin Glacial Episode resulted in the evolution of at least eight distinct forms (Koelz, 1929; Scott & Crossman, 1998). Morphological differences occur across a variety of traits including body and head shape, lower jaw position, eye size, fin length, and gill raker counts (Koelz, 1929), though subtle variations among forms and between lakes can often make visually distinguishing them difficult without all possible forms present (Eshenroder et al., 2016; Turgeon et al., 2016). Different forms typically occur in specific depth ranges, and stable isotope analysis supports niche differentiation indicating that many of the Great Lakes forms occupy different trophic levels with observed changes in proportion of pelagic and benthic food sources (Schmidt, Harvey, & Vander Zanden, 2011; Schmidt, Vander Zanden, & Kitchell, 2009; Sierszen et al., 2014). All forms likely undergo seasonal spawning migrations, forming nearshore aggregations in November and December (Stockwell, Hrabik, Jensen, Yule, & Balge, 2010; Yule, Stockwell, Evrard, Cholwek, & Cullis, 2006; Yule, Addison, Evrard, Cullis, & Cholwek, 2009). However, very little is known about behavioral differences between forms during overlapping periods of spawning, maintaining the possibility that hybridization during spawning events could be preventing genetic divergence. Analyses using RFLPs, mtDNA, and microsatellites have resulted in estimates of low or no genetic differentiation among Great Lakes forms (Bernatchez, Colombani, & Dodson, 1991; Reed, Dorschner, Todd, & Phillips, 1998; Turgeon & Bernatchez, 2003; Turgeon et al., 2016) leaving the question of reproductive isolation - particularly among deepwater forms – unanswered.

Over the past century, anthropogenic impacts have greatly reduced the original diversity of the *C. artedi* species complex in the Great Lakes, underscoring the need for establishing well-informed conservation units in ciscoes. The introduction of invasive forage fish, overfishing, and habitat loss led to large decreases in cisco abundance and lake-wide extirpation to complete extinction of historically documented deepwater forms (Commission & Christie, 1973; Smith, 1968, 1970; Wells & McClain, 1973). Of the eight accepted forms originally described, only *C. artedi* and three deepwater forms – *C. hoyi*, *C. kiyi*, and *C. zenithicus* - are extant in the Great Lakes (referred to henceforth by specific epithet; Bailey and Smith, 1981; Todd and Smith, 1992). A fifth deepwater form, *C. nigripinnis,* has been reduced to nearby Lake Nipigon (Ontario, Canada), though an extant *nigripinnis*-like form is still periodically caught in Lake Superior (Eshenroder et al., 2016). Lake Superior’s peripheral location relative to both large human populations (and associated fishing pressure) and the canal construction that opened the Great Lakes to invasion from non-native species appears to have provided some protection from the impacts that extirpated cisco forms from the other four lakes (Koelz, 1926). Of the four lakes in which members of the *C. artedi* complex remain, Lake Superior is the only lake where all extant Great Lakes forms can still be regularly found (Eshenroder et al., 2016).

Recent evidence in the Great Lakes for declining abundance of invasive fish such as alewife and increasing abundance in cisco (Bronte et al. 2003, Mohr and Ebener 2005, Schaeffer et al. 2007) has led to growing interest in re-establishing lost populations. An understanding of the roles of heritable genetic differences and reproductive isolation in the establishment and persistence of remnant forms is vital for developing both informed conservation units and restoration strategies. The main goal of our study was to employ genomic methods to improve our understanding of genetic variation among these forms. Specifically, we (1) examined genetic differentiation and diversity among putative forms of cisco (2) compared the performance of SNPs and microsatellites in this system, and (3) leveraged a newly built cisco linkage map (Blumstein, 2019) to investigate adaptive divergence among forms. In order to remove the potentially confounding factor of distinguishing spatial genetic structure from form-based genetic structure, we focused on a single region in Lake Superior where multiple forms of cisco are found in sympatry, the Apostle Islands.

## Materials and Methods

Tissue samples preserved in >95% ethanol were collected for the three most common cisco forms– *artedi*, *hoyi*, and *kiyi* in the Apostle Islands (Fig. 1, Table 1). Samples of putative *artedi* were collected in November 2017 using top and bottom gillnet surveys off Madeline Island conducted by the Wisconsin Department of Natural Resources. Putative *artedi, hoyi* and *kiyi* were collected in July 2005 off Stockton Island by the U.S. Geological Survey Great Lakes Science Center (GLSC). In addition, we included a small number of available samples from two rare forms of cisco – *zenithicus* and the *nigripinnis-*like – from Minnesota, Michigan, and Ontario waters in Lake Superior collected by the GLSC in summer 2007, 2012-2015. All samples were identified to putative form using a suite of standardized morphological characteristics (Eshenroder et al., 2016). DNA was isolated from fin tissue samples with Qiagen DNeasy® Blood & Tissue Kits.

**Fig. 1.**
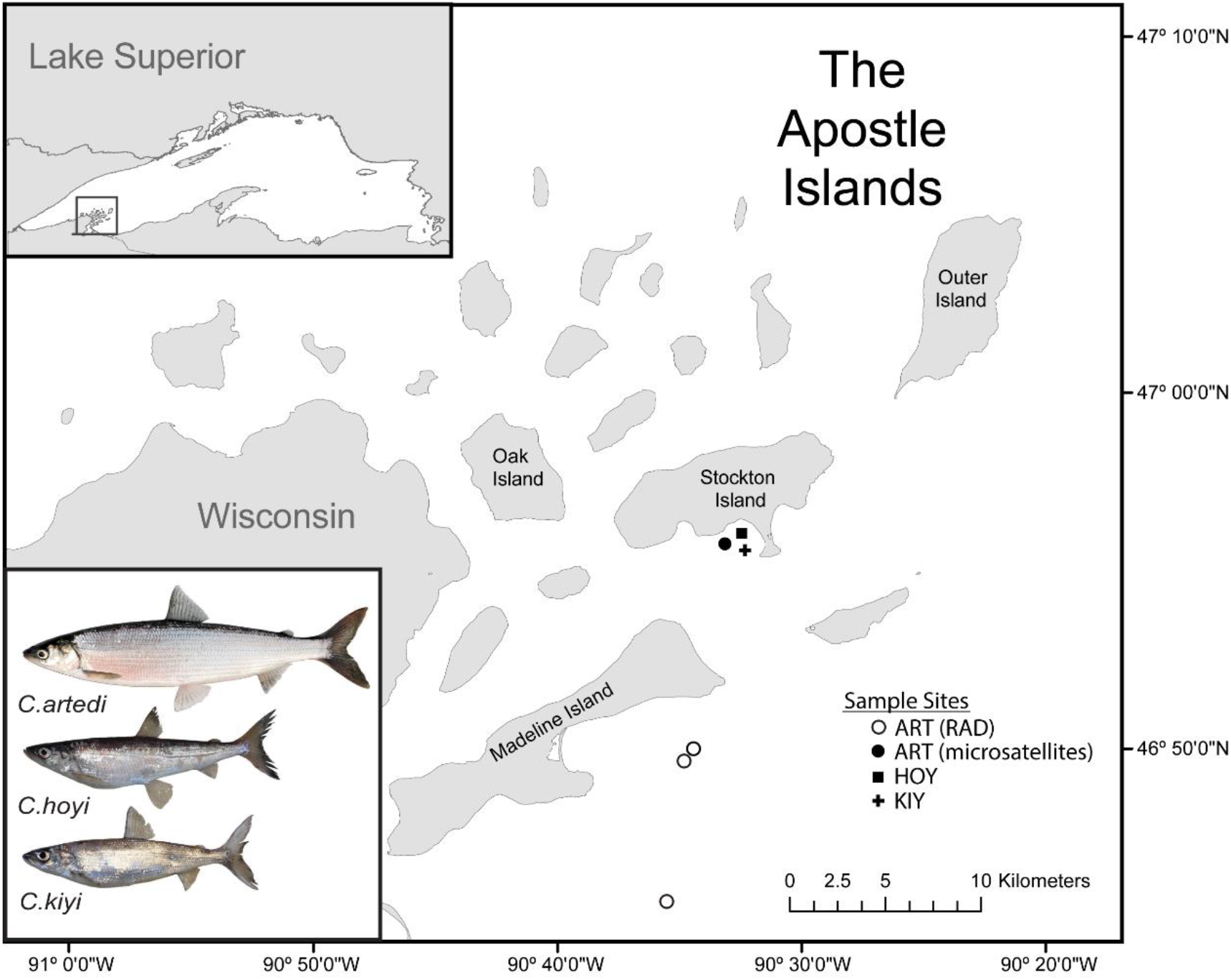
Map of sample sites in the Apostle Islands. INSET (upper right): Location of the Apostle Islands, gray box, within Lake Superior. INSET (lower right): Photos of the three most common cisco forms found in Lake Superior from Eshenroder et al. (2016, used with permission from the author).

**Table 1.**
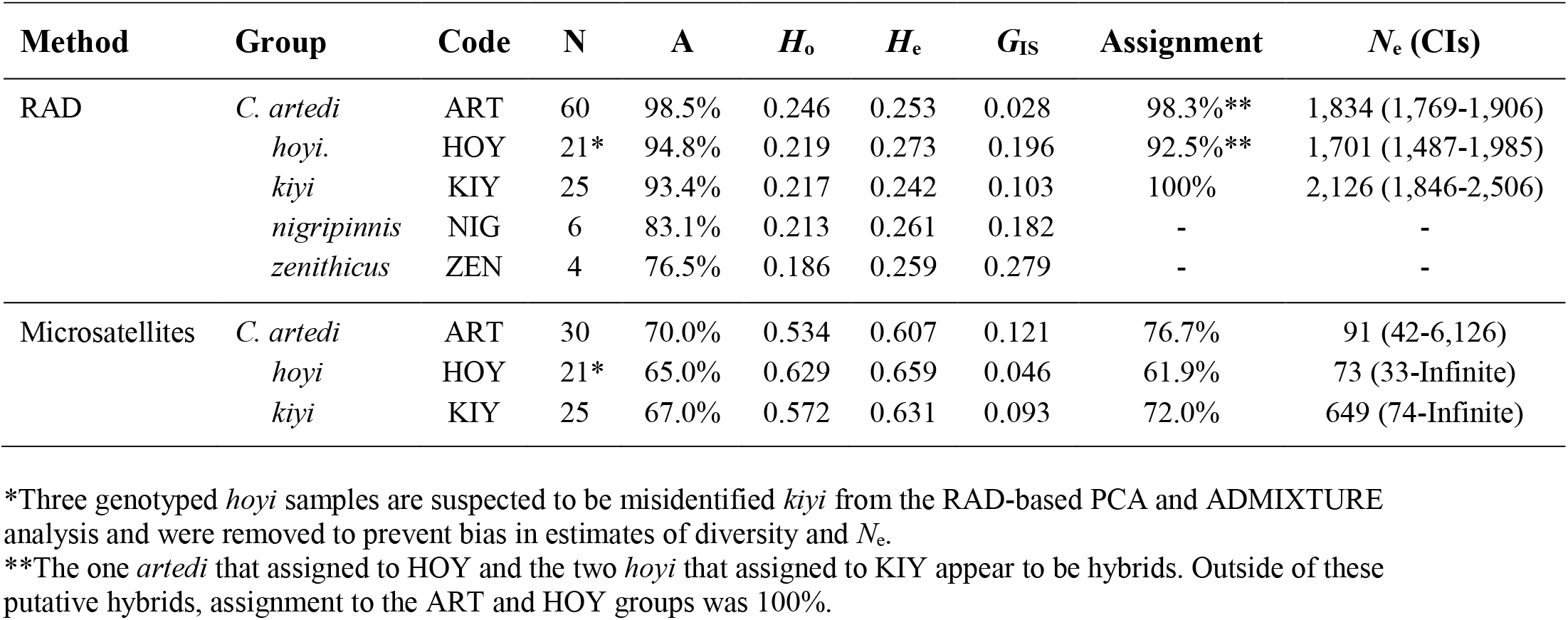
Sample statistics, diversity, and effective population size estimates. **N** is the number of individuals successfully genotyped, **A** is the percentage of total sampled alleles found in each group, ***H_o_/H_e_*** are observed/expected heterozygosity, ***G*_IS_** is the inbreeding coefficient, **Assignment** is the percentage of individuals that were correctly assigned to their population of origin in a leave-one-out test, and ***N_e_*** is effective population size calculated using the LDNE method and reported with 95% confidence intervals.

### RAD library prep and sequencing

Restriction site-associated DNA (RAD) libraries were prepared following the BestRAD protocol (Ali et al., 2016). Extracted DNA was quantified using a Quant-it™ PicoGreen® dsDNA Assay (Invitrogen, Waltham, MA) and normalized to a concentration of approximately 50 ng/µl for a 2 µl digestion reaction with the restriction enzyme *SbfI* followed by ligation with barcoded adaptors. Individually barcoded libraries were pooled into master libraries of 96 and fragmented to ∼300-500bp with 12-14 30s cycles in a Q500 sonicator (Qsonica, Newtown, CT). Fragmented DNA containing library adapters was bound to Dynabeads™ M-280 Streptavidin magnetic beads (Invitrogen) and washed with buffer to remove non-target fragments before an incubation step to release DNA from the beads. Following purification with AMPure XP beads (Beckman Coulter, Brea, CA) master libraries were input into the NEBNext® Ultra™ DNA Library Prep Kit for Illumina® at the End Prep step for ligation of master library barcodes, a 250-bp insert size-selection, and a 12-cycle PCR enrichment. Successful size-selection and enrichment were confirmed with visualization of products on a 2% agarose E-Gel (Invitrogen). Products underwent a final AMPure XP purification clean-up followed by quantification with a Qubit® 2.0 Fluorometer. All prepared libraries were sent to Novogene (Sacramento, CA) for sequencing on the Illumina NovaseqS4 platform.

### Read processing and SNP filtering

Raw sequences generated from RAD sequencing were processed in the software pipeline Stacks v2.3d (Catchen, Amores, Hohenlohe, Cresko, & Postlethwait, 2011; Catchen, Hohenlohe, Bassham, Amores, & Cresko, 2013). Sequences were demultiplexed by barcode, filtered for presence of the enzyme cut-site and quality, and trimmed in the subprogram *process_radtags* (parameter flags: -e *SbfI* -c -q -r -t 140 -- filter_illumina --bestrad). Filtered reads for each individual were aligned to create matching stacks with *ustacks* following guidelines suggested from empirical testing to avoid under- or over-merging loci from RAD datasets (Paris, Stevens, Catchen, & Johnston, 2017; parameter flags: --disable-gapped -m 3 -M 5 - H --max_locus_stacks 4 --model_type bounded --bound_high 0.05). A catalog of consensus loci built from 51 *artedi* sampled in Lake Huron used for the development of a cisco linkage map (Blumstein, 2019) was appended in *cstacks* with an additional 75 individuals in the *C. artedi* species complex from across the Great Lakes, including 19 fish used in the current study: five *hoyi* and four *kiyi* from the Apostle Islands and five each of *nigripinnis*-like and *zenithicus* from various locations across Lake Superior (parameter flags: -n 3 -p 6 –disable_gapped). Locus stacks for each individual were matched to the catalog using *sstacks* (parameter flag: --disable_gapped), data were oriented by locus in *tsv2bam*, and reads were aligned to loci and SNPs were called with *gstacks*. SNPs genotyped in greater than 30% of individuals (parameter flag: -r 0.3) were exported with the subprogram *populations* in genepop and variant call format (vcf) files.

Primary SNP filtering was performed with vcftools v0.1.15 (Danecek et al., 2011) and included (1) removing loci genotyped in fewer than 80% of individuals, (2) removing individuals missing more than 50% of loci, and (3) removing loci with a minor allele count less than 3. In addition, since all salmonids including members of the *C. artedi* species complex have experienced a recent genome duplication, putatively paralogous loci were identified with the program HDPlot (McKinney, Waples, Seeb, & Seeb, 2017), and any loci with heterozygosity greater than 0.55 or a read ratio deviation greater than 5 and less than –5 were removed. Finally, loci on the same RAD tag may be linked so only the SNP with the highest minor allele frequency on each tag was included in the final dataset. All file format conversions were performed using PGDSpider v2.1.1.5 (Lischer & Excoffier, 2011).

### Microsatellite amplification and genotyping

We used the methods described in Stott et al. (in press) to genotype 12 microsatellites developed for coregonines (Bernatchez, 1996; Patton, Gallaway, Fechhelm, & Cronin, 1997; Rogers, Marchand, & Bernatchez, 2004) and salmonids (Angers, Bernatchez, Angers, & Desgroseillers, 1995; Estoup, Presa, Krieg, Vaiman, & Guyomard, 1993) in *artedi*, *hoyi,* and *kiyi* from the Apostle Islands: *Bwf*1, *Bwf2*, *C2*- 157, *Cocl*23, *CoclLav*6, *CoclLav*27, *CoclLav*32, *CoclLav*72, *Sfo*8, *Sfo*23, *Str*-60. *Sfo8* consistently amplifies two genomic regions resulting in alleles sorting into upper (U: 215-281bp) and lower (L: 163-193bp) size ranges, therefore *Sfo8* is considered two loci. Fragment analysis was performed using a Genetic Analyzer 3.0 (Life Technologies), and genotypes were assigned at each locus using GeneMapper 3.7 (Life Technologies). We used Genepop v4 (Rousset, 2008) to conduct exact tests for deviations from Hardy–Weinberg and linkage equilibrium (α = 0.01). Three loci were removed for being out of Hardy-Weinberg Equilibrium in all three forms (*Bwf*1, *Cocl*23, *CoclLav*6), and no loci showed significant linkage disequilibrium.

### Genetic differentiation and diversity

To ensure that our putative form designations were appropriate, we used two approaches to assess genetic similarity among individuals. First, we conducted a principal component analysis (PCA) in the R package ‘adegenet’ (Jombart, 2008) for both the RAD and microsatellite datasets. Next, we estimated the number of ancestral populations, *K*, contributing to contemporary genetic clustering for the RAD dataset using the program ADMIXTURE v1.3 (Alexander, Novembre, & Lange, 2009). We tested *K* from 1-5 with ADMIXTURE’s cross-validation procedure and a k-fold of 10 (parameter flag: --cv=10) to examine support for each *K*. A *Q*-score of less than 70% was used to identify putative hybrids following similar thresholds applied in the literature (Kapfer, Sloss, Schuurman, Paloski, & Lorch, 2013; Marie, Bernatchez, & Garant, 2011; Weigel et al., 2018), and hybrid combinations were assigned to the two populations representing the largest *Q*-scores within the hybrid individual. ADMIXTURE analysis was only conducted on the RAD dataset because this program is not compatible with microsatellite data.

PCA and ADMIXTURE analysis with RAD genotypes revealed that three individuals originally identified as *hoyi* fell within the cluster of *kiyi* samples and, given the discrete clustering of *kiyi* and the strong possibility that these three *hoyi* samples were misidentified in the field, we removed these individuals from both RAD and microsatellite datasets for all further analyses to prevent bias in estimates of diversity, differentiation, and *N*_e_.

We calculated a variety of summary statistics for the groups comprised of the five forms (*artedi* -ART, *hoyi* – HOY, *kiyi* – KIY, *nigripinnis* – NIG, and *zenithicus* – ZEN) including percentage of total observed alleles, observed and expected heterozygosity, and an inbreeding coefficient (*G*_IS_). Summary statistics for each locus and populations were calculated using both microsatellite and RAD datasets in GenoDive v2.0b23 (Meirmans & Van Tienderen, 2004). Genetic differentiation among all forms was estimated across all loci in the RAD dataset with pairwise *F*_ST_ (Weir & Cockerham, 1984) in Genepop and tested using exact tests (Goudet et al., 1996; Raymond and Rousset, 1995; alpha = 0.01) in Arlequin (Excoffier & Lischer, 2010). We also calculated locus-specific overall and pairwise-*F*_ST_ values (Weir & Cockerham, 1984) in Genepop using a dataset that included the three forms with n>10 (ART, HOY, and KIY). To compare genetic differentiation between RAD and microsatellite datasets, we used GenoDive to estimate standardized pairwise genetic differentiation, *G*’_ST_ (Hedrick, 2005), which employs an additional correction for bias from sampling a limited number of populations (Meirmans & Hedrick, 2011).

The rate at which individuals were able to be assigned back to their form with both RAD and microsatellite datasets was tested using population assignment in GenoDive for all forms with n>10. Assignment was performed by calculating the home likelihood (L_h_) that an individual genotype is from a specific group given the allele frequencies (Paetkau, Calvert, Stirling, & Strobeck, 1995) using the leave-one-out method to avoid the bias from a target individual’s contribution to the allele frequencies of a source population. Zero frequencies were replaced with 0.005 and a significance threshold of alpha=0.002 was applied separately to each group over 1000 replicated datasets (c.f. Perreault-Payette et al., 2017).

Effective population size (*N*_e_) was estimated for all forms with n>10 using both RAD and microsatellite datasets with the bias-corrected linkage disequilibrium method (LDNE; Hill, 1981; Waples, 2006; Waples & Do, 2010) in the software package NeEstimator v2.1 (Do et al., 2014). We used a p-crit of 0.05 for the RAD dataset (Waples, Larson, & Waples, 2016) and 0.02 for the microsatellite dataset (Waples & Do, 2010). For the RAD dataset, only comparisons between sampled loci that were found on different linkage groups (LGs) of the cisco linkage map (Blumstein, 2019) were included to correct for physical linkage (Waples et al., 2016). *N*_e_ calculations using the linkage disequilibrium method can be biased slightly downward when individuals from multiple cohorts are included in the sample due to a slight Wahlund effect (7% downward bias on average; Waples, Antao, & Luikart, 2014). However, this small bias should not greatly affect the interpretation of the *N*_e_ results.

Finally, we used EASYPOPv2.0.1 (Balloux, 2001) to compare observed patterns of differentiation and hybridization with simulations encompassing a variety of demographic scenarios. Base parameters for all simulations were informed where possible by cisco life history traits and included random mating, same number of females and males per population (total n=2,000 per population based on our *N*_e_ estimates for the three main cisco forms), equal proportions of female and male migration, biallelic loci with free recombination, a mutation rate equivalent to that measured in human and fish nuclear genomes (µ=1.0 x 10^-8^; Bernardi & Lape, 2005; Conrad et al., 2011) and the KAM mutation model. First, since successful hybridization among forms will result in introgression and decreased differentiation, we modeled the impacts of hybridization on genetic differentiation with three scenarios: two populations with a starting level of differentiation equal to 1) approximately the level we observed amongst cisco forms with our RAD dataset (*F*_ST_ ≈0.05), 2) approximately twice the amount observed (*F*_ST_ ≈0.10), and 3) approximately four times the amount observed (*F*_ST_ ≈0.20). Preliminary levels of differentiation were achieved with low migration rates (m) set over 1,000 generations (m=0.001, m=0.0005, and m=0.0001, respectively) and followed by 5, 10, or 15 generations of one of three higher migration (i.e. hybridization) rates (m=0.01, 0.05, and 0.10) for every initial level of differentiation. Each unique parameter combination was run using a dataset of 1,000 loci and replicated 50 times. Estimates of *F*_ST_ for each replicate were generated in the R package ‘diveRsity’ (Keenan, McGinnity, Cross, Crozier, & Prodöhl, 2013) with a genepop file containing a subset of 100 randomly selected individuals per population (50 females/50 males). Second, we simulated migration between three populations representing our three common cisco forms (ART, HOY, KIY) in order to reconstruct datasets with similar characteristics to our empirical data and compare signatures of hybridization among them. Preliminary differentiation among the three populations (P1-P3) was set at *F*_ST_ ≈0.05 using the same method described above, and we allowed populations to hybridize for 2, 5, or 10 generations with m=0.05, which was equal to roughly the proportion of hybrids observed in our wild populations between ART-HOY and HOY-KIY. Since no putative ART-KIY hybrids were observed in our RAD dataset, we chose to implement a one-dimensional stepping stone model of migration (Kimura & Weiss, 1964) among simulated populations. Each unique parameter set for these simulations was run using 5,000 loci, a genepop file was output containing a subset of 100 randomly selected individuals per population (50 females/50 males), and ADMIXTURE was used to generate *Q*-scores for each individual. Hybrids were identified using the same *Q* < 0.70 threshold applied above.

### Differentiation across the genome

We examined genetic differentiation across the genome by pairing our data with the *artedi* linkage map constructed by Blumstein (2019). Catalog IDs were identical between the current study and Blumstein (2019) therefore no alignment step was needed to compare loci. To identify putative genomic islands of divergence that were highly differentiated from the rest of the genome, we used a Gaussian kernel smoothing technique (Gagnaire, Pavey, et al., 2013; Hohenlohe et al., 2010; Larson et al., 2017) that incorporated locus-specific differentiation and genomic position on the linkage map. A window size of 5 cM and a stepwise shift of 1 cM was used for this analysis, and values of genetic differentiation were weighted according to their window position as described by Gagnaire et al. (2013). Highly differentiated windows were identified by randomly sampling N loci from the genome (where N was the number of loci in the window) and comparing the average differentiation of those loci to the average differentiation of the loci in the window. This sampling routine was conducted 1,000 times for each window. If a window exceeded the 90^th^ percentile of the sampling distribution, the number of bootstrap replicates was increased to 10,000. Contiguous windows that contained at least two loci and exceed the 99^th^ percentile of the distribution after 10,000 bootstrap replicates were classified as putative islands of divergence. We investigated genomic differentiation using four locus-specific metrics: overall *F*_ST_ and pairwise *F*_ST_ between each of the three putative forms with n>10 (ART–HOY, ART–KIY, and HOY–KIY). We then plotted overall *F*_ST_ across the genome and constructed a bubble plot to visualize the number of significant windows on each LG for each comparison. We found over 50 significant windows for each comparison (see results). Conducting an in-depth investigation of all significant windows was not feasible, therefore we isolated in-depth analysis to one LG (Cart21) that contained the most significant windows in the dataset.

To investigate this highly differentiated LG, we aligned consensus sequences for all loci from Cart21 to chromosome Ssa05 in Atlantic salmon (*Salmo salar*), which is syntenic to Cart21 in *artedi* (Blumstein, 2019). Sequence was obtained from genome version ICSASG_v2 (Lien et al., 2016) and alignments were conducted with BLASTN. The best alignment for each locus was retained, and all alignments had e-values < 1e-58. We then visualized the relationship between recombination and physical distance using alignments to Ssa05 and information from the *artedi* linkage map to determine whether this highly differentiated region is characterized by lower recombination. We also obtained annotation information from the Atlantic salmon genome to determine whether genes of interest were co-located with areas of high divergence. Finally, we plotted the allele frequencies of the 10 SNPs on Cart21 with the highest overall *F*_ST_ to investigate whether these SNPs show consistent patterns of population structure.

## Results

### Sequencing and genotyping

A total of 137 individuals were RAD sequenced producing more than 455 million reads and an average of 3,346,457 reads per sample. After filtering, 119 individuals with representatives from all five putative cisco forms in Lake Superior were genotyped at 29,068 loci (Table 1). More than half of these loci (n=15,348 loci) were also placed on the linkage map. Since both RAD sequencing and microsatellite amplification were performed on the same *hoyi* and *kiyi* samples, microsatellite genotypes used in our analyses were restricted to the same individuals that successfully genotyped with our RAD loci. An additional 30 *artedi* from Stockton Island were genotyped at the 9 microsatellite loci. Paired-end assemblies for each locus are available from Blumstein (2019).

### Genetic differentiation and diversity

PCA showed a sharp contrast in resolution between marker sets with microsatellites producing one large cluster of overlapped forms across the first two principal components and the RAD dataset producing three major clusters primarily composed of ART, HOY, and KIY (Fig. 2). The ART cluster separates from HOY and KIY along the first principal component (PC), and HOY and KIY form discrete clusters along the second PC with three exceptions. Three individuals originally identified as *hoyi* fell within the KIY cluster (see methods and ADMIXTURE results below). Of the rare forms, *nigripinnis* (NIG) loosely grouped in the center of the PCA, with five of the six samples falling out between the ART, HOY, and KIY clusters (Fig. 2). The sixth sample fell within the ART cluster, and like the three *hoyi* in the KIY cluster, possibly represents a misidentified specimen. Unlike the NIG samples, which suggest the possibility for a distinct cluster with the addition of more specimens, the four ZEN samples closely associated with either the HOY cluster (n=1) or the KIY cluster (n=3). Low representation of both NIG and ZEN in our RAD dataset reduces our ability to draw strong conclusions based on these PCA results, so both groups were unaltered for estimates of diversity, inbreeding and differentiation and dropped for the remaining analyses.

**Fig. 2.**
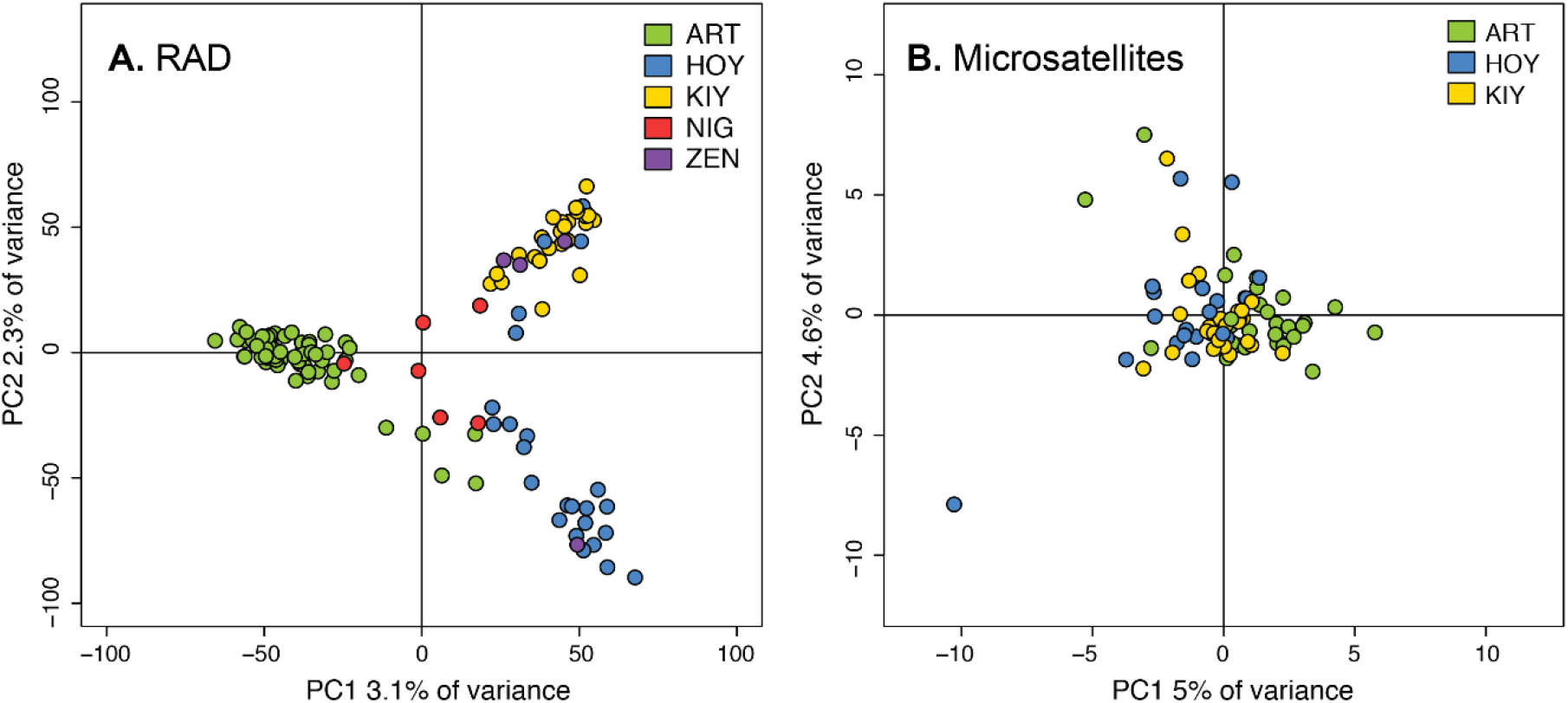
Principal components analysis with RAD (A) and microsatellite (B) data. The percentage of variance explained by each principal component (PC) is labeled on the x- and y-axes.

The most supported number of ancestral populations (*K*) estimated using the cross-validation procedure in ADMIXTURE was two (Fig. 3). Examining additional *K*s for significant sub-structuring among forms generated results that corroborated those from the PCA. When *K*=2, the ART cluster splits from HOY, KIY, and ZEN. Individuals in the NIG cluster exhibited mixed ancestry between the two major groups as seen on PC1 of the PCA. When *K*=3, the major genetic ancestries differentiate the ART, HOY, and KIY clusters as seen on PC2. The three putatively misidentified *hoyi* first noted in the PCA all had *Q* estimates of 100% for the KIY cluster and were removed from further analyses. Additional *K*s did not differentiate either the NIG or ZEN cluster but begin to differentiate small subsets of individuals within groups, which was likely statistical noise.

**Fig. 3.**
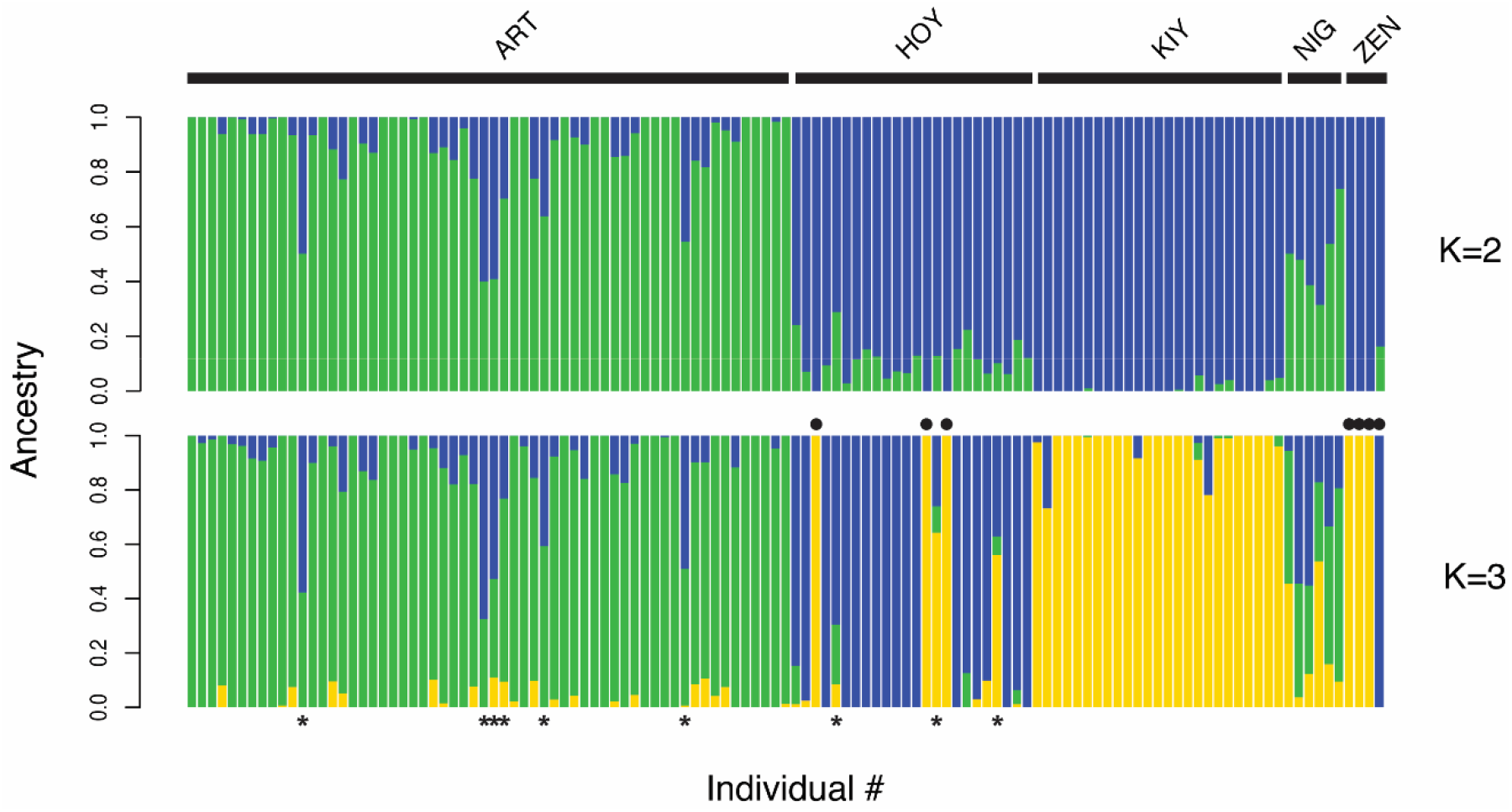
Genetic lineages in Apostle Island ciscoes estimated with ADMIXTURE. Each vertical bar represents a single individual and is colored by the proportion of ancestry (*Q*) assigned to each genetic lineage (*K*). In the *K*=3 plot (bottom), filled circles represent putatively misidentified individuals within forms (with *Q*-scores of 100% to other forms) and asterisks represent putative hybrids based on a *Q*-score threshold of less than 70%.

Observed and expected heterozygosity ranged from 0.186-0.246 and 0.242-0.273 (respectively) in the RAD dataset and 0.534-0.629 and 0.607-0.659 (respectively) in the microsatellite dataset (Table 1). Inbreeding coefficients were not substantially different from zero in both datasets. The largest *G*_IS_ was measured in ZEN from only four samples (0.279) and the rest were between −0.028-0.196. All estimates of pairwise genetic differentiation among forms with n>10 (ART, HOY, KIY) with the RAD dataset were significant (Table 2). The magnitude of genetic differentiation followed similar trends observed in the PCA, with ART being slightly more differentiated from deepwater forms (*F*_ST_=0.049-0.056). This pattern remained the same with a standardized measure of genetic differentiation in the RAD and microsatellite datasets (Table 3). All pairwise comparisons in both datasets were significant with higher overall values of *G*’_ST_ generated using microsatellites (0.110-0.122) than SNPs (0.060-0.75). See Tables S1 and S2 for locus-specific summary statistics.

**Table 2.** Pairwise differentiation among putative forms calculated with the RAD dataset. *F*_ST_ values on the lower diagonal. Significant values are in bold.

| Group | ART | HOY | KIY | NIG | ZEN |
| --- | --- | --- | --- | --- | --- |
| ART | - |  |  |  |  |
| HOY | **0.049** | - |  |  |  |
| KIY | **0.056** | **0.045** | - |  |  |
| NIG | 0.019 | **0.013** | **0.034** | - |  |
| ZEN | **0.061** | **0.017** | 0.020 | 0.014 | - |

**Table 3:** Standardized pairwise genetic differentiation, *G*’_ST_, (Hedrick, 2005) for SNP and microsatellite datasets. Values measured using the SNP dataset are in the lower diagonal, and values measured using the microsatellite dataset are in the upper diagonal.

| Group | ART | HOY | KIY |
| --- | --- | --- | --- |
| ART | - | 0.122 | 0.113 |
| HOY | 0.064 | - | 0.110 |
| KIY | 0.075 | 0.060 | - |

Population self-assignment rate using the microsatellite dataset ranged from 61.9-76.7% with *artedi* exhibiting the highest likelihood of being assigned back to the ART group (Table 1). Assignment rate with the RAD dataset was 100% for the KIY group and 98.3% and 92.5% for the ART and HOY groups (respectively), a result of one putative *artedi* assigning to HOY and two putative *hoyi* individuals assigning to KIY. In the ADMIXTURE analysis, these three individuals exhibited relatively high *Q* estimates for ancestry to the populations to which they were assigned (*artedi, Q*_HOY_ = 67.4%; *hoyi, Q*_KIY_= 64.2% and 56.2%). In the PCA generated from the same data, these individuals were oriented between the main clusters. In the microsatellite dataset with the same *hoyi* and *kiyi* samples, neither of the two potentially misclassified *hoyi* assigned to the HOY group, with one being assigned to ART and one to KIY. Assignment scores are reported in supplementary tables S3-S5.

Estimates of *N*_e_ with the microsatellite dataset ranged from 73 in HOY to 659 in KIY (Table 1). Only the estimate for ART produced both upper and lower bound confidence intervals (*N*_e_: 91, CI: 42-6,126), whereas the estimates for HOY and KIY produced confidence intervals with an ‘*infinite’* upper bound. An ‘*infinite’* upper bound is typically an indication that the data are not powerful enough to produce an accurate estimate of *N*_e_ given the sample size, population size, and/or marker resolution (Do et al. 2014; Waples & Do 2010). For the RAD dataset, estimates of *N*_e_ were generated with loci that were placed on the linkage map and ranged from 1,701-2,126 with confidence intervals within 10% of these values.

We documented a relatively high level of hybridization in our empirical dataset, with ART-HOY hybrids comprising 8.6% of the total number of sampled *artedi* and *hoyi*, HOY-KIY hybrids comprising 4.3% of sampled *hoyi* and *kiyi*, and ART-KIY hybrids comprising 0% (Fig. 3, Table S6). Additionally, most of these appear to be F1 hybrids with similar contributions from two genetic groups (Fig. 3). The fact that we observed frequent hybridization coupled with relatively high genetic differentiation was puzzling and prompted us to conduct two types of simulations to investigate how hybridization (i.e. migration) can influence genetic differentiation. Simulated hybridization over a 15-generation period resulted in declines in genetic differentiation among populations for all tested levels of migration (Fig. 4). When the migration rate was set to 5 or 10%, levels of differentiation more than halved after only five generations for all initial levels of *F*_ST_. A migration rate of 1% resulted in steadily declining genetic differentiation but only began to approach a halved *F*_ST_ after 15 generations.

**Fig. 4.**
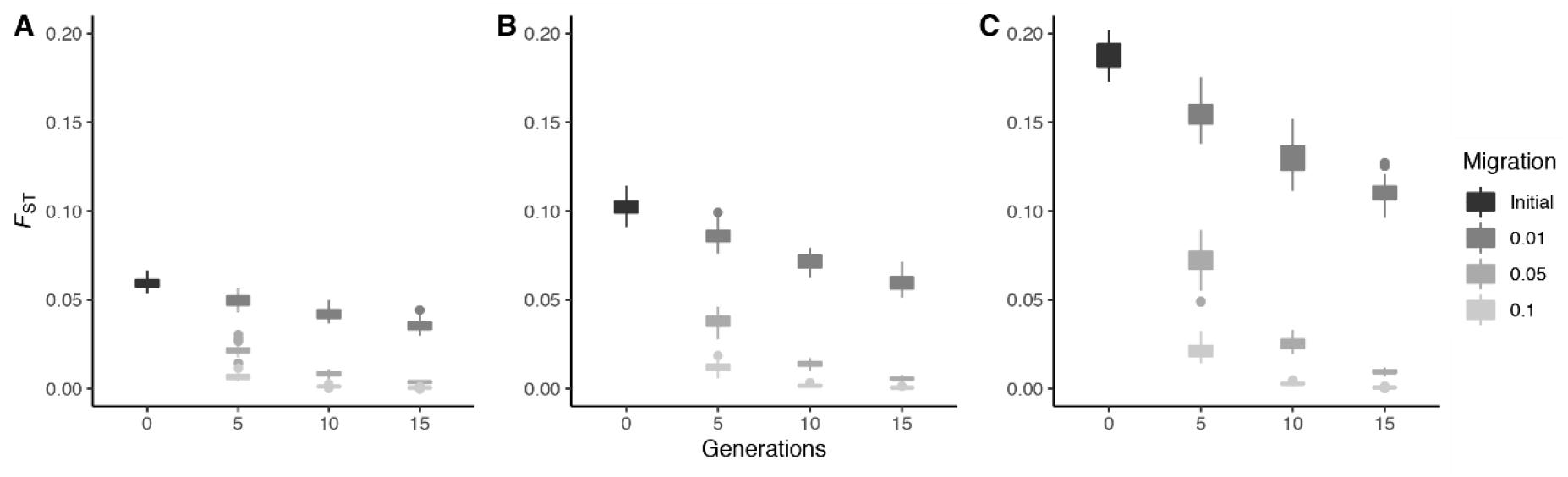
Simulated impacts of differing rates of hybridization under three different levels of preliminary differentiation. Each EASYPOP simulation was run using 1000 biallelic loci in two randomly mating populations each comprised of 1000 females and 1000 males with 50 replicates of each unique set of parameters for three starting levels of differentiation: A) a migration rate of 0.001 over 1000 generations resulting in an *F*_ST_≈0.05 followed by 5, 10, or 15 generations of hybridization at 1, 5, and 10% per generation, B) a migration rate of 0.0005 over 1000 generations resulting in an *F*_ST_≈0.10 followed by 5, 10, or 15 generations of hybridization at 1, 5, and 10% per generation, and C) a migration rate of 0.0001 over 1000 generations resulting in an *F*_ST_≈0.20 followed by 5, 10, or 15 generations of hybridization at 1, 5, and 10% per generation.

**Fig. 5.**
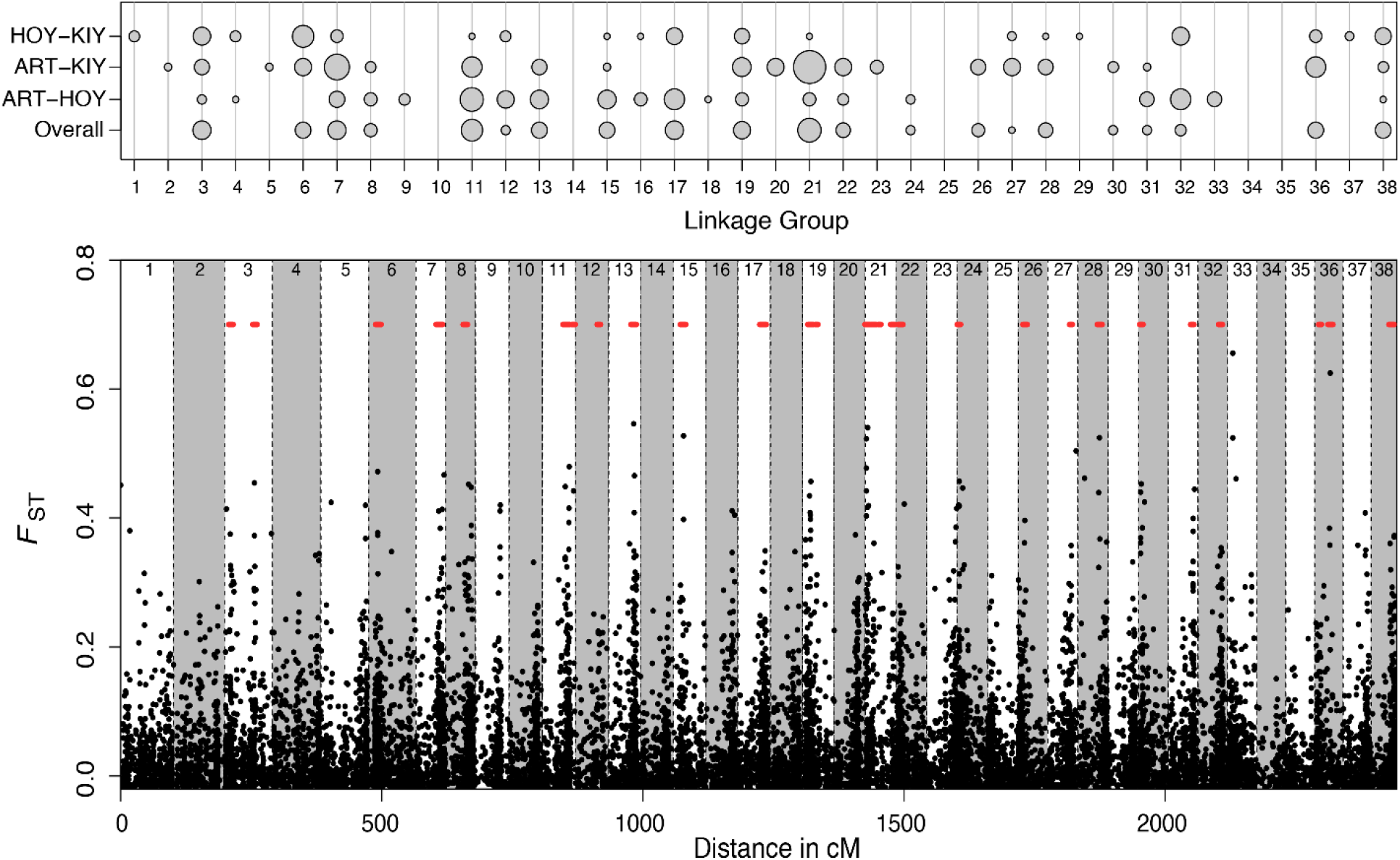
Genetic differentiation across the genome visualized with a bubble plot (top) and plot with the overall *F*_ST_ of each marker (bottom). The size of each bubble in the bubble plot represents the number of genomic windows that were significantly differentiated from the rest of the genome according to kernel smoothing analysis for each form comparison. The ‘overall’ designation is overall *F*_ST_ across the dataset. Each black dot in the graph of differentiation across the genome represents a marker, and red lines denote significantly differentiated windows. Linkage groups are separated by dashed lines. Form abbreviations are in Table 1. See Figs. S2-S5 for visualizations of genetic differentiation for each chromosome and form comparison.

Simulations of stepping stone migration between three populations and subsequent ADMIXTURE analysis resulted in an overall pattern very similar to that observed in our empirical data (Figs 3, S1; Table S6). The major goal of these simulations was to determine whether individuals that we observed in our empirical data with *Q*-scores between 0.1-0.2 for alternative forms were advanced hybrid backcrosses or whether these observations were statistical noise. Results from the simulations with 2 generations of migration (G2), where only F1 migrants are possible, demonstrated that these 0.1-0.2 *Q*-scores are likely statistical noise, as they were present in this simulation (Fig S1). Simulations with 5 (G5) and 10 (G10) generations of migration illustrate that advanced backcrosses will likely have *Q*-scores for alternate forms of at least 0.2. These simulations therefore support our hypothesis that most of the hybrids observed in the empirical data are F1 crosses.

### Differentiation across the genome

Genetic differentiation across the genome was generally high, with many loci displaying overall *F*_ST_ values > 0.2 (Figs 5, S2-S5). This differentiation was also not localized to a few LGs, as every LG had at least one locus with *F*_ST_ > 0.2. Kernel smoothing analysis to investigate putative islands of divergence revealed 389 genomic windows that displayed significantly elevated differentiation compared to the rest of the genome (Figs 5, S2-S5). The number of significant windows was highest for the overall *F*_ST_ comparison (115), followed by the ART-HOY comparison (101), ART-KIY comparison (98), and HOY-KIY comparison (75). Significantly differentiated windows were found on all but five LGs (average per LG=10, SD=10, range=0-36). LG Cart21 displayed the highest number of differentiated windows, prompting us to conduct an in-depth investigation of this LG to investigate patterns and potential drivers of divergence (Fig. 6).

**Fig. 6.**
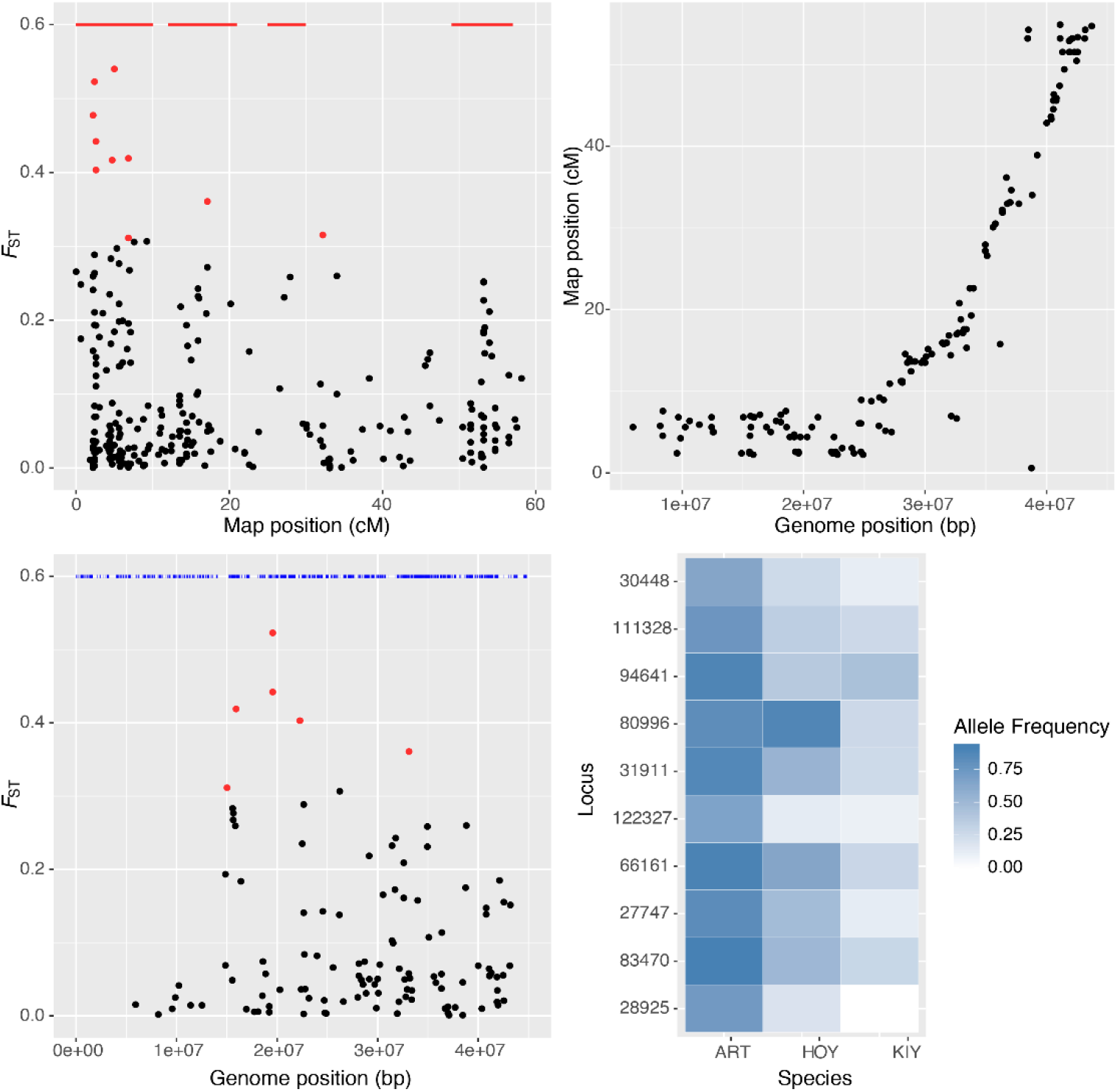
Investigation of genetic differentiation on linkage group Cart21, the linkage group with the most significantly differentiated windows. (a) Genetic differentiation (overall *F*_ST_) at 351 loci that were placed on Cart21 in the cisco linkage map. The top ten loci with the highest *F*_ST_ are colored red. Red lines denote significantly differentiated windows. (b) Recombination distance on cisco linkage group Cart21 (y-axis) versus physical distance on Atlantic salmon chromosome Ssa05 (x-axis). (c) Genetic differentiation (overall *F*_ST_) at 152 loci from Cart21 that successfully aligned to Ssa05, the syntenic chromosome in Atlantic salmon. Six of the top ten loci from panel a aligned to Ssa05 and are colored red. Blue lines indicate the position of genes on Ssa05. (d) Allele frequencies of the loci with the highest *F*_ST_ from Cart21. Loci are ordered from highest *F*_ST_ (bottom) to lowest. Form abbreviations are found in Table 1.

Cart21 contained 36 significantly differentiated windows; with the largest number of windows found for the ART-KIY comparison (18), followed by the overall *F*_ST_ comparison (13), ART-HOY comparison (4), and HOY-KIY comparison (1). We were able to place 351 loci on Cart21, and 12 of these loci displayed overall *F*_ST_ values > 0.3 (Fig. 6a). The largest cluster of high-*F*_ST_ loci was found between 0-10 cM on the linkage map. This region appears to be characterized by relatively low recombination, as loci found in the first 10 cM of Cart21 span about 25 megabases of the Atlantic salmon genome (Fig. 6b). Alignments to the Atlantic salmon genome were possible for 151 loci on Cart21, and these alignments revealed that the highest *F*_ST_ loci were found between positions 15 million and 25 million on Ssa05 (Fig. 6c). Some of these loci were found in genes with functions that include cell signaling and membrane transport. However, there are over 2,000 genes on Ssa05, making it difficult to reach any robust conclusions about the functional significance of our loci. Allele frequencies at the high-*F*_ST_ were generally the most diverged between ART and the other two forms, KIY in particular (Fig. 6d).

## Discussion

The resolution of genetic structure in recently diverged species complexes has proved challenging with traditional genetics methods, prompting the reevaluation with genomics of a taxonomically uncertain species complex in the Laurentian Great Lakes. Using 29,068 SNPs to examine differentiation and diversity of sympatric coregonines in the Apostle Islands of Lake Superior, we were able to unambiguously assign individuals to the three major forms as well as identify putative F1 hybrids and mis-identified individuals. Despite a century of anthropogenic impacts in the Great Lakes that has seen the extirpation and extinction of historically documented forms in the *C. artedi* species complex, estimates of *N*_e_ and diversity in Apostle Island populations do not suggest the three major extant forms are experiencing bottleneck effects. Genetic differentiation among forms was notably high despite the presence of hybrids. However, simulations to explore the impacts of hybridization on differentiation and the interpretation of low levels of statistical noise in ADMIXTURE analyses indicate a high likelihood that hybridization beyond the F1 state is not successful. This is further supported by the discovery of widespread differentiation between forms across the genome, indicating that much of the divergence observed has been driven by long term reproductive isolation and drift.

### Hypotheses for high genetic differentiation among forms

Genetic differentiation of the three primary cisco forms in our study (*artedi*, *hoyi*, *kiyi*) was relatively high compared to previous research in cisco using allozymes, mtDNA, microsatellites, AFLPs, and RAD data, which has largely suggested that forms are not frequently diverged within lakes (but see Stott et al., in press; Turgeon, Estoup, & Bernatchez, 1999; Turgeon et al., 2016). For example, Piette-Lauzière et al. (2019) documented neutral *F*_ST_ values near or below 0.01 between forms in small lakes within Algonquin Provincial Park (Ontario, Canada) using RAD data, which were much lower than the *F*_ST_ values of ∼0.05 observed in our study. Unfortunately, genetic data for cisco in the Great Lakes is relatively sparse, however, a previous study using mtDNA and microsatellites did not find evidence of differentiation among forms (Turgeon & Bernatchez, 2003). Results from the microsatellites genotyped in the current study are similar and indicate that these markers are unable to differentiate species in Lake Superior, even though genetic structure was relatively high according to the RAD data. Interestingly, our estimates of genetic divergence among forms more closely mirror two studies in lake whitefish and European whitefish that used RAD sequencing (Feulner & Seehausen, 2019; Gagnaire, Pavey, et al., 2013) than previous studies in cisco that genotyped mtDNA and microsatellites.

The high genetic divergence among forms observed in cisco is not typical of other fishes in the Laurentian Great Lakes. Both lake trout (*Salvelinus namaycush*) and brook trout (*Salvelinus fontinalis*) display significant life history polymorphisms in the Great Lakes, with lake trout exhibiting morphologically distinct ecotypes related to depth and brook trout exhibiting both fluvial and adfluvial life histories. Two recent studies used RAD sequencing to investigate life history polymorphism in these species, and neither was able to document strong signals of divergence among forms (Elias, McLaughlin, Mackereth, Wilson, & Nichols, 2018; Perreault-Payette et al., 2017). It is possible that divergence within these forms was reduced through introgression mediated by stocking, as these species were stocked heavily, whereas cisco was not (Baillie, Muir, Scribner, Bentzen, & Krueger, 2016; Wilson et al., 2008). However, it is also possible that reproductive isolation among cisco forms is more complete, reducing the potential for introgression to erode divergence among forms.

Very little is known about the spawning biology of forms outside of *artedi*, although our observation of relatively frequent F1 hybrids between ART-HOY and HOY-KIY suggests that there is a least some overlap in reproductive timing among the three forms. It also notable that we did not observe any putative hybrids between ART-KIY. These three cisco forms are encountered in different depths in Lake Superior, with *artedi* inhabiting waters < 80 m deep, *hoyi* inhabiting depths between 60-160 m, and *kiyi* inhabiting depths from 80-200 m (Eshenroder et al., 2016). This depth stratification likely explains our observation that *hoyi* hybridizes with *artedi* and *kiyi* but *artedi* and *kiyi* do not hybridize with each other.

Our observation that F1 hybrids are relatively common but hybrid backcrosses appear to be uncommon is consistent with substantially reduced fitness and genetic incompatibilities in hybrid backcrosses. Successful hybridization can homogenize genetic structure within a few generations, as evidenced by our simulations and by a large body of literature in species such as European whitefish and cichlids (reviewed in Seehausen, 2006). However, the fact that we do not observe successful backcrosses suggests that negative interactions between hybrid genomes, such as Dobzhansky-Muller incompatibilities (Dobzhansky, 1936; Muller, 1942), may be present in these backcrosses (Dagilis, Kirkpatrick, & Bolnick, 2019; Ellison & Burton, 2008). Multiple lines of evidence for hybrid incompatibilities have been found in a sister taxon of cisco, lake whitefish (*Coregonus clupeaformis*, reviewed in Bernatchez et al., 2010). For example, Rogers and Bernatchez (2006) found that hybrid backcrosses had ∼6 times higher mortality than F1 crosses, Renaut, Nolte, and Bernatchez (2009) found disruption of gene expression pattern in backcrosses, and Whiteley, Persaud, Derome, Montgomerie, and Bernatchez (2009) documented reduced sperm performance in backcrosses. Our results combined with those from lake whitefish provide evidence that hybrid backcrosses may experience dramatically reduced fitness in cisco. However, future research is necessary to empirically test this hypothesis and investigate potential mechanisms for genomic incompatibilities in hybrid backcrosses.

### Genetic differentiation across the genome

Comparison of patterns of genomic divergence found in our study with previous research suggests that diversification of cisco forms in the Great Lakes is likely polygenic and that these forms have been isolated without gene flow for a relatively long period of time. We identified over 100 significantly differentiated genomic windows in our study, and genetic differentiation among forms was consistently high across the genome. This result is similar to two other investigations of genomic divergence in coregonines (Feulner & Seehausen, 2019; Gagnaire, Pavey, et al., 2013), but differs substantially from genome scans in some other salmonids that have revealed supergenes responsible for substantial phenotypic divergence across large geographic areas and differing genetic backgrounds (e.g. Barson et al., 2015; Pearse et al., 2018; Prince et al., 2017). Specifically, Feulner and Seehausen (2019) investigated genomic divergence in three species of whitefishes (*Coregonus* spp) from two lakes in Switzerland and found high divergence on all chromosomes, and Gagnaire et al. (2013) assessed divergence between species pairs of lake whitefish (*C. clupeaformis*) across five lakes in the St. John River Basin (Maine, USA and Québec, Canada) and also found consistently high differentiation across the genome. By comparison, Barson et al. (2015) found a single genomic region that is largely responsible for age-at-maturity in Atlantic salmon (*Salmo salar*), Prince et al. (2017) discovered a small genomic region that is highly associated with run timing in steelhead (*Oncorhynchus mykiss*) and Chinook salmon (*Oncorhynchus tshawytscha*), and Pearse et al. (2018) documented a large inversion associated with life history polymorphism (anadromy vs residency) in steelhead. Taken together, these results suggest divergence of coregonines in both Europe and North America is likely polygenic. This conclusion is supported by a large body of research in lake whitefish demonstrating that traits that differ among species (e.g. body size, growth rate) are controlled by many quantitative trait loci (QTL) (Gagnaire, Normandeau, et al., 2013; Laporte et al., 2015; Rougeux, Gagnaire, Praebel, Seehausen, & Bernatchez, 2019). Additionally, signatures of parallel evolution at the genomic level are relatively rare in coregonines (Feulner & Seehausen, 2019; Gagnaire, Pavey, et al., 2013), providing evidence that adaptive divergence in this taxon is likely controlled by many genes of small effect that generally differ across speciation events.

Even when speciation is controlled by many genes of small effect, patterns of genetic differentiation may not be homogenous across the genome (Nosil & Feder, 2012; Nosil, Funk, & Ortiz-Barrientos, 2009). Heterogenous genomic divergence is hypothesized to be especially common during sympatric speciation (i.e. divergence with gene flow), as selectively advantageous loci that are clustered together are protected from between population recombination (Via, 2012; Via & West, 2008). Prolific islands of divergence have been identified in stickleback (*Gasterosteus aculeatus*) forms that diverged within the last 150 years (Marques et al., 2016) and in species with high gene flow, such as stick insects (*Timema cristinae*, Soria-Carrasco et al., 2014) and *Heliconius* butterflies (Nadeau et al., 2012). Often these islands are found in areas of low recombination (Burri et al., 2015; Samuk et al., 2017), as we observed in LG Cart21. However, as populations drift apart, high levels of divergence accumulate across the genome, making it difficult to differentiate genomic islands involved in the original divergence from accumulated genetic drift in populations that have been isolated for a long time period (Via, 2012). This pattern was observed in sockeye salmon (*Oncorhynchus nerka*), where an island of divergence that is highly visible across multiple drainages in Alaska was partially obscured in the Copper River, which has experienced lower gene flow and therefore much more genetic drift than the other systems (Larson et al., 2019). The high and relatively homogenous patterns of genetic differentiation observed in our study suggest that genomic islands involved in diversification may be obscured in a similar fashion.

### Conservation implications

We documented high genetic differentiation among the three major cisco forms in Lake Superior and, based on this information, we suggest that separate conservation units could be constructed for each form. This strategy differs from the current conservation paradigm for cisco, which recommends that the entire *C. artedi* species complex be considered *C. artedi* sensu lato (translation, in the broad sense) and that units of conservation should be designed to preserve environments that have facilitated the evolution of different forms (i.e. lakes) rather than on forms at a larger scale (Turgeon & Bernatchez, 2003; Turgeon et al., 2016). These recommendations were informed by the best available data, which up to this point, have been generated with commonly employed markers for the detection of taxonomic and conservation units - AFLPs, mtDNA, and microsatellites (Reed et al., 1998; Turgeon & Bernatchez, 2003; Turgeon et al., 2016) - none of which have been able to consistently resolve different forms within the Great Lakes (but see Stott et al., in press for a single lake example). Our results suggest that “last generation” markers may be insufficient for capturing differentiation in evolutionarily young species, such as cisco, and highlight the utility of genomic data for designating conservation units in these species. However, it is important to note that we only surveyed animals from one lake, and a larger genotyping effort across the Great Lakes is necessary to accurately inform conservation units for Great Lakes ciscoes. Additionally, our ability to draw conclusions regarding the relationships of the rare forms *nigripinnis* and *zenithicus* to the three major forms was limited by low sample size, therefore, genotyping additional contemporary and/or historic specimens may help resolve the placement these forms.

The potential of genomic data to revolutionize construction of conservation units has been frequently discussed (reviewed in Allendorf et al., 2010; Funk et al., 2012), and many studies have found that genomic data provides increased resolution for delineating population structure compared to last generation markers, such as microsatellites (Hodel et al., 2017; Vendrami et al., 2017; Wagner et al., 2013). However, conservation units that were constructed with these last generation markers have generally proven to be robust and are usually only updated slightly, if at all, based on genomic data (e.g. Hecht, Matala, Hess, & Narum, 2015; Larson et al., 2014; Moore et al., 2014). Our findings do not follow this pattern and suggest that current conservation units for cisco may not accurately reflect the diversity of this species. This finding has larger implications for constructing conservation units. Specifically, our data suggest that conservation units constructed based on data from last generation markers that found either low or no significant structure among groups of animals with different phenotypes could potentially be re-evaluated with genomic tools to ensure within-species diversity is being adequately conserved.

### Conclusions and future directions

Our study provides the first evidence that cisco forms within the Great Lakes are genetically differentiated. We documented high genetic differentiation among the three major forms in Lake Superior, and highly differentiated markers were distributed across the genome, with islands of divergence found on nearly every linkage group. Additionally, we identified putative F1 hybrids but no hybrid backcrosses, suggesting that fitness breakdown of backcrosses may aid in maintaining differentiation of these forms. The results of this study provide the foundation for a new understanding of the ecology and evolution of the *C. artedi* species complex within the Great Lakes. The ability to differentiate forms with genomics provides researchers with a powerful tool for ground truthing morphological phenotypes and identifying cisco species at any life stage. In particular, the ability to identify larval ciscoes will allow researchers to estimate recruitment, which is vital for management and conservation, and will also significantly improve our understanding of early life history characteristics and reproductive dynamics in this species. Finally, our results suggest that some management units created using last generation markers may not adequately reflect species diversity and could be re-evaluated with genomic data.

## Supplementary data

**Table S1.** Summary statistics for 26,789 loci genotyped with RAD sequencing. The “LG” and “cM” columns denote map location, columns ending in “AF” denote population allele frequencies, “*H*_O_” is observed heterozygosity, *H*_E_ is expected heterozygosity, the “Sequence P1 column” is the sequence from the P1 read for each RAD tag, and the “Sequence PE” is the sequence obtained from paired-end assemblies. Form abbreviations are identical to those in Table 1.

**Table S2.** Summary statistics for 9 microsatellite loci included in this study. Abbreviations are total number of alleles per locus (*A*), effective number of alleles per locus (*Neff*), observed heterozygosity (*H*_O_), and expected heterozygosity (*H*_E_). *F*_ST_ (Weir and Cockerham, 1984) was estimated in Genepop and all other statistics were estimated in Genodive.

**Table S3.** Initial assignment scores from Genodive for all sampled individuals using RAD data.

**Table S4.** Assignment scores from Genodive when tests were limited to the three major forms using RAD data.

**Table S5.** Assignment scores from Genodive for the three major forms using microsatellite data.

**Table S6.** Type, number, and proportion of hybrids observed in empirical data and simulated populations. Data simulated in EASYPOP were comprised of 5,000 biallelic loci from three populations (P1-P3) that experienced stepping-stone migration (m=0.05) for 2, 5, or 10 generations (G2, G5, G10) after 1,000 generations of low migration (m=0.001) to approximate an *F*_ST_ similar to that observed between our three major cisco forms (ART, HOY & KIY, *F*_ST_ ≈0.05).

**Fig S1.** Levels of genetic admixture in three simulated populations after 2, 5, and 10 generations of stepping-stone migration. Simulations were run with random mating of 1,000 females and 1,000 males in each population using 5,000 biallelic loci, and preliminary conditions that produced a similar level of differentiation observed in our RAD dataset among forms (*F*_ST_≈0.05; 1,000 generations with an island migration rate of 0.001). Each ADMIXTURE plot represents a random subset of 50 males and 50 females from each population.

**Fig S2.** Overall *F*_ST_ for all loci that could be placed on the linkage map. Each dot is a marker and red lines indicate genomic windows that were significantly differentiated from the rest of the genome according to kernel smoothing analysis.

**Fig S3.** Pairwise-*F*_ST_ for the ART-HOY comparison for all loci that could be placed on the linkage map. Each dot is a marker and red lines indicate genomic windows that were significantly differentiated from the rest of the genome according to kernel smoothing analysis. Form abbreviations are in Table 1.

**Fig S4.** Pairwise-*F*_ST_ the ART-KIY comparison for all loci that could be placed on the linkage map. Each dot is a marker and red lines indicate genomic windows that were significantly differentiated from the rest of the genome according to kernel smoothing analysis. Form abbreviations are in Table 1.

**Fig S5.** Pairwise-*F*_ST_ the HOY-KIY comparison for all loci that could be placed on the linkage map. Each dot is a marker and red lines indicate genomic windows that were significantly differentiated from the rest of the genome according to kernel smoothing analysis. Form abbreviations are in Table 1.

## Data Archiving Statement

Upon acceptance, demultiplexed sequence data used in this research along with corresponding metadata will be archived in the NCBI sequence read archive using the publicly accessible Genomic Observatories Metadatabase (GeOMe, http://www.geome-db.org/) and microsatellite genotypes will be archived on DRYAD.

## Acknowledgements

We thank Brad Ray with the Wisconsin Department of Natural Resources and Dan Yule and Mark Vinson with the United States Geological Survey for providing the Lake Superior samples used in this project. We are grateful to Kristen Gruenthal for laboratory support during RAD library preparation, to Garrett McKinney for discussions on analysis, and to Julie Turgeon, John Robinson, and Jared Homola for comments on the manuscript. This research was supported by the Turing High Performance Computing cluster at Old Dominion University and was funded by grants from the Great Lakes Restoration Initiative, the Great Lakes Fishery Trust, and the Great Lakes Fish and Wildlife Restoration Act. Any use of trade, product, or company name is for descriptive purposes only and does not imply endorsement by the U.S. Government.

