## Supplementary figures and images for "Genotyping-by-sequencing illuminates high levels of divergence among sympatric forms of coregonines in the Laurentian Great Lakes"

### Fig S3

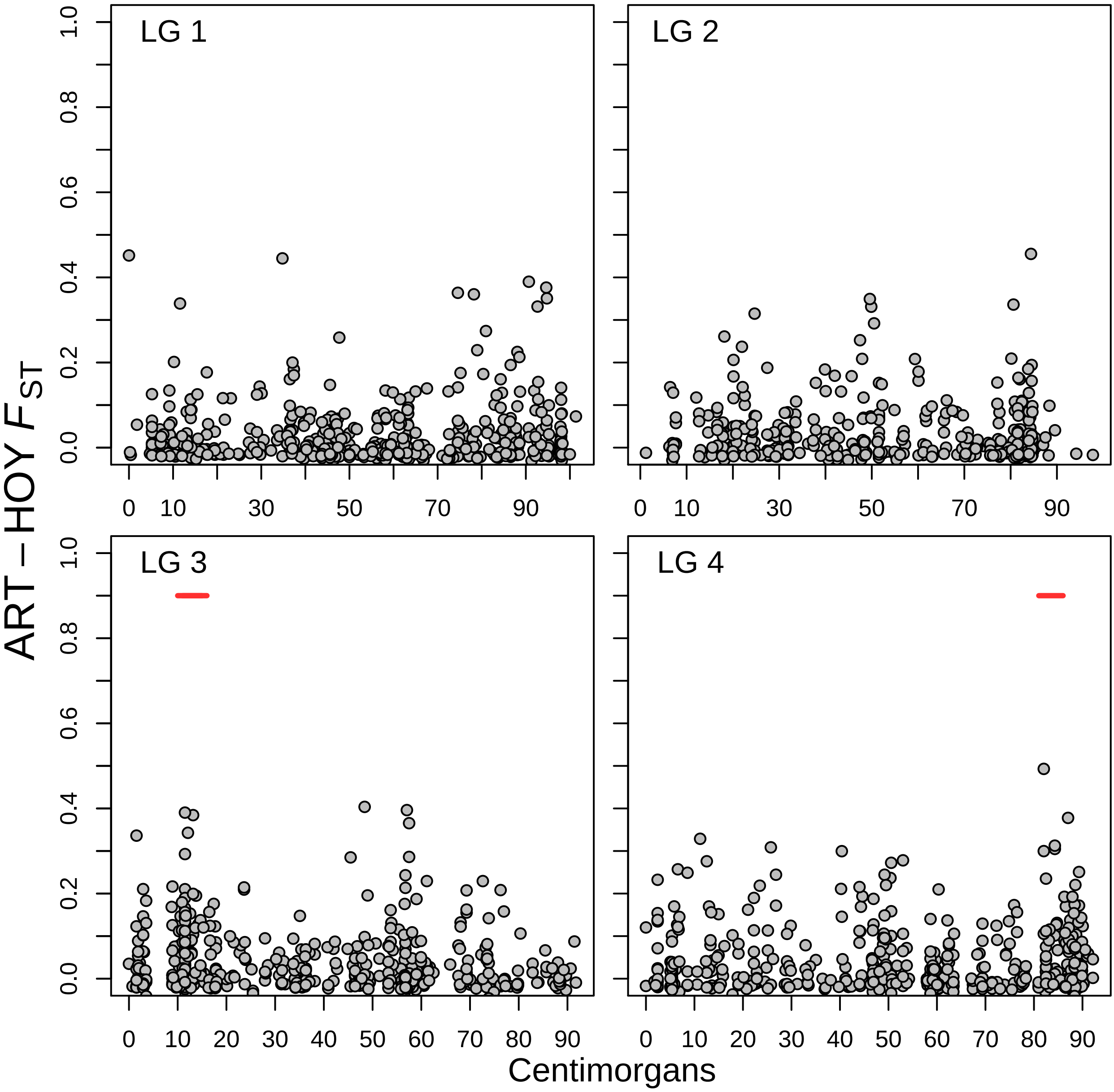

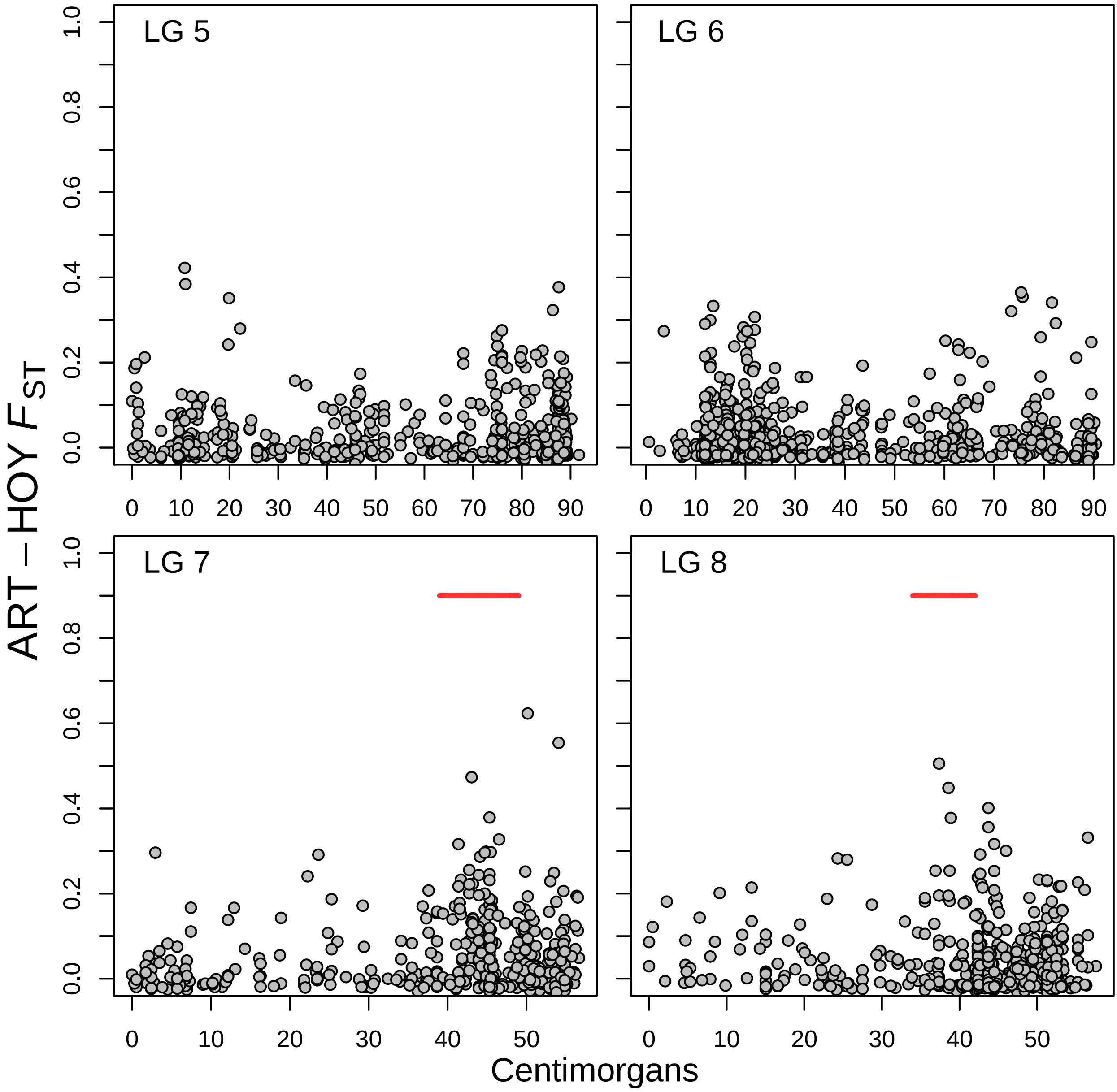

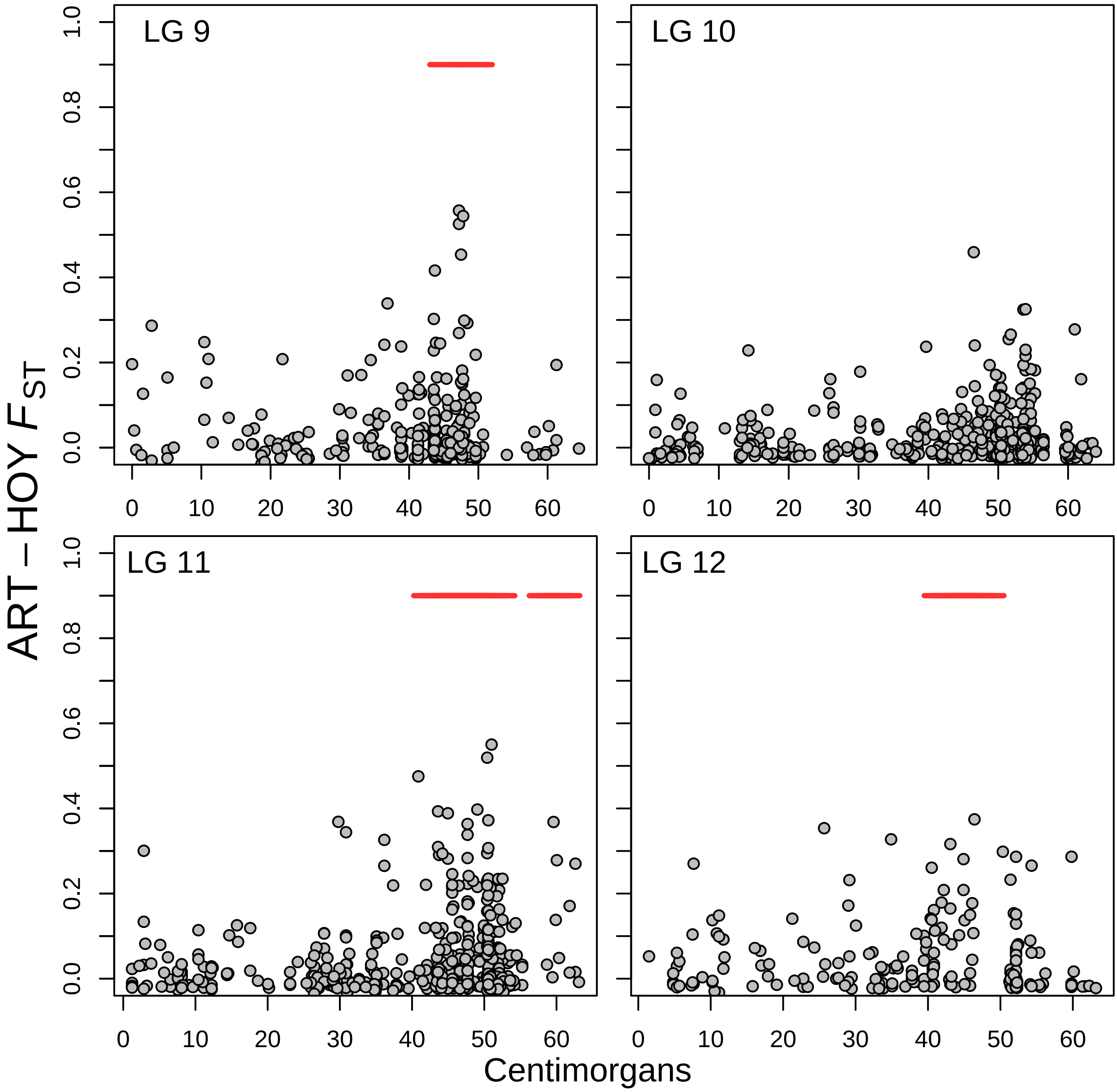

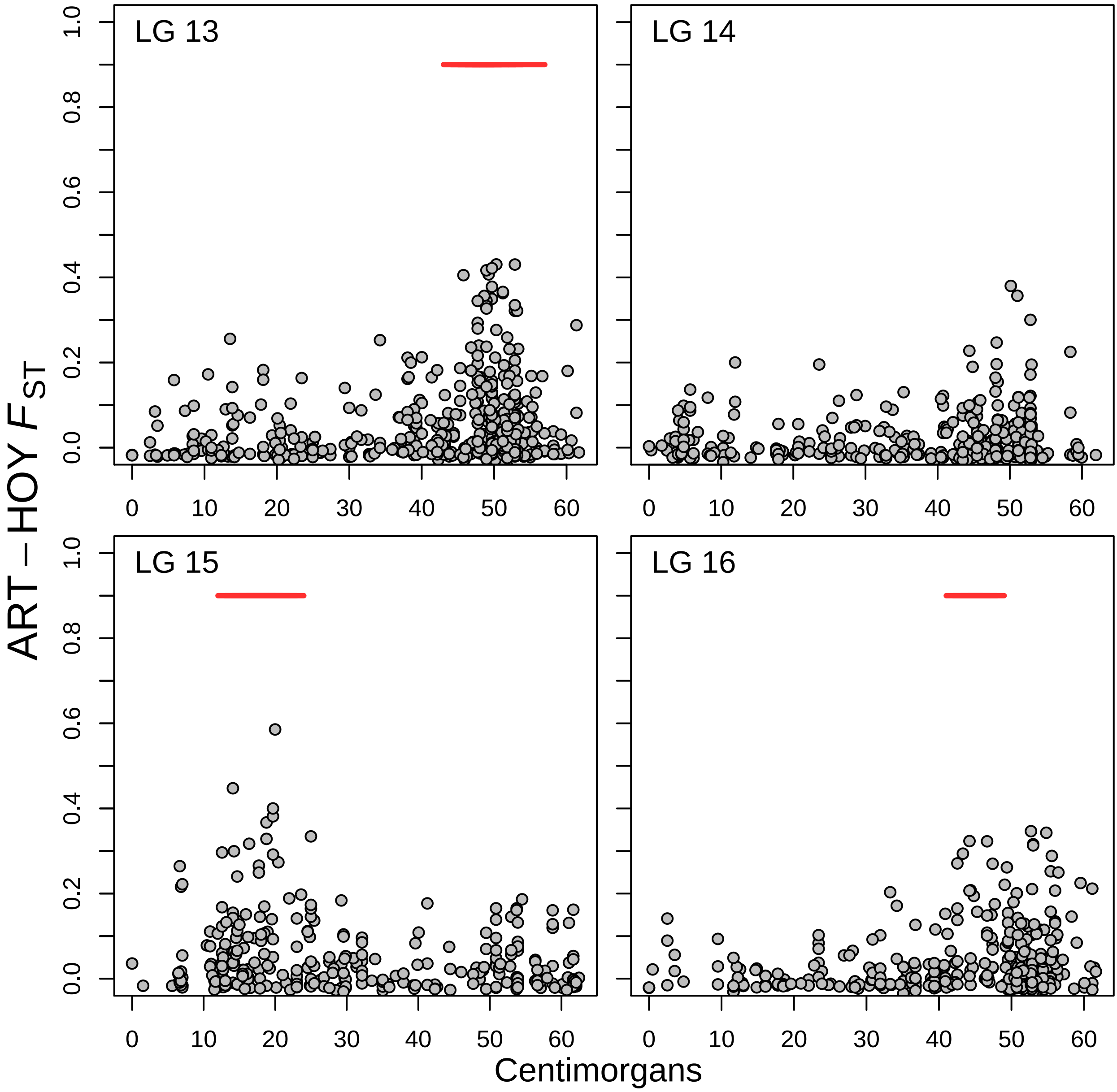

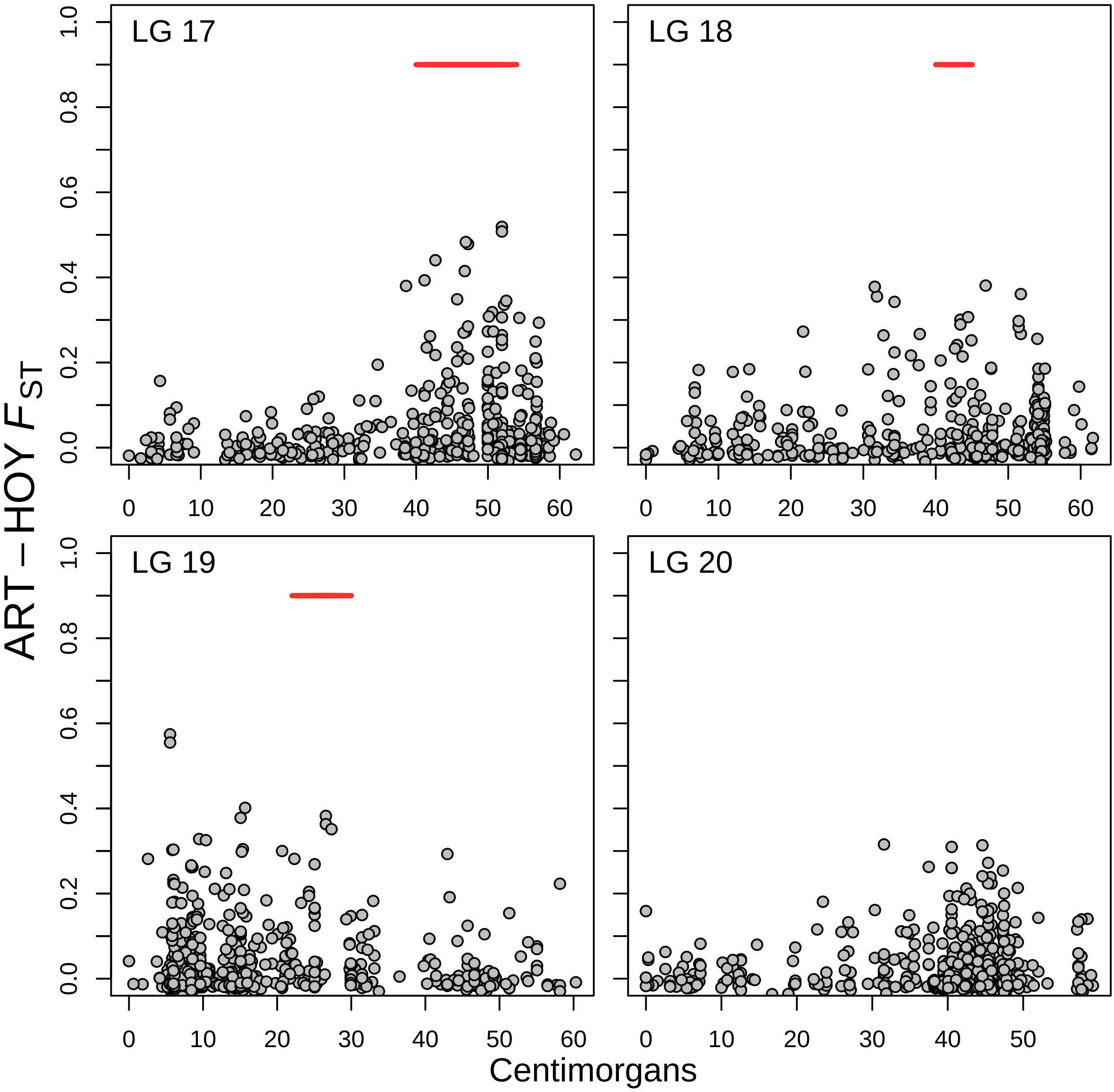

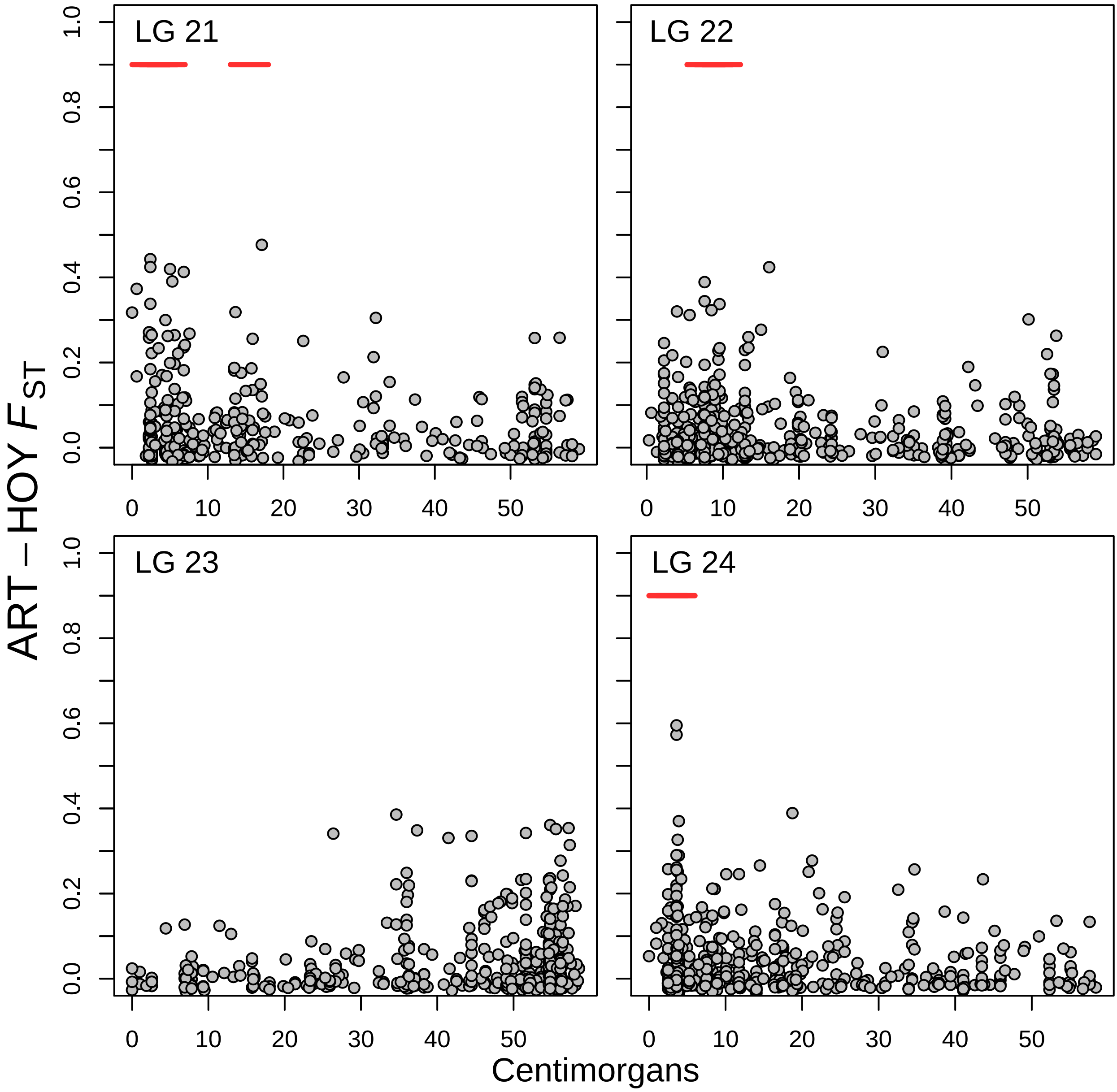

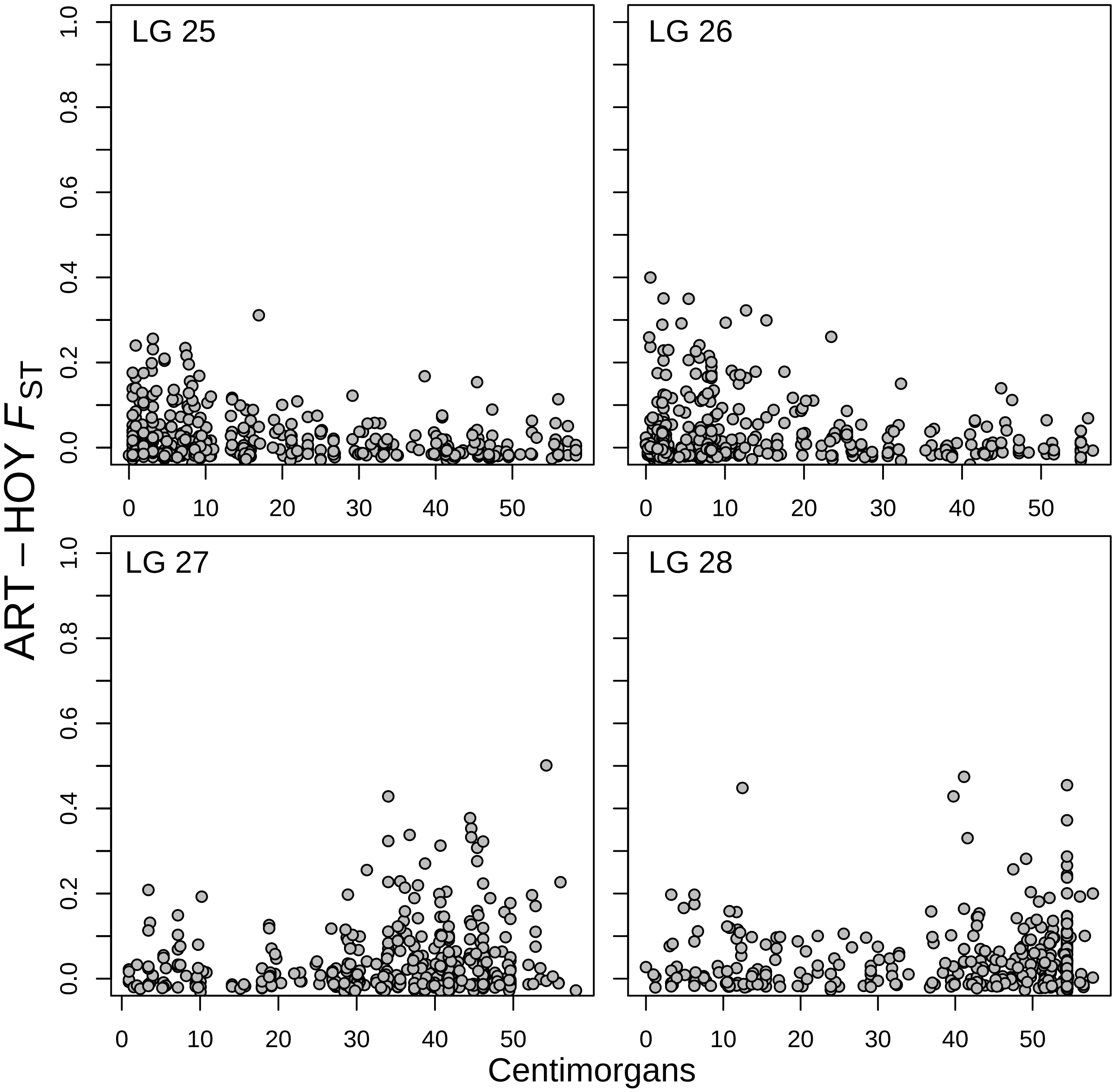

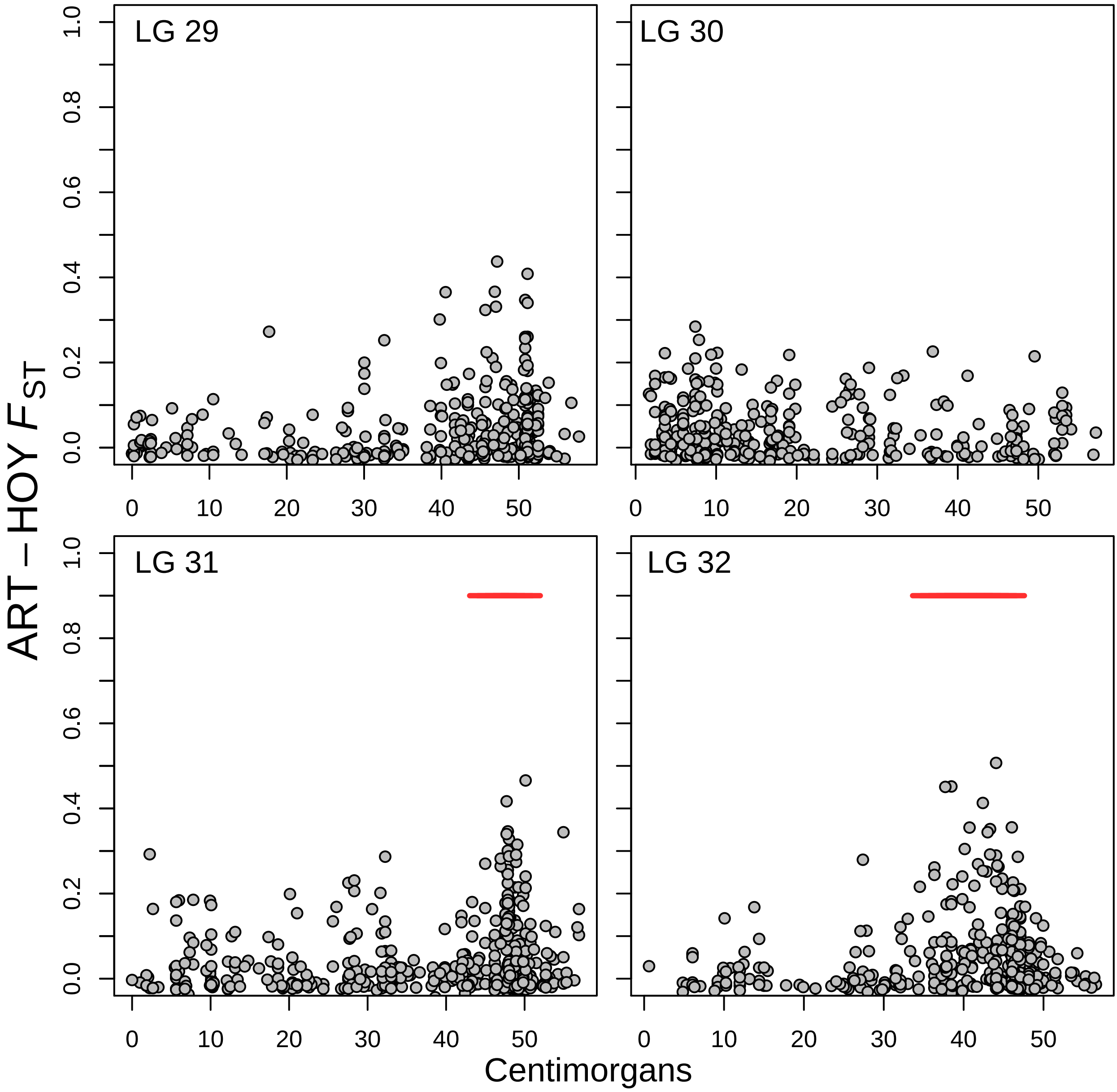

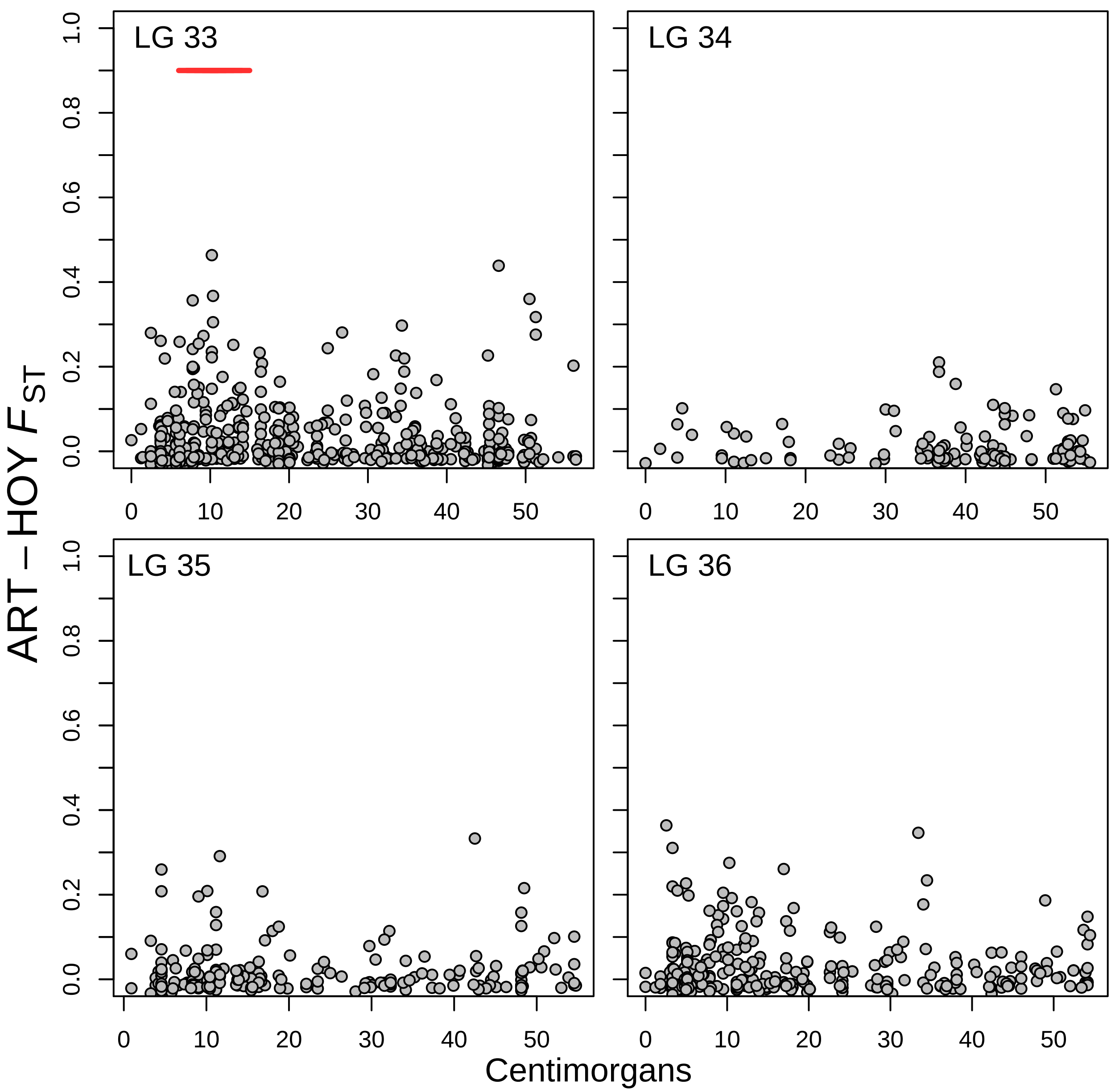

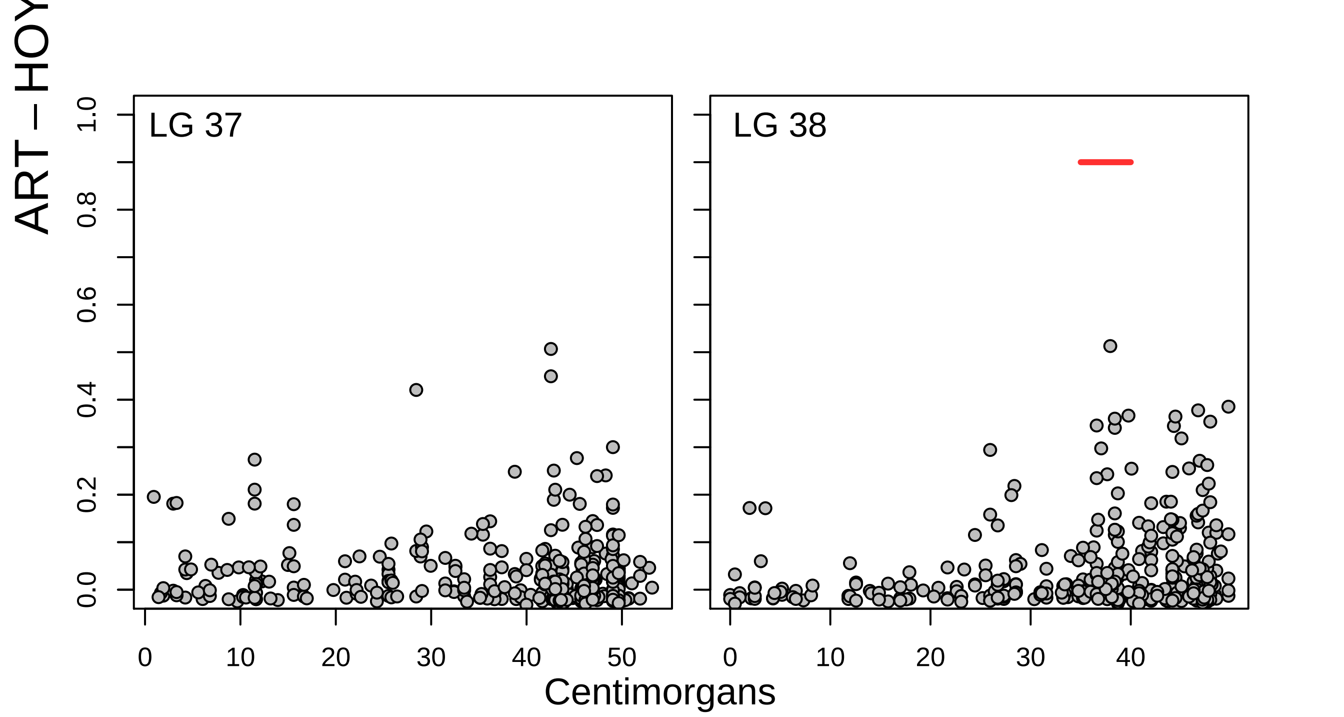

### Fig S4

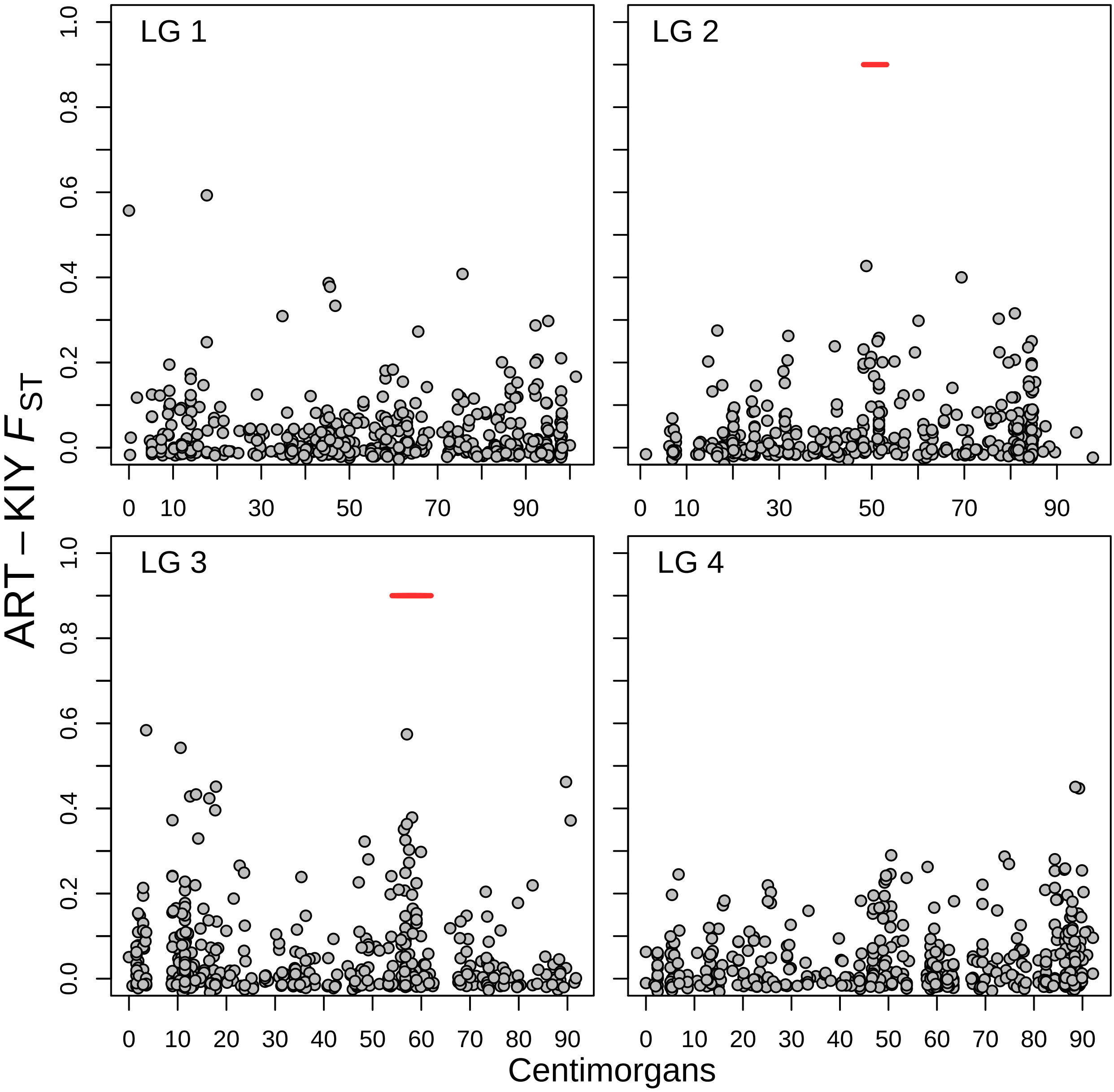

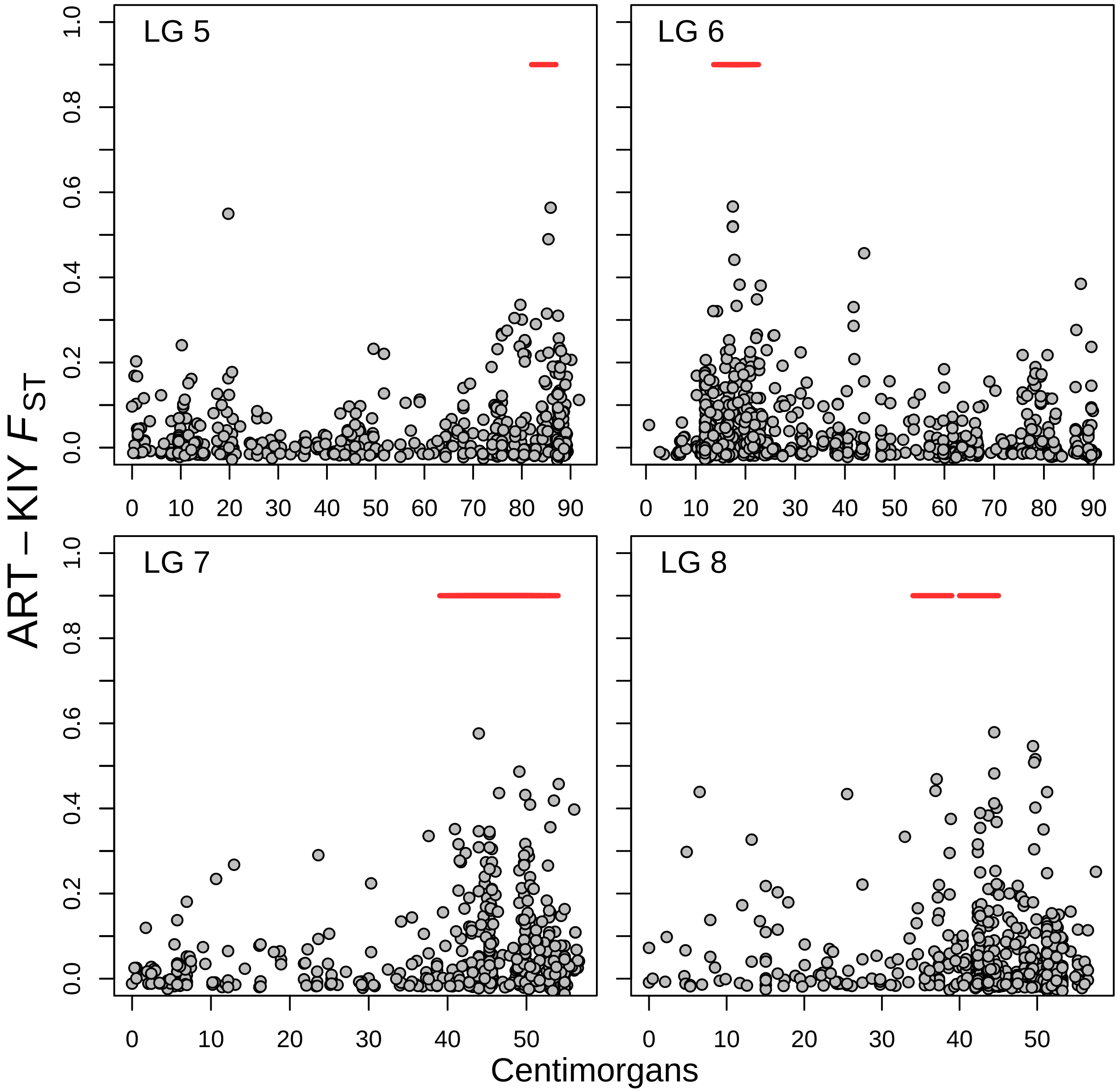

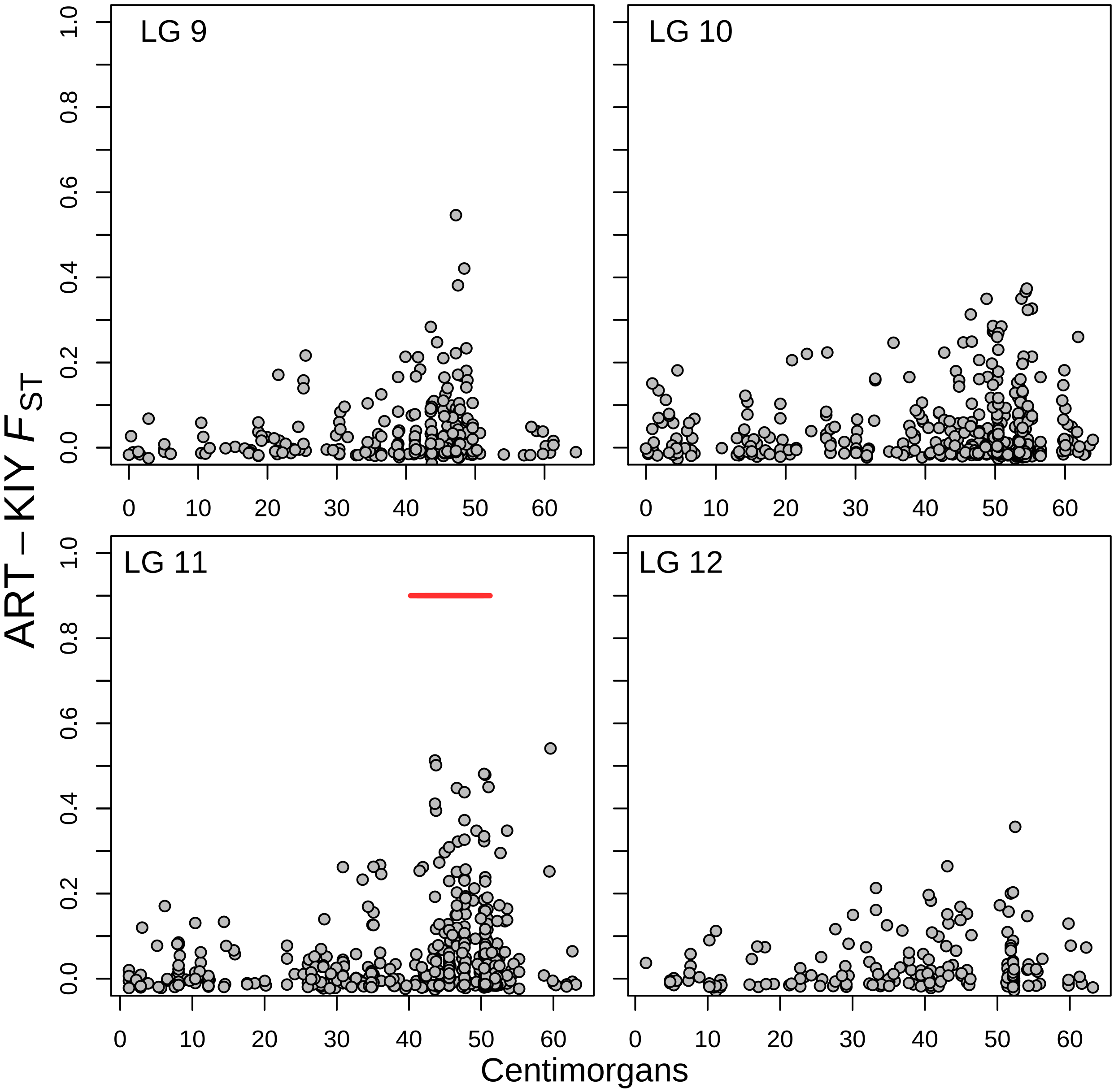

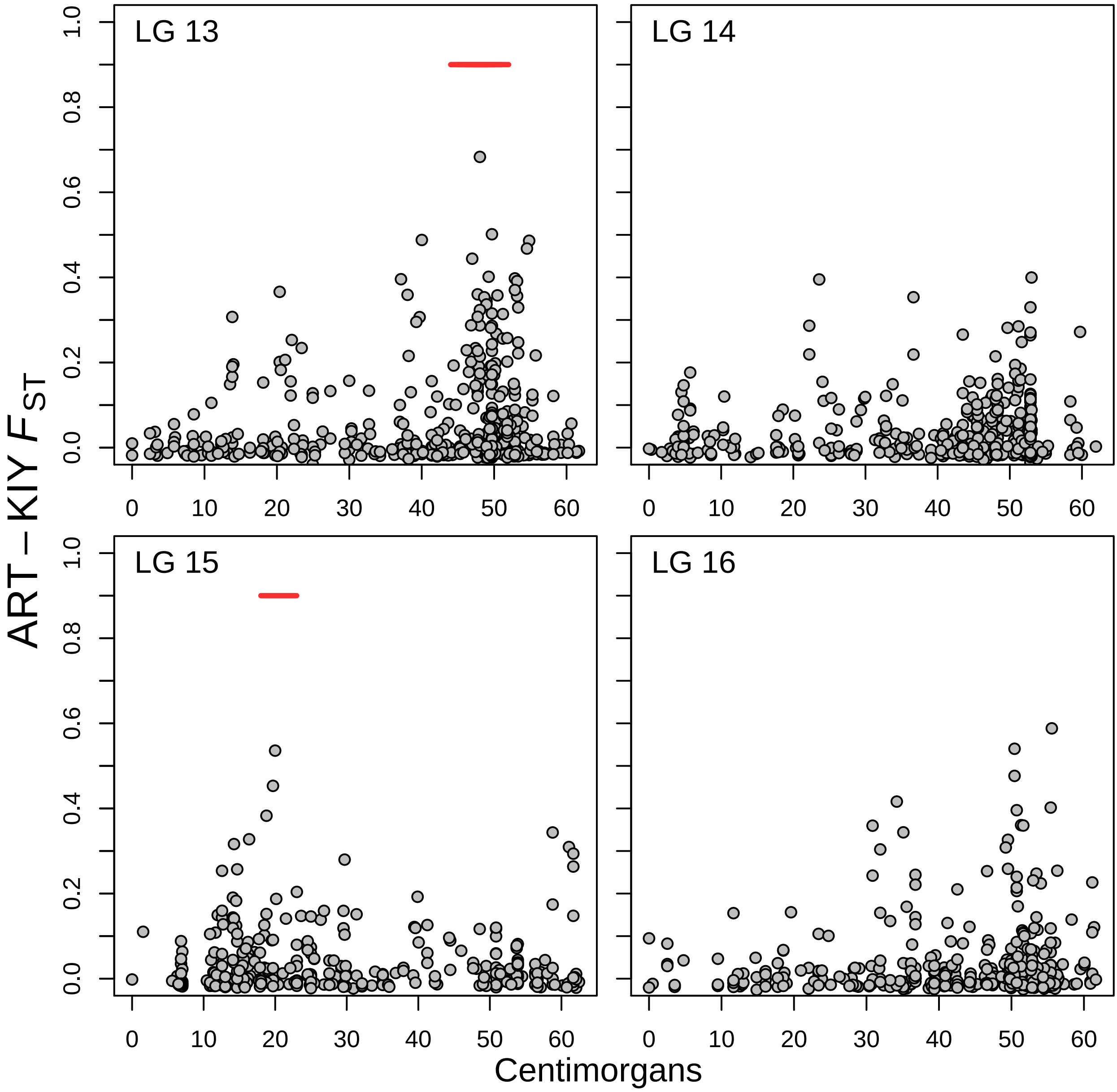

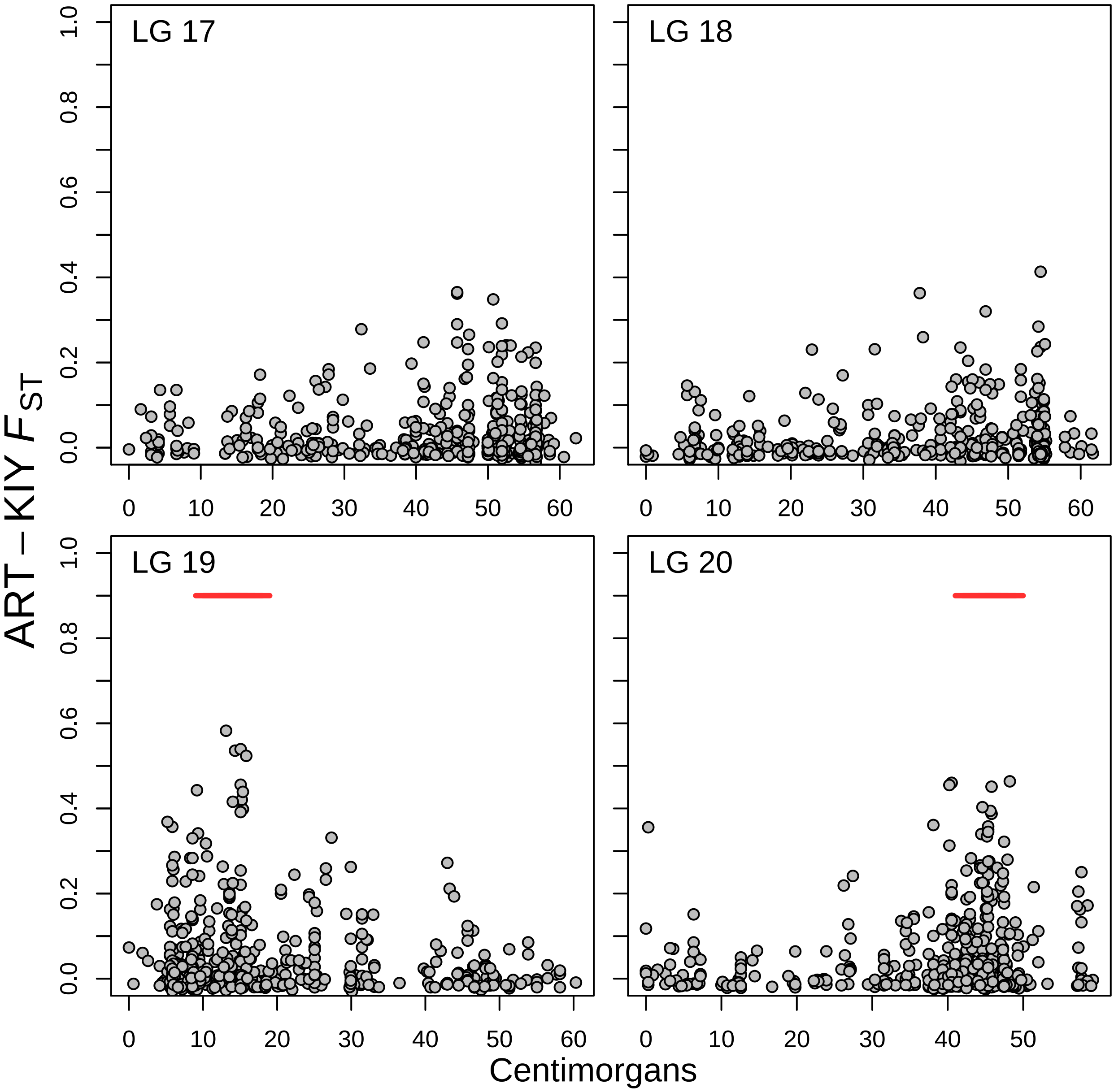

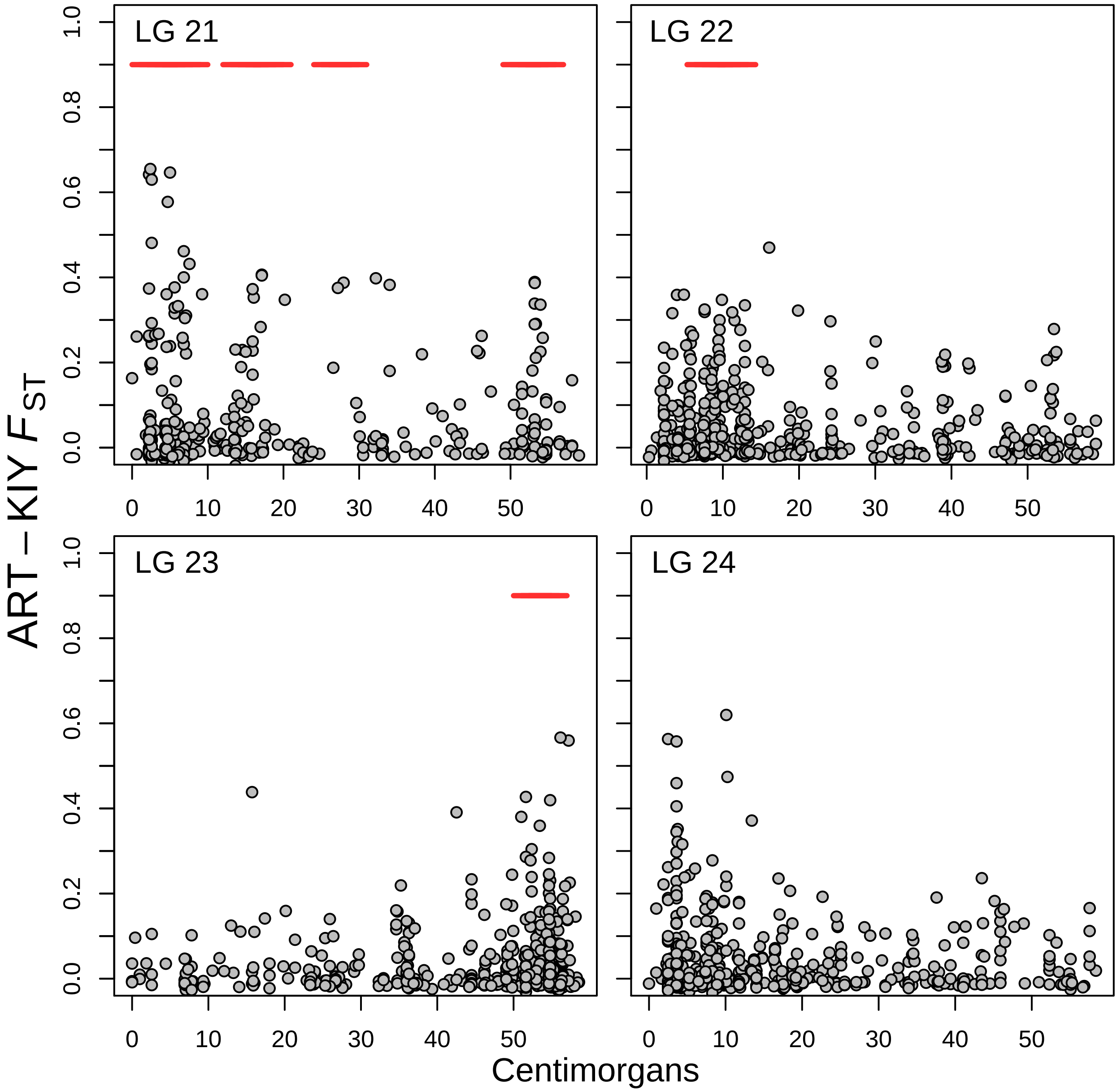

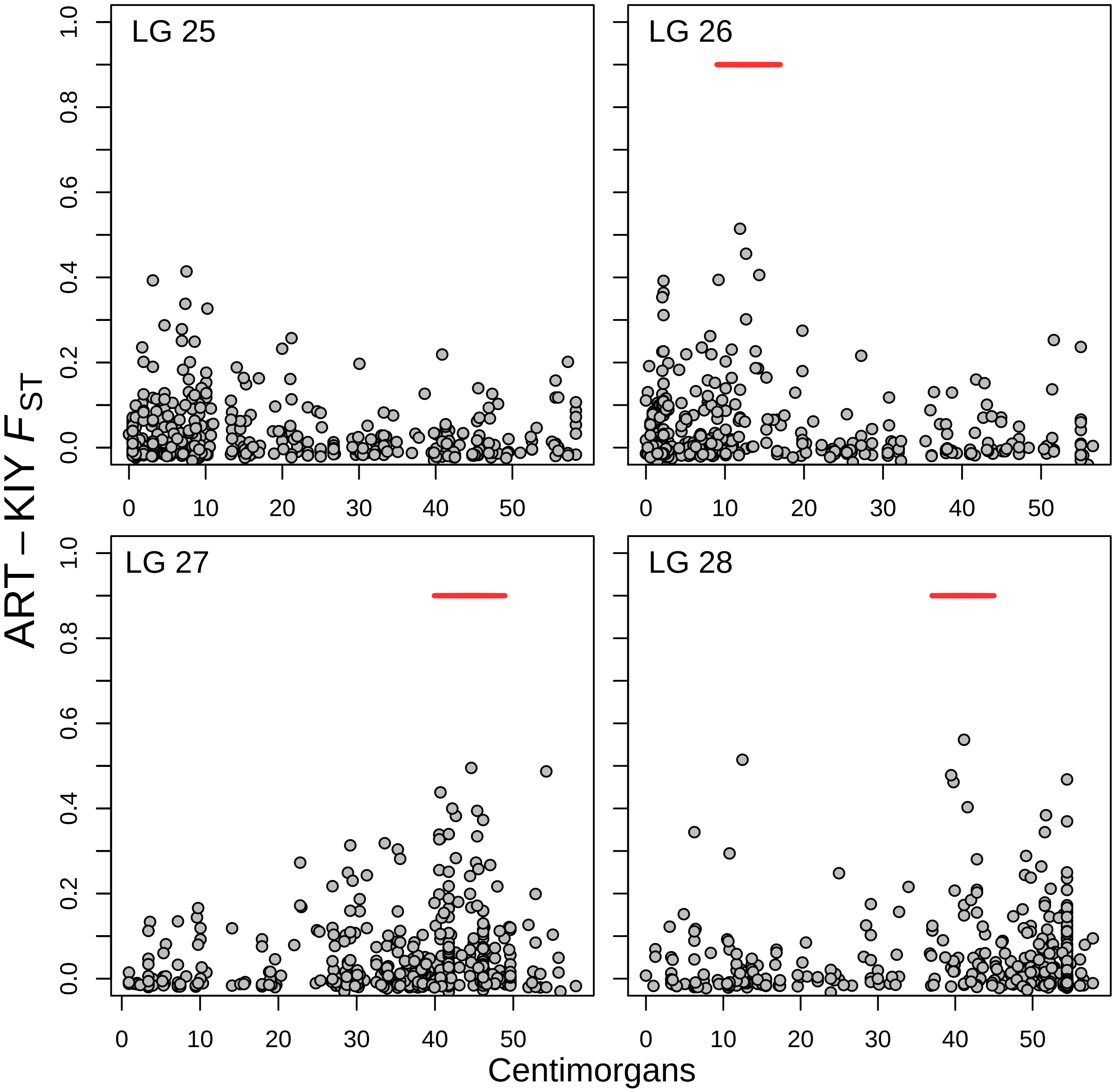

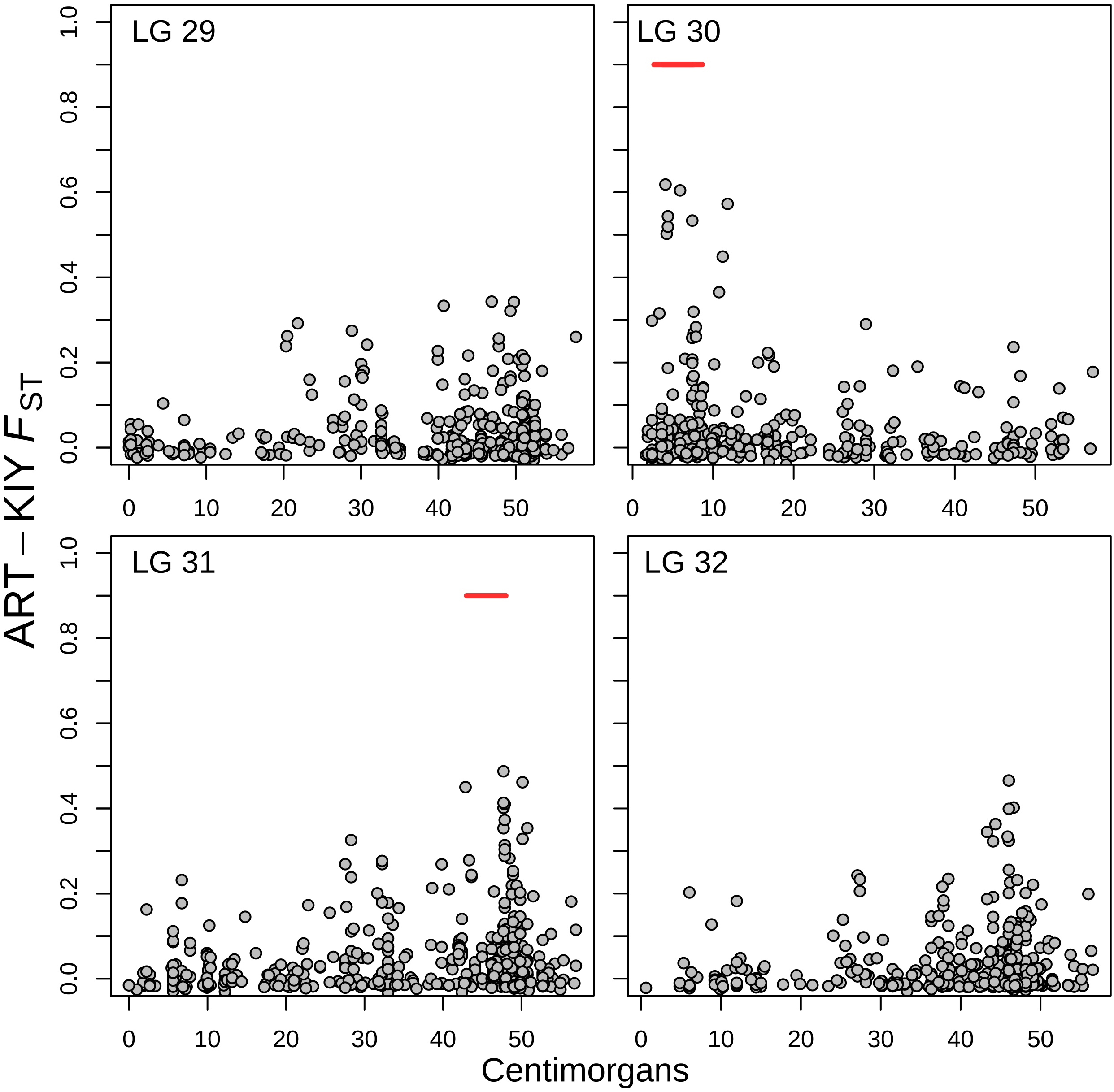

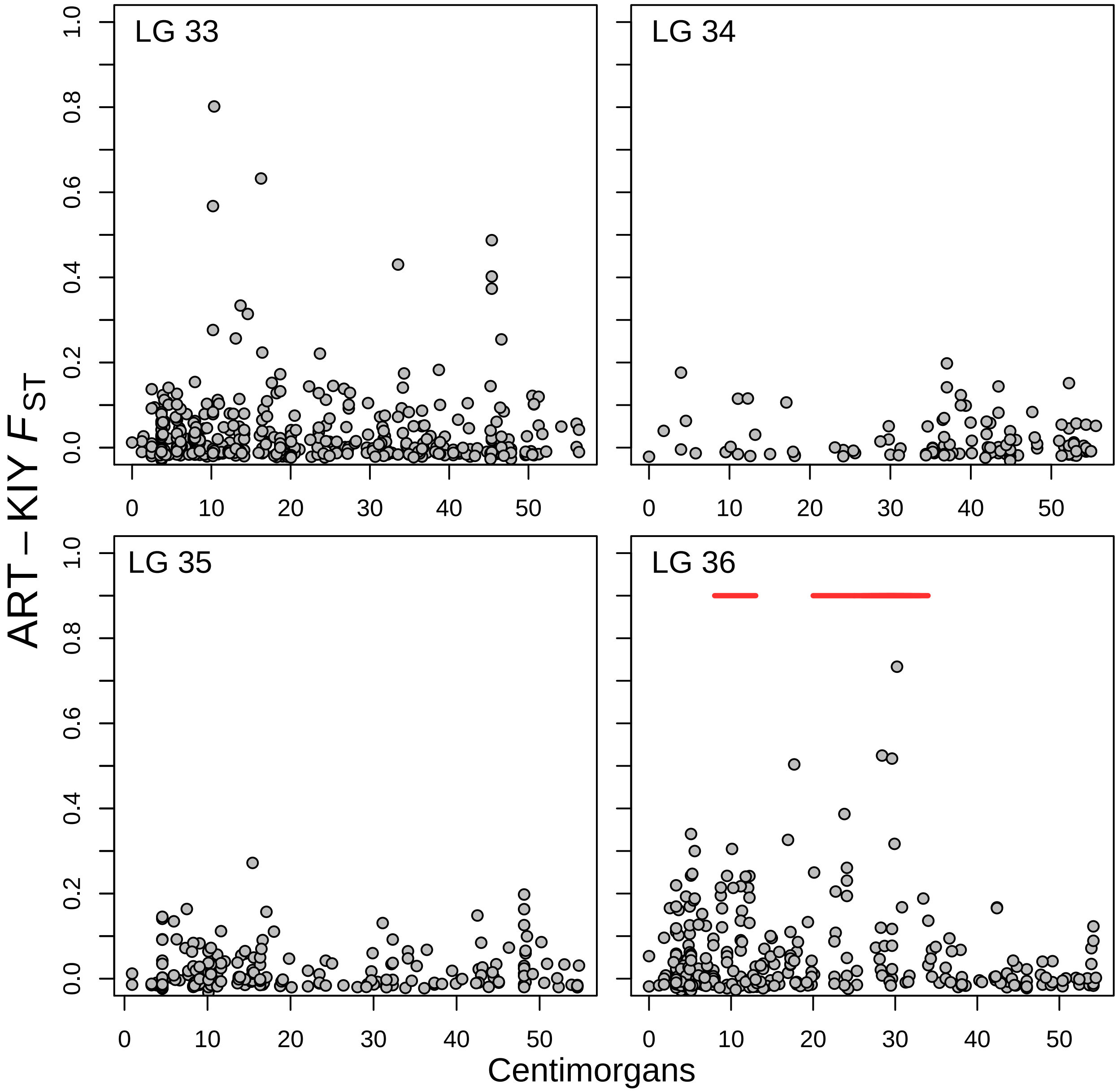

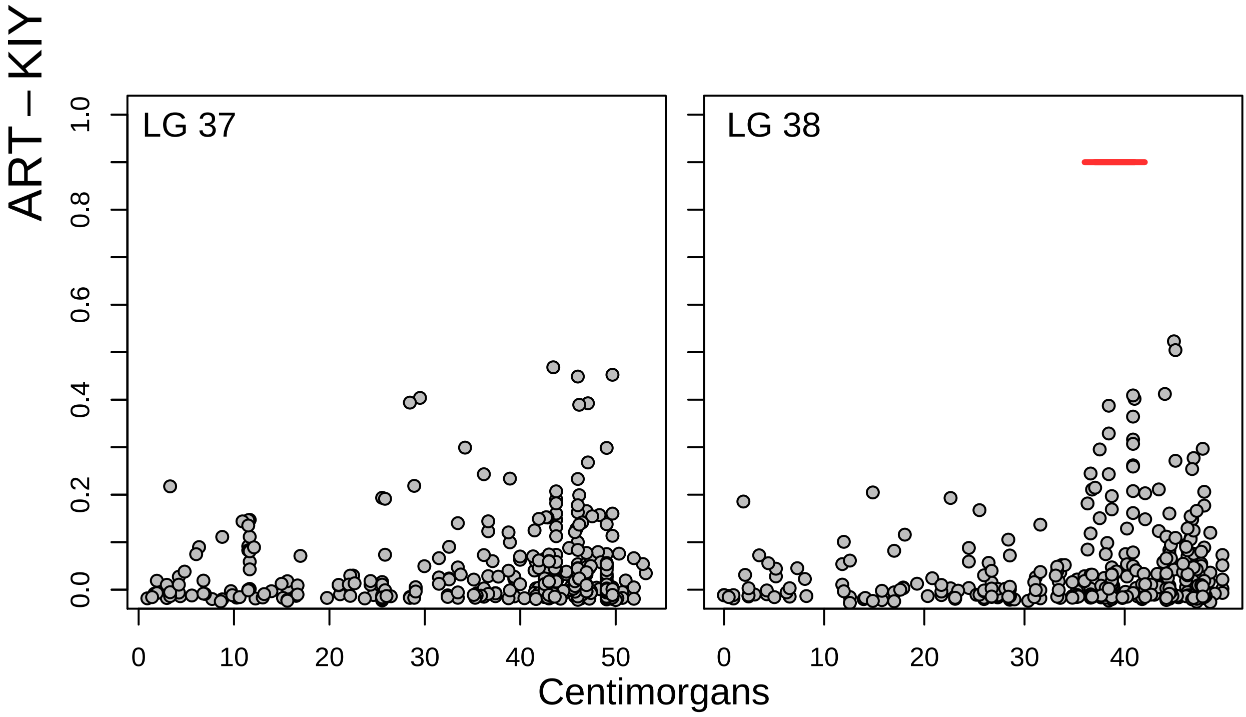

### Fig S5

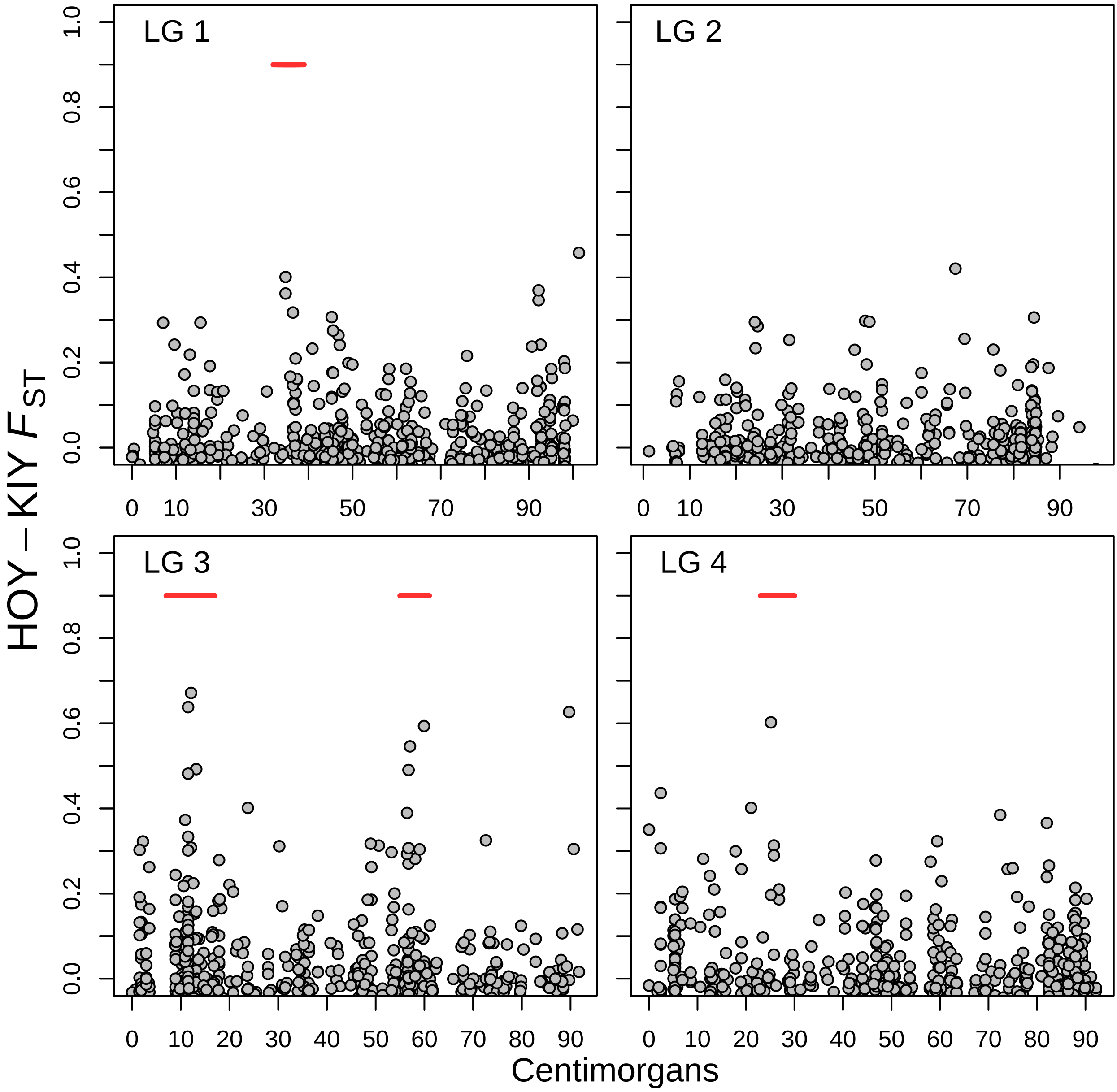

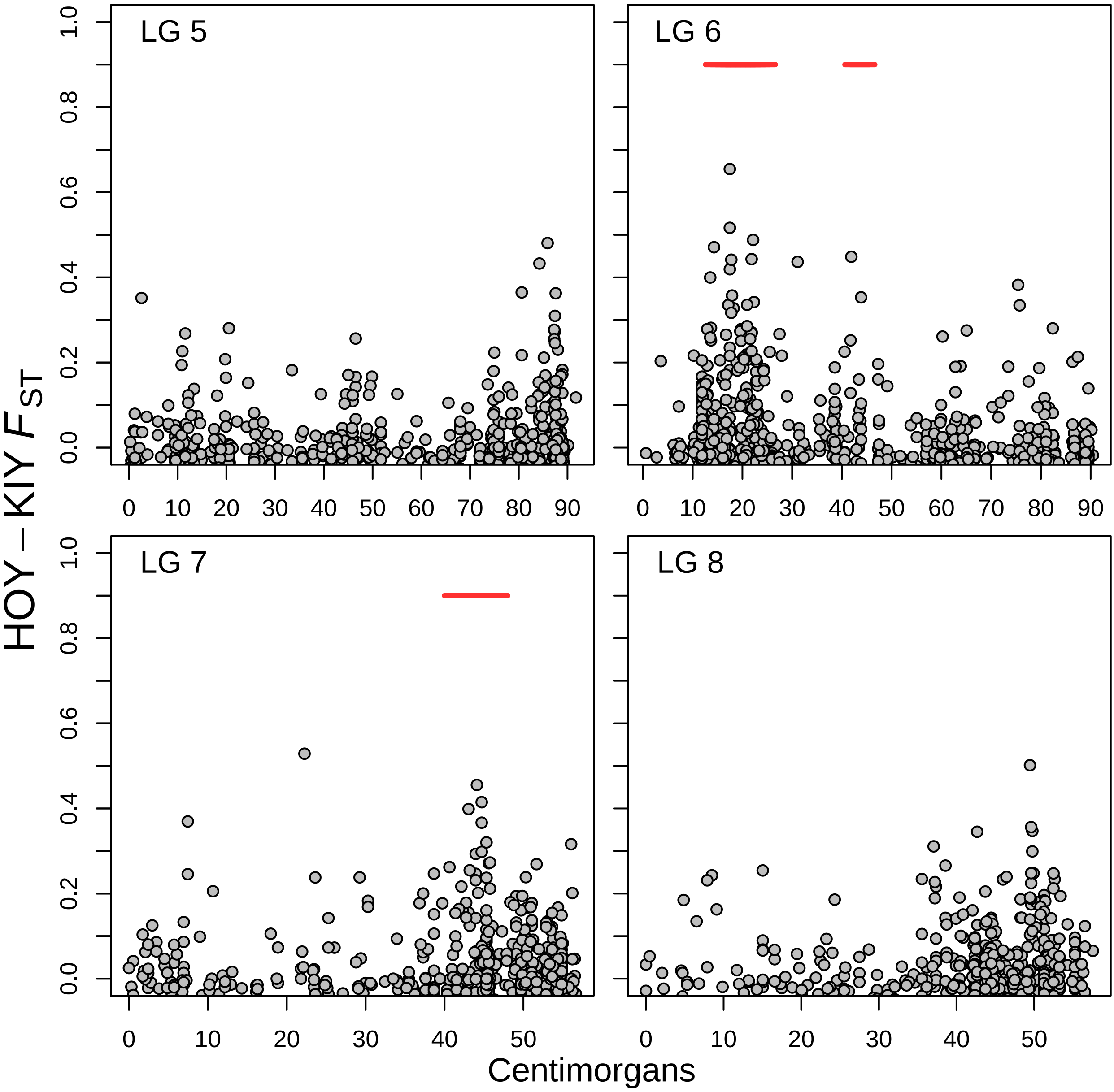

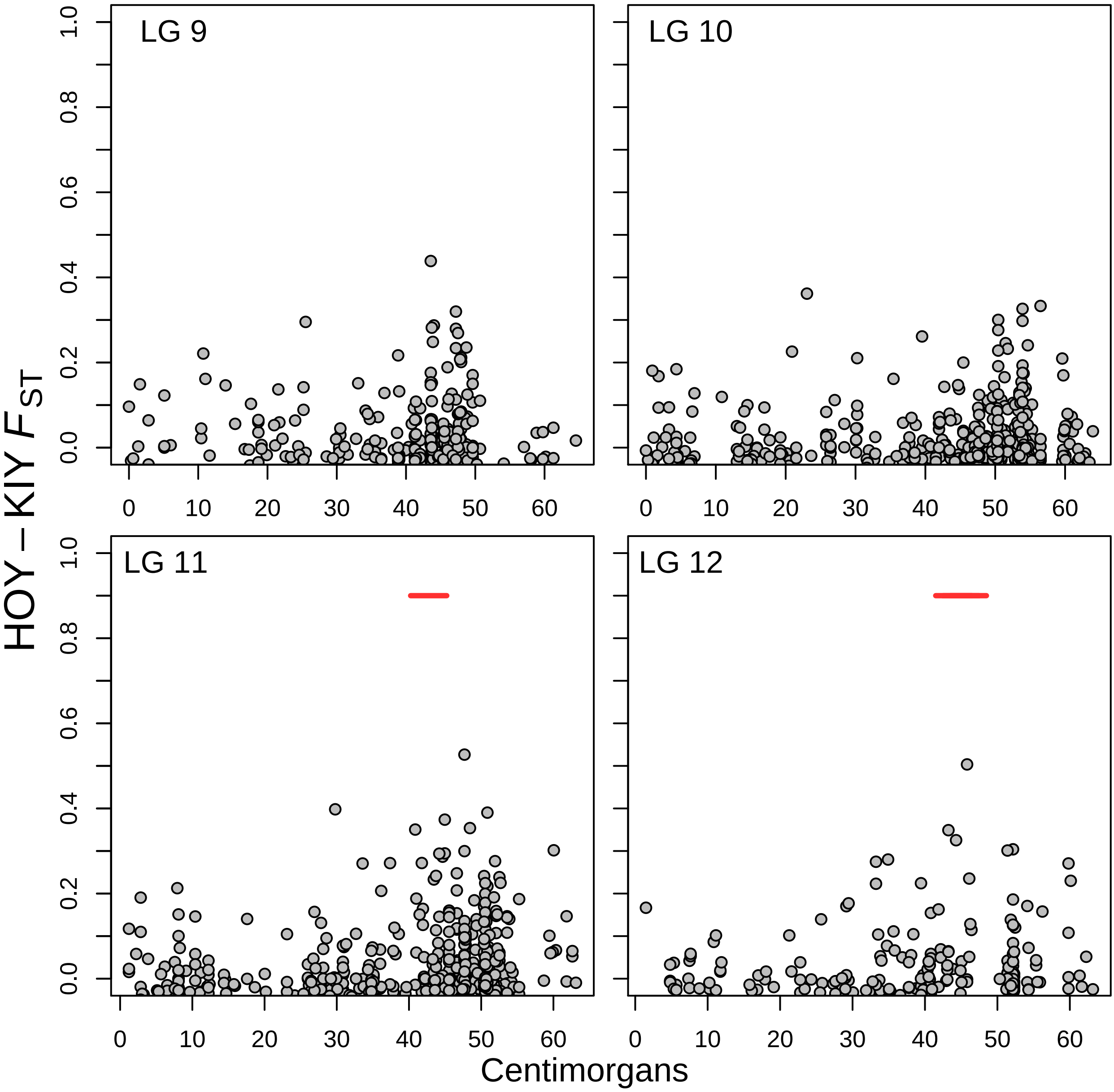

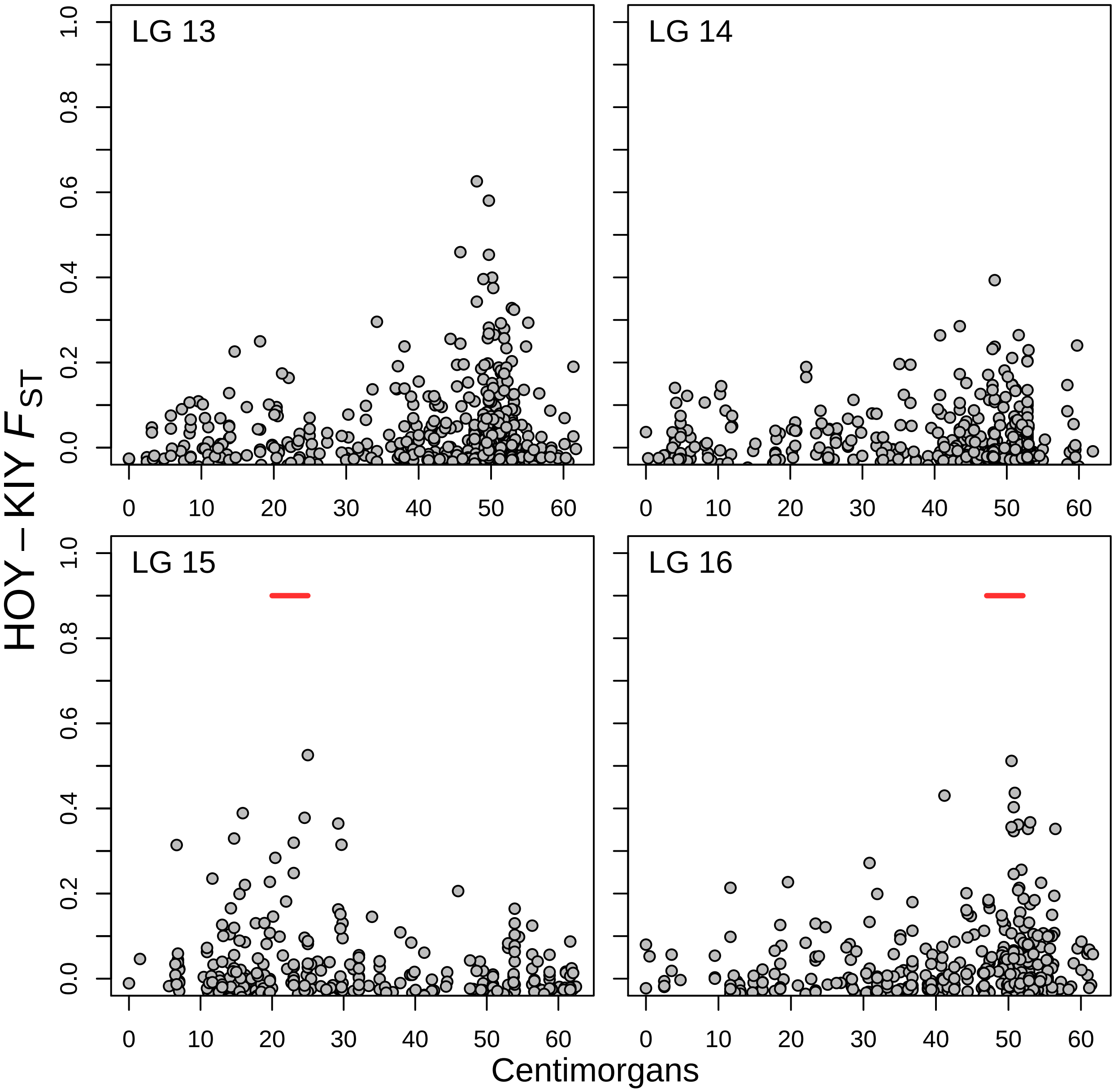

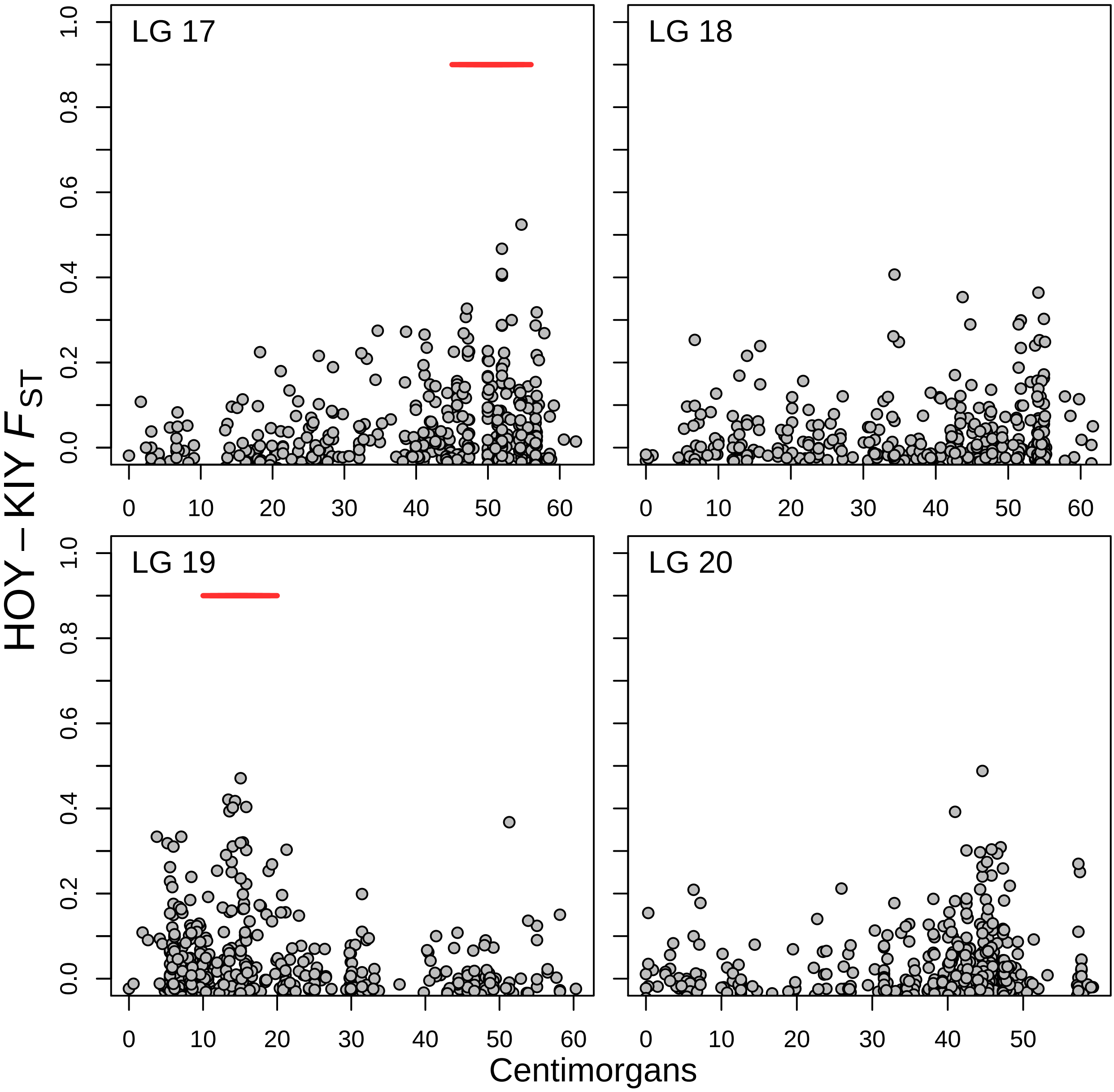

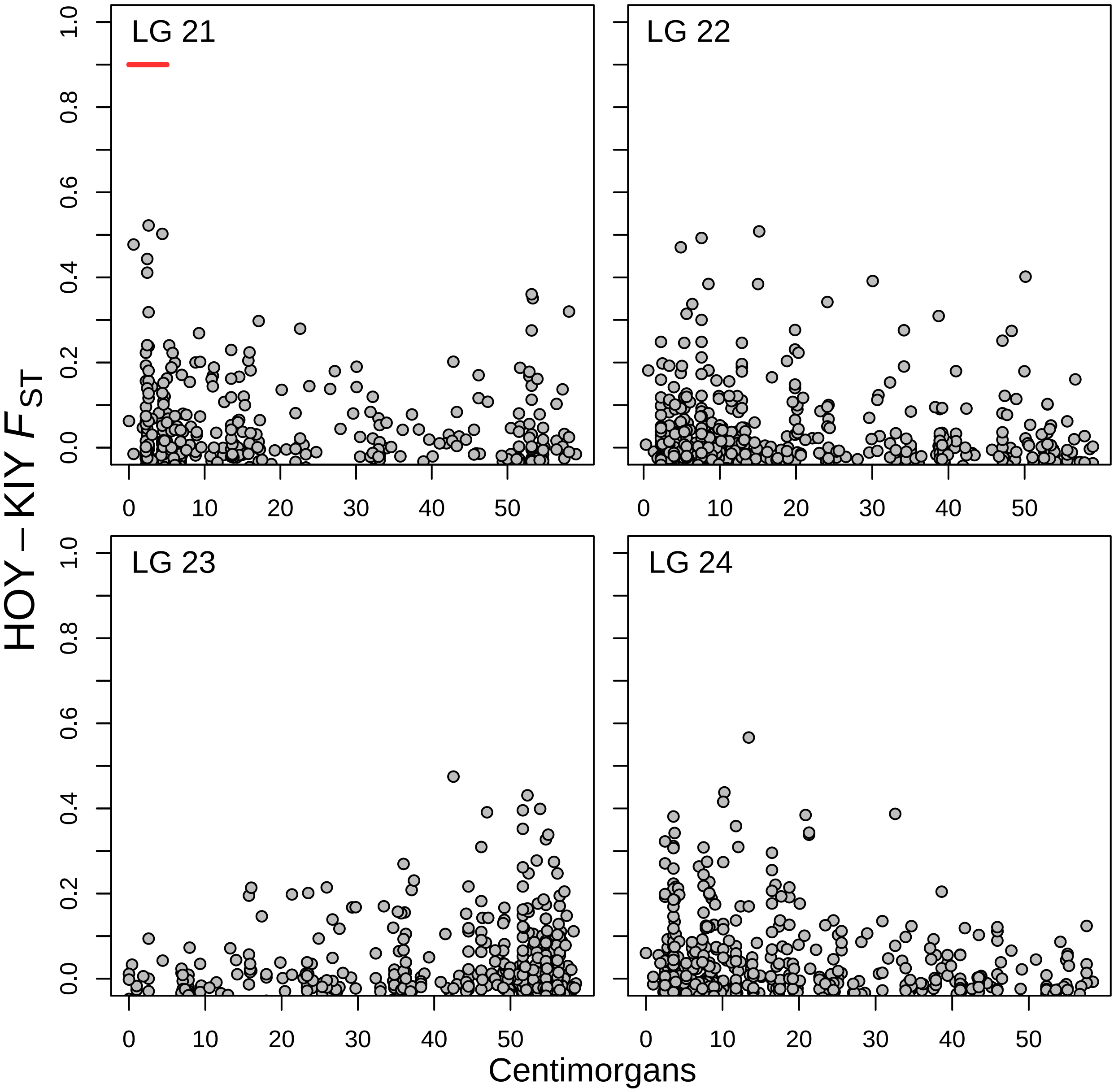

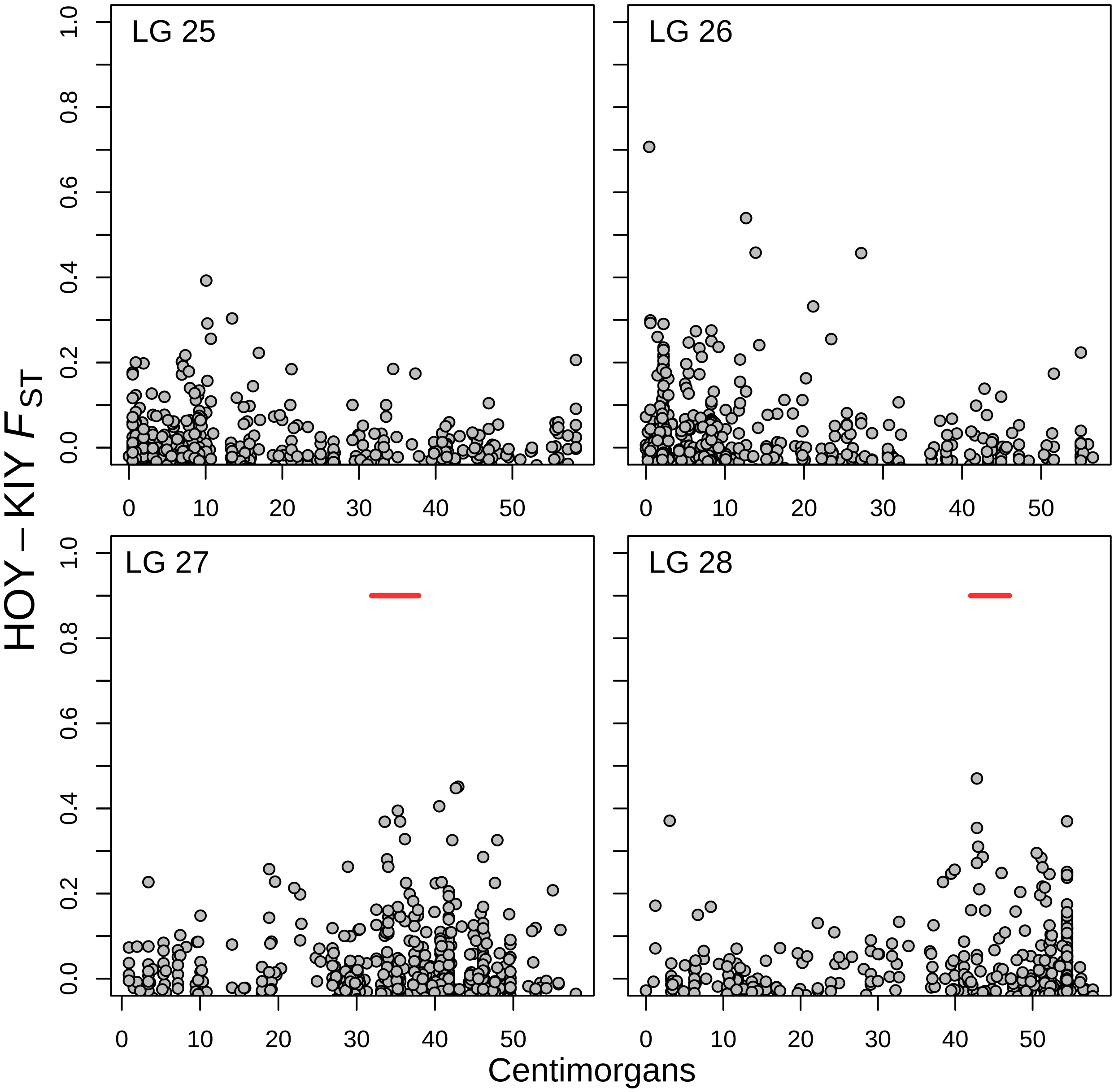

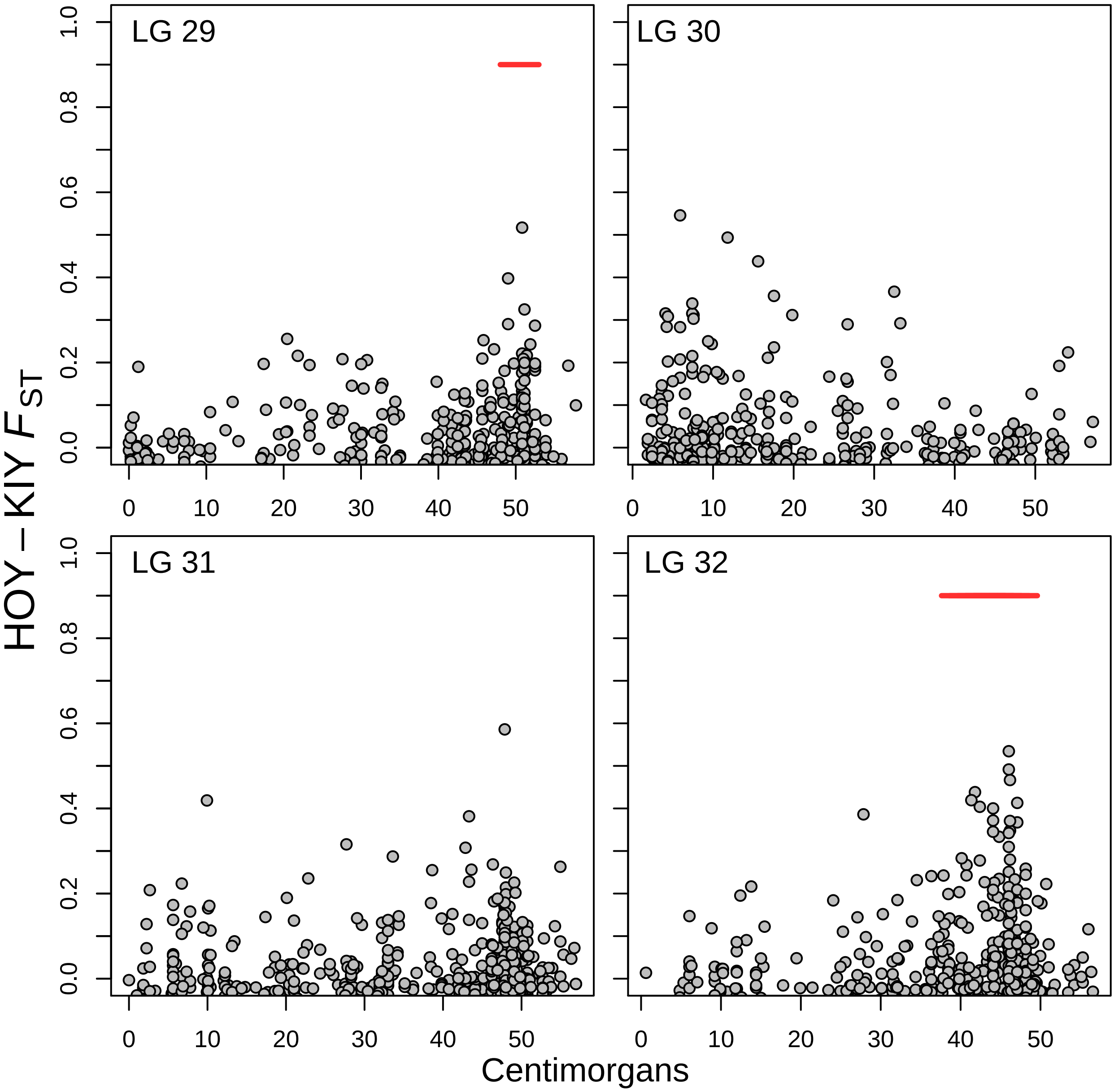

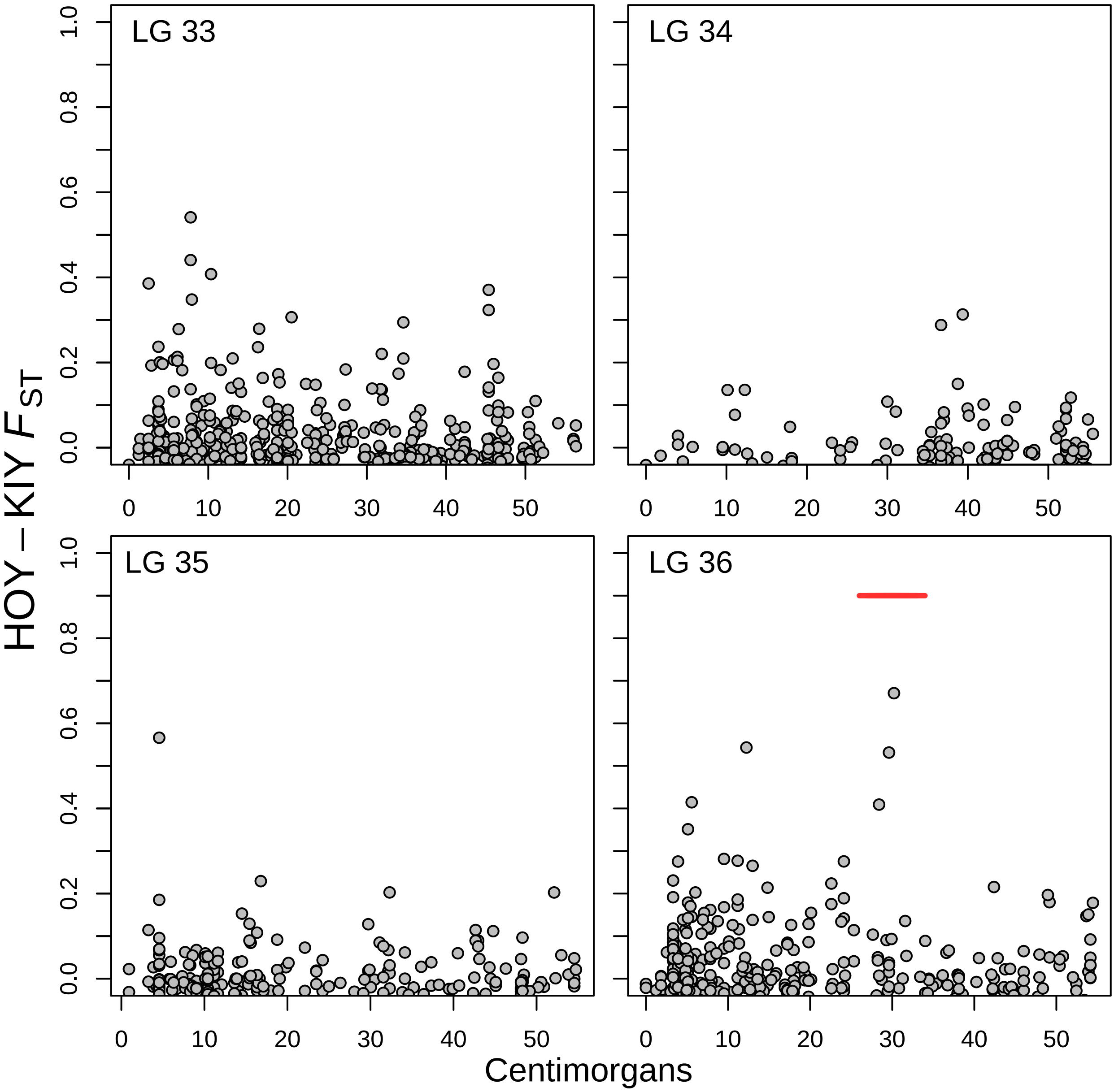

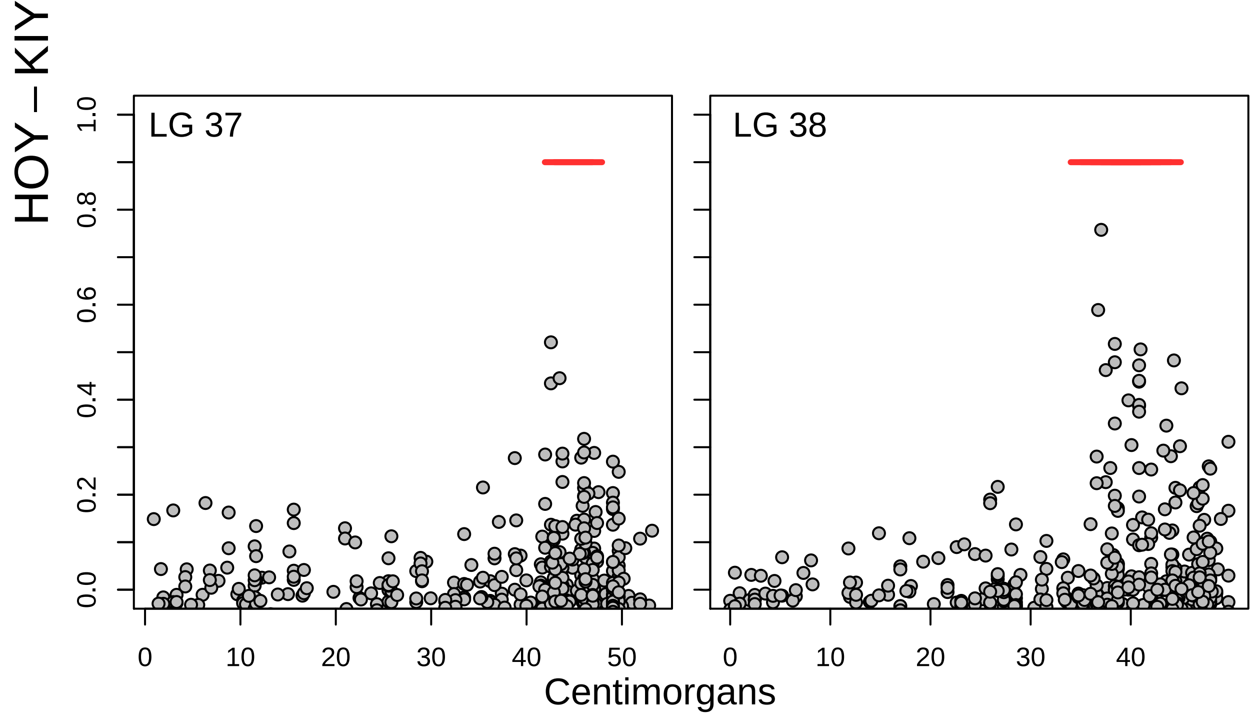

### Fig. S1

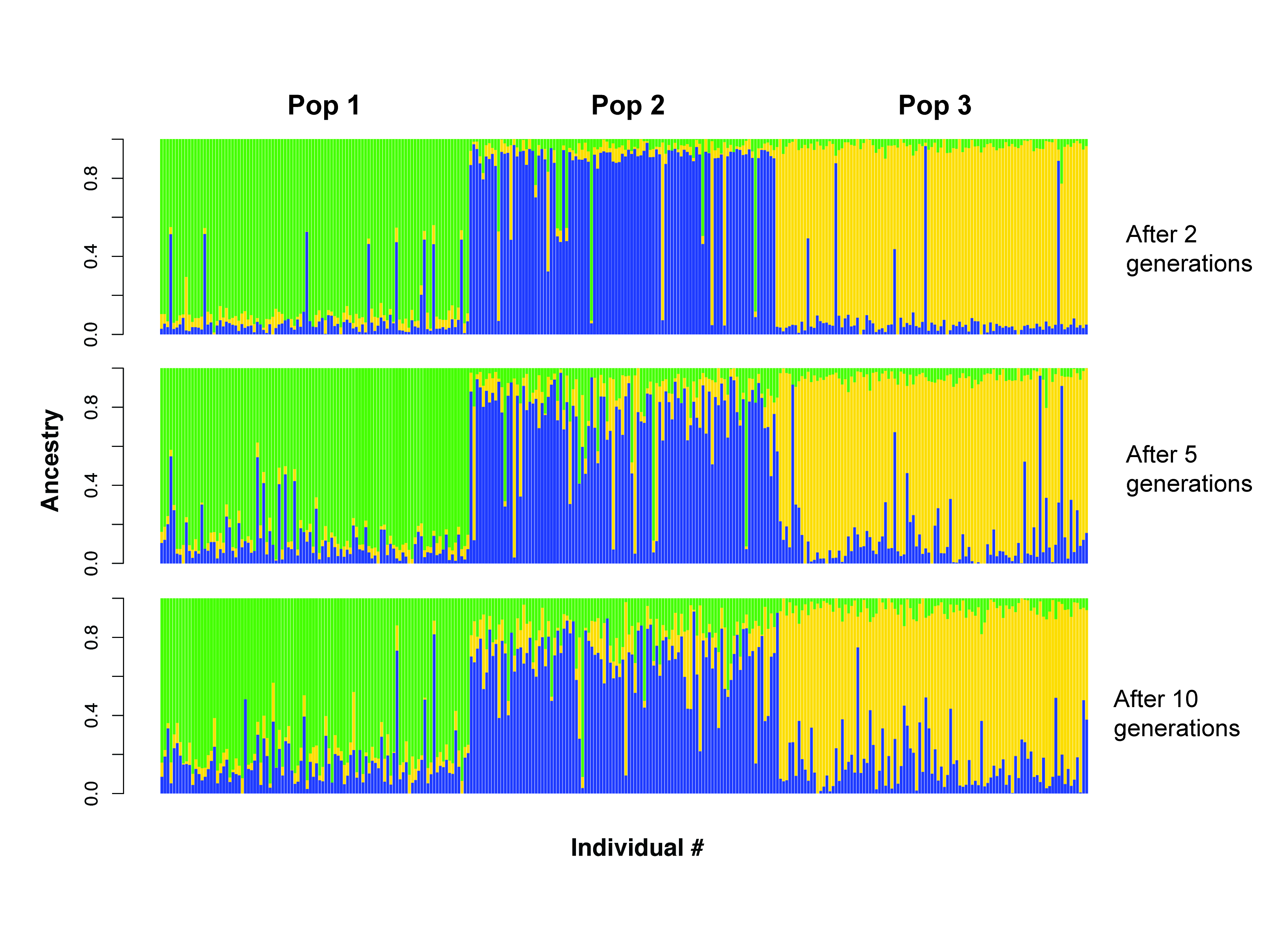

### Fig. S2

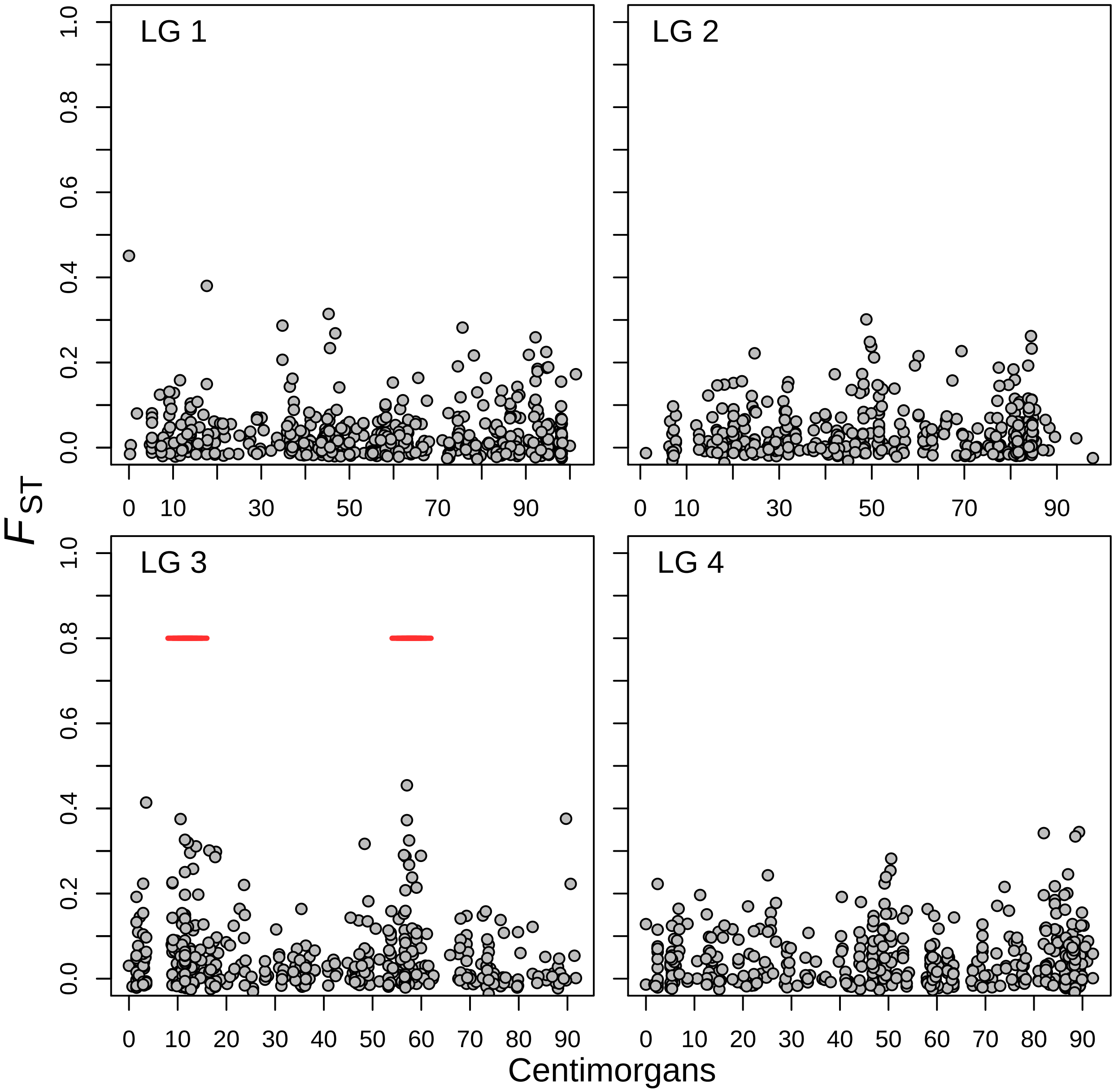

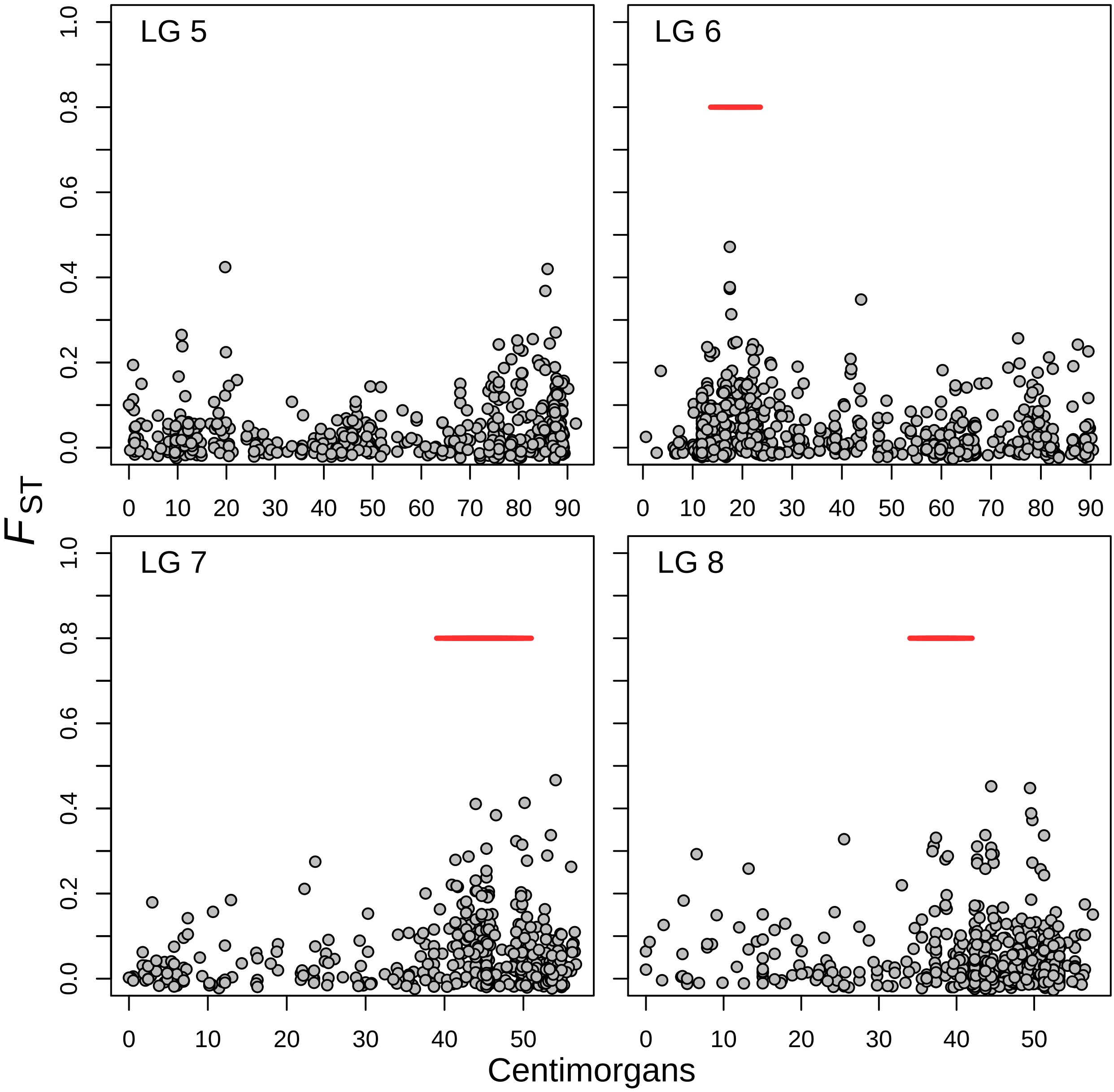

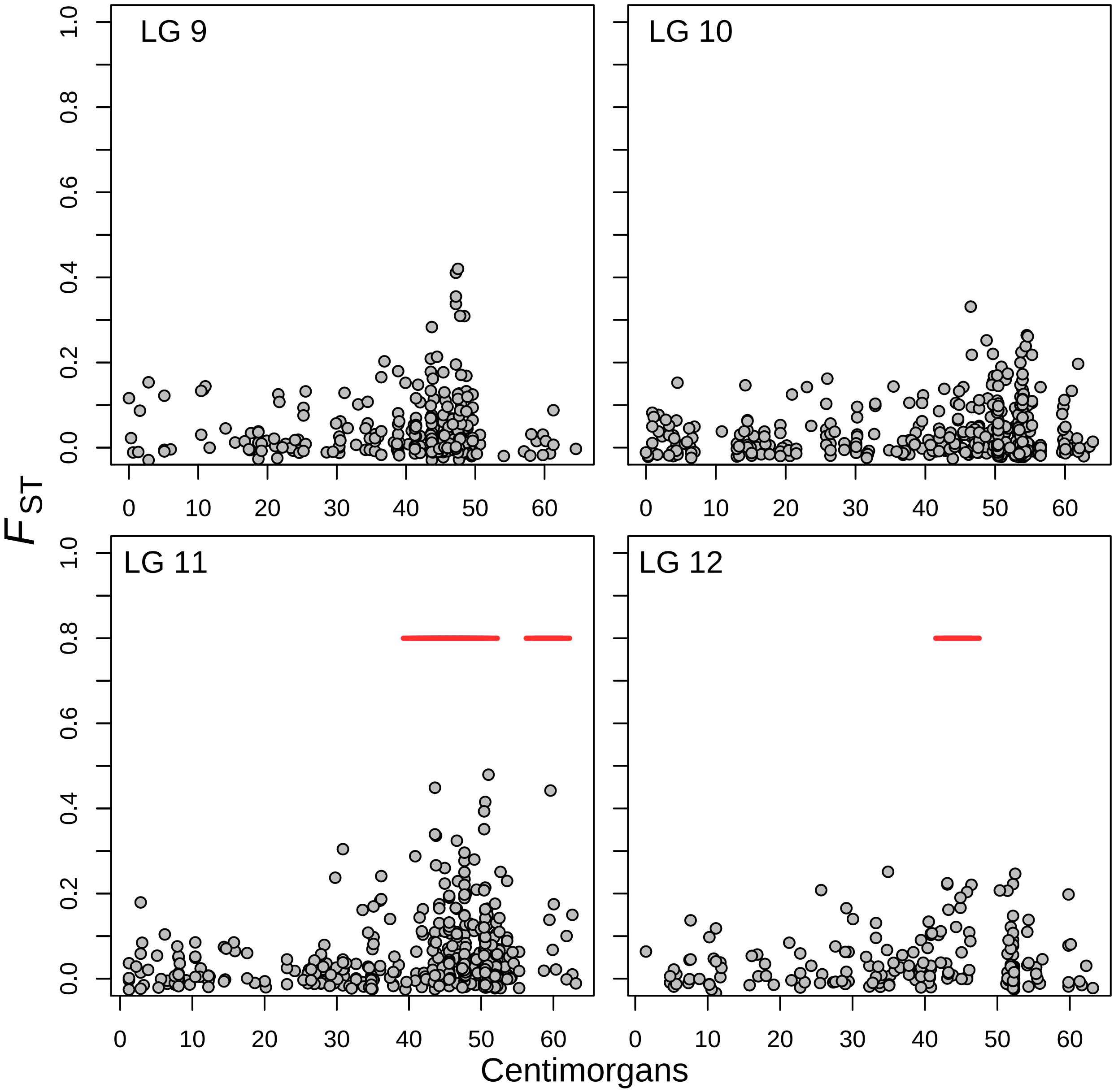

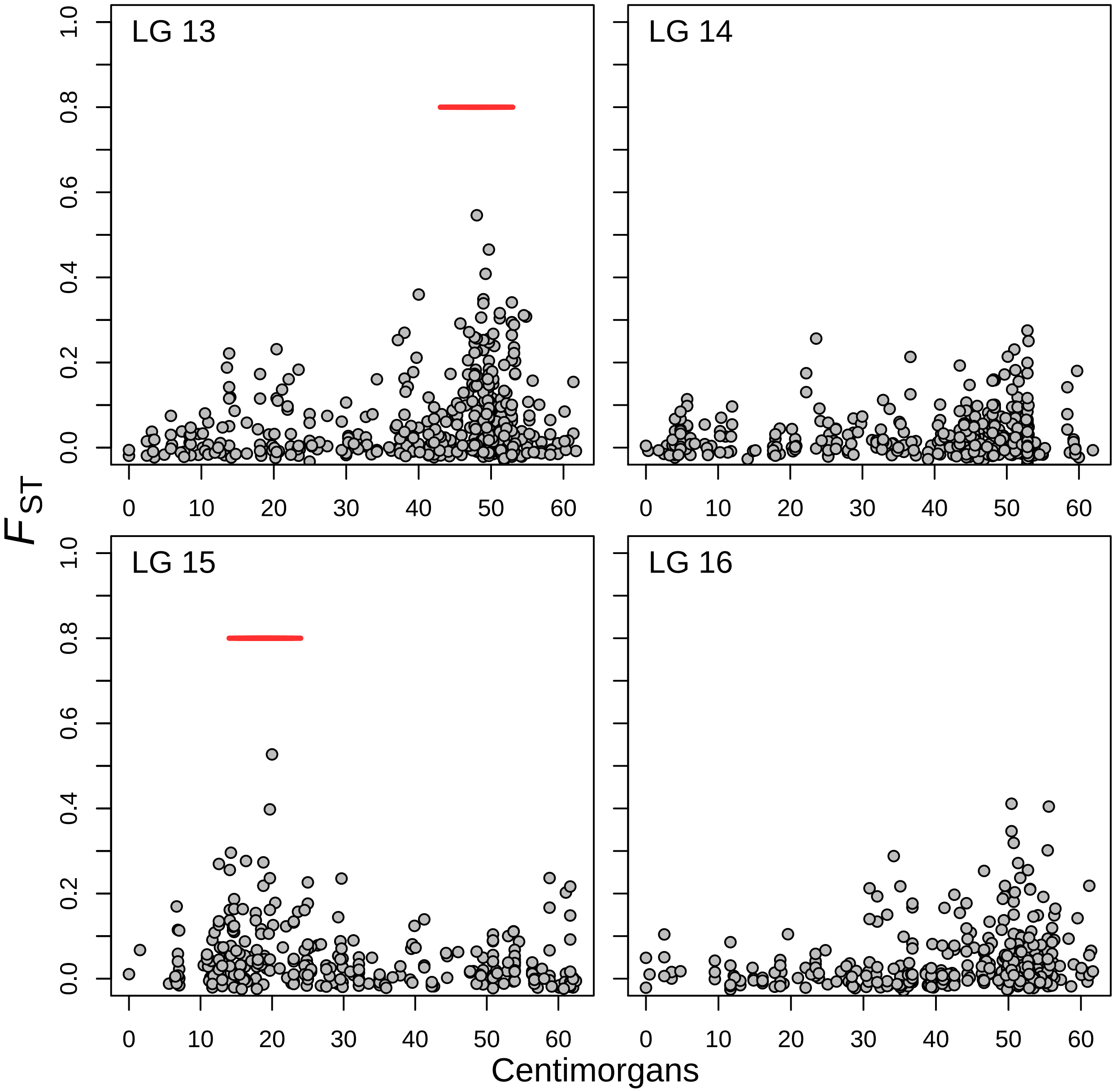

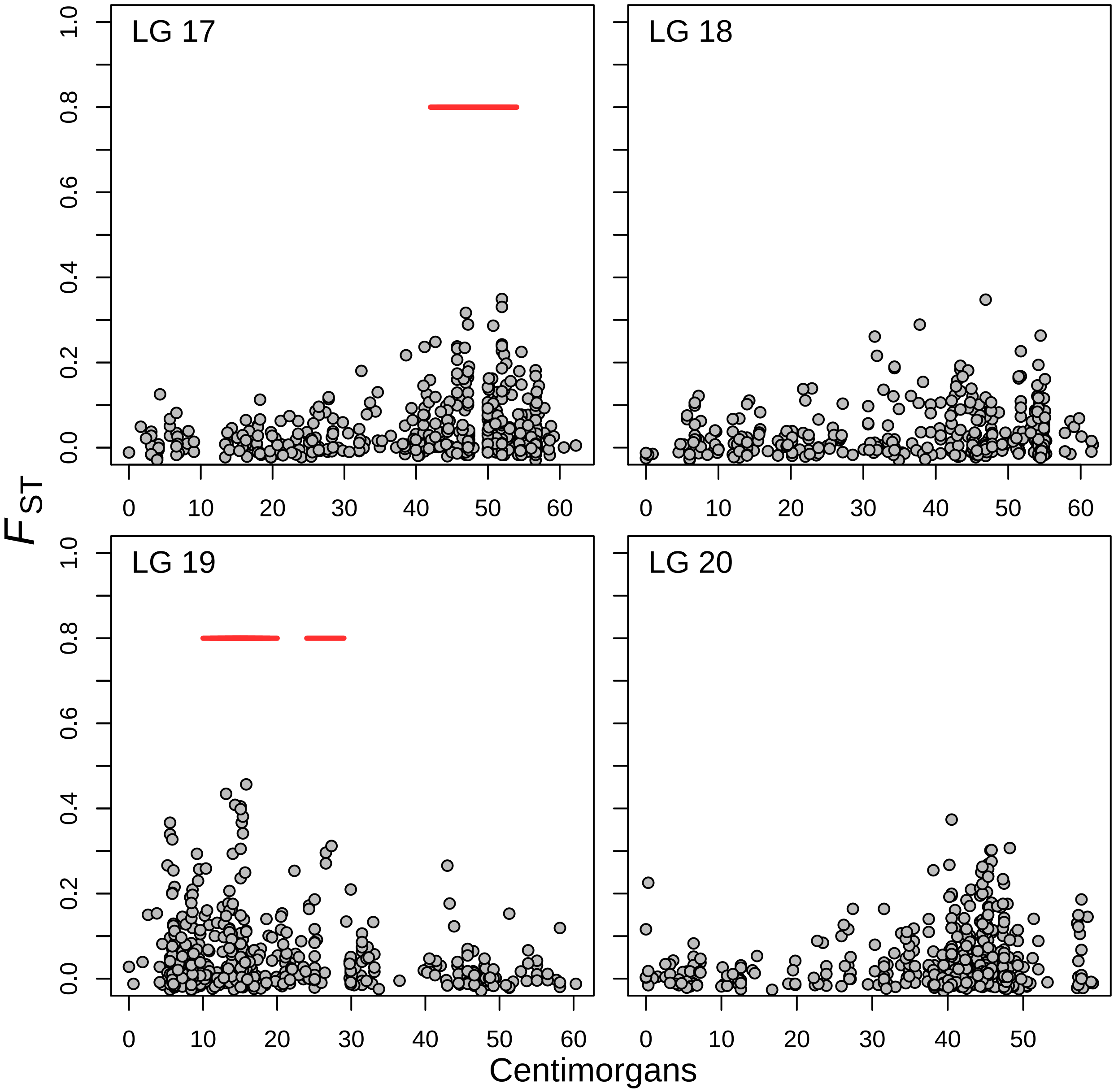

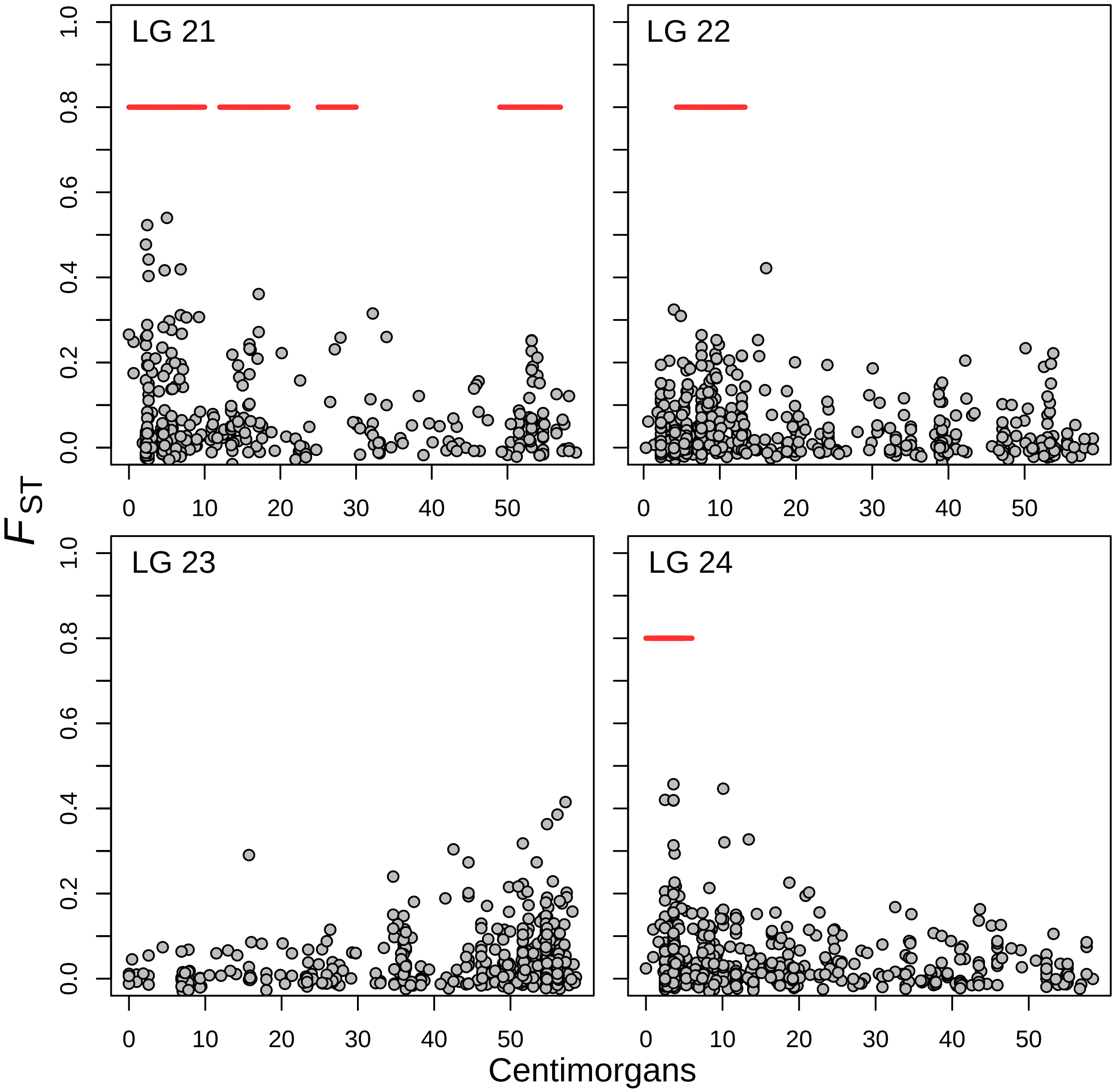

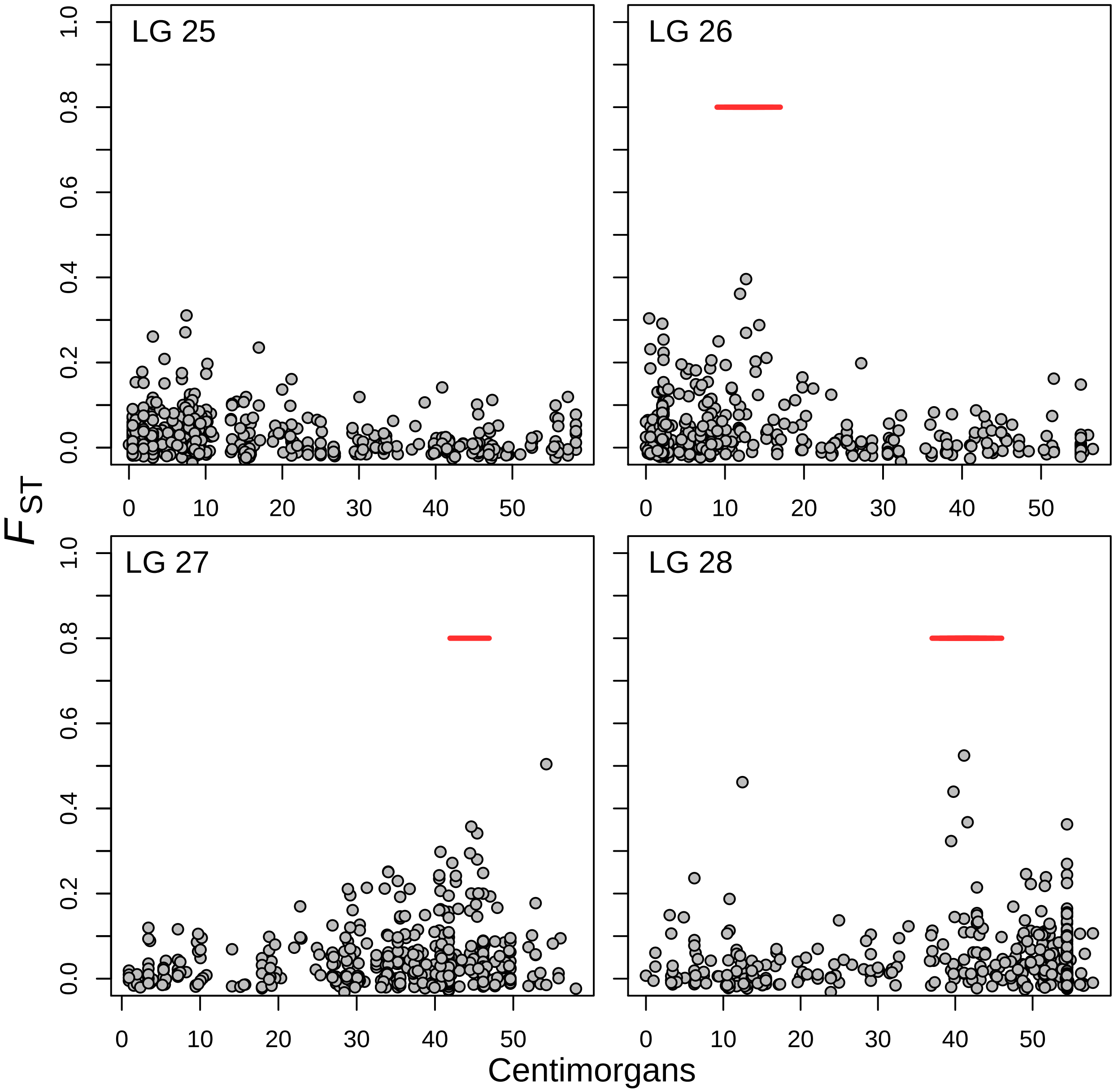

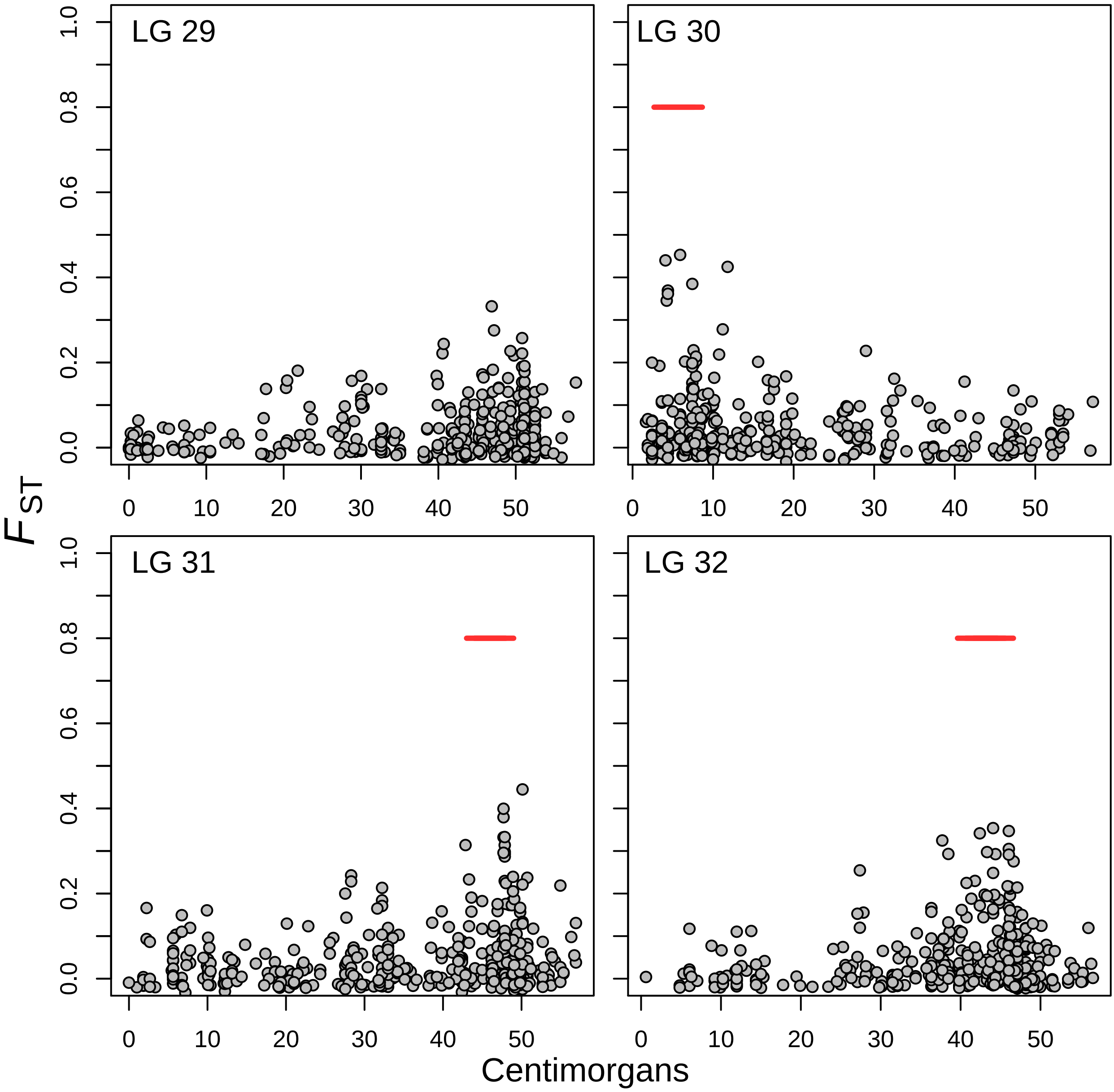

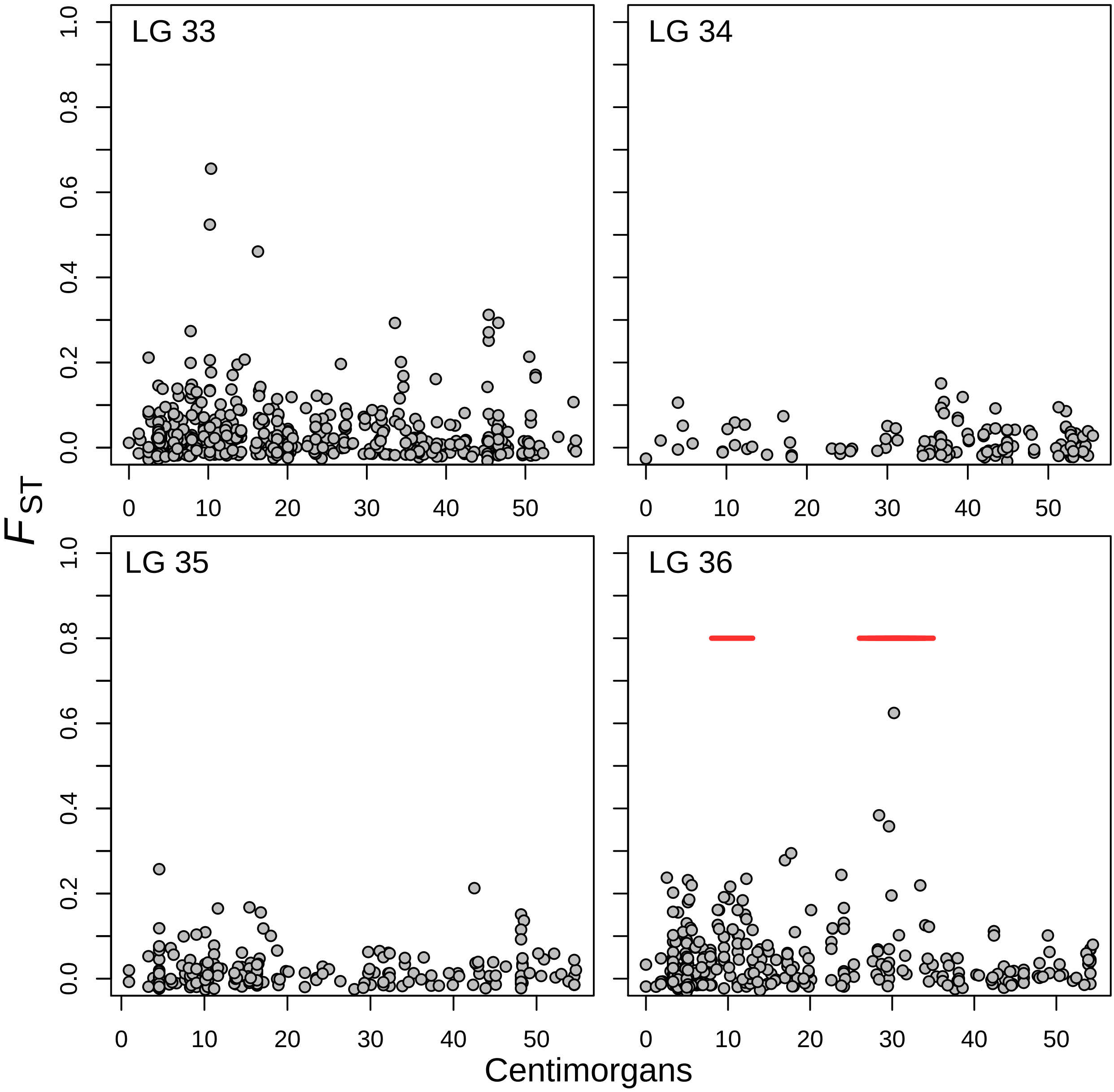

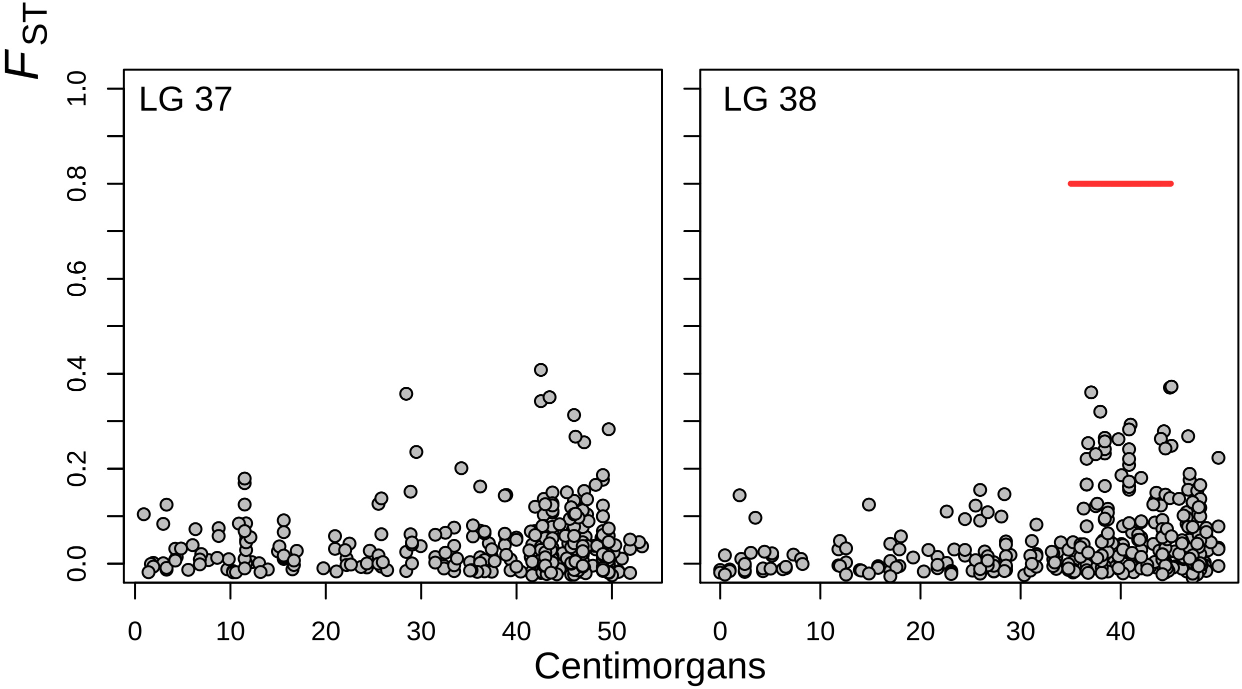
